# Identification of two novel lipoteichoic acid synthases essential for viability in *Clostridioides difficile*

**DOI:** 10.64898/2026.11.20.767628

**Authors:** Brianne R. Zbylicki, Elizabeth A. Warren, Hannah M. Laster, Kevin W. Bollinger, Ute Müh, Sonya Kumar Bharathkar, Beth M. Stadtmueller, David S. Weiss, Craig D. Ellermeier

**Author notes:** Present Address: Department of Biomedical Engineering, Translational Therapeutics and Regenerative Engineering Center, The Johns Hopkins University School of Medicine, Baltimore, Maryland, USA.

## Abstract

Lipoteichoic acids (LTAs) are major components of the Gram-positive bacterial cell envelope that contribute to cell morphology, division, and resistance to antimicrobial stress. *Clostridioides difficile* produces a type V LTA composed of repeating GlcNAc-GlcNAc-glyceric acid subunits anchored to the membrane by a glycolipid. Here, we identify two previously uncharacterized enzymes involved in *C. difficile* LTA synthesis: LtaB, which is required for LTA production and cell envelope integrity, and LtaA, which influences apparent LTA polymer length and partially compensates for reduced LtaB activity. Loss of LtaB causes growth and morphological defects, including aberrant septation and cell lysis, while LtaB depletion in the absence of LtaA is lethal. LtaA and LtaB contain two distinct domains: an LTA synthase-like domain and a glycosyltransferase (GT) C-like domain. Site-directed mutagenesis of conserved residues shows that the GT-C-like domain of LtaB is critical for both LTA synthesis and viability, whereas mutations in the LTA synthase-like domain have little effect on viability and only modest effects on LTA chain length. Production of several LtaB homologs from related species restored viability and, in some cases, LTA synthesis when a *C. difficile* Δ*ltaA* mutant was depleted of LtaB. These findings establish LtaA and LtaB as distinct but functionally overlapping enzymes required for type V LTA synthesis in *C. difficile*.

**Significance or Importance:** Lipoteichoic acids are important components of the cell envelopes of Gram-positive bacteria, yet the enzymes responsible for synthesizing structurally diverse LTAs remain poorly understood. *Clostridioides difficile*, a major cause of antibiotic-associated disease, produces a unique type V LTA, but the pathway responsible for its synthesis has remained unknown. Here, we identify two previously uncharacterized enzymes, LtaA and LtaB, that together are required for LTA production, cell envelope integrity, and viability in *C. difficile*. Our findings identify GT-C-like proteins involved in type V LTA biogenesis and reveal distinct but partially overlapping functions for LtaA and LtaB.

## Introduction

*Clostridioides difficile* is a Gram-positive, anaerobic, spore-forming pathogen and the most common cause of healthcare-associated diarrhea. In the United States, *C. difficile* is estimated to cause over 200,000 infections and 12,000 deaths annually, resulting in approximately one billion dollars in healthcare costs, and is classified by the Centers for Disease Control and Prevention as an urgent threat to public health (1).

The cell envelope of *C. difficile* consists of a proteinaceous Surface-layer (S-layer), a thick peptidoglycan layer, a cytoplasmic membrane, and several cell envelope-associated polysaccharides (2). *C. difficile* produces three distinct polysaccharides: PS-I, PS-II, and PS-III (3–5). PS-I consists of a repeating pentaglycosyl phosphate subunit (3) and is present in only a minority of *C. difficile* strains, whereas PS-II and PS-III are widely conserved (3, 4). PS-II is analogous to wall teichoic acid (WTA) and is composed of repeating hexaglycosyl phosphate units that are covalently linked to peptidoglycan. In *C. difficile*, PS-II anchors the major S-layer protein SlpA, as well as additional S-layer proteins, to the cell wall (3, 4, 6–8). PS-III is a lipoteichoic acid (LTA) composed of repeating disaccharides of N-acetylglucosamine-N-acetylglucosamine-glyceric acid (GlcNAc(GroA)-GlcNAc) linked by 6-*P*-6 phosphodiester bonds and anchored in the cytoplasmic membrane by the glycolipid triglucosyldiacylglycerol (TGDAG) (Fig. 1A,B) (4). Structural analyses indicate that PS-III is heterogeneous: approximately 30% of the repeats in phenol-soluble PS-III contain glucosamine (GlcN) in place of GlcNAc, and PS-III may also be modified by D-alanylation (4, 9–12).

**Figure 1.**
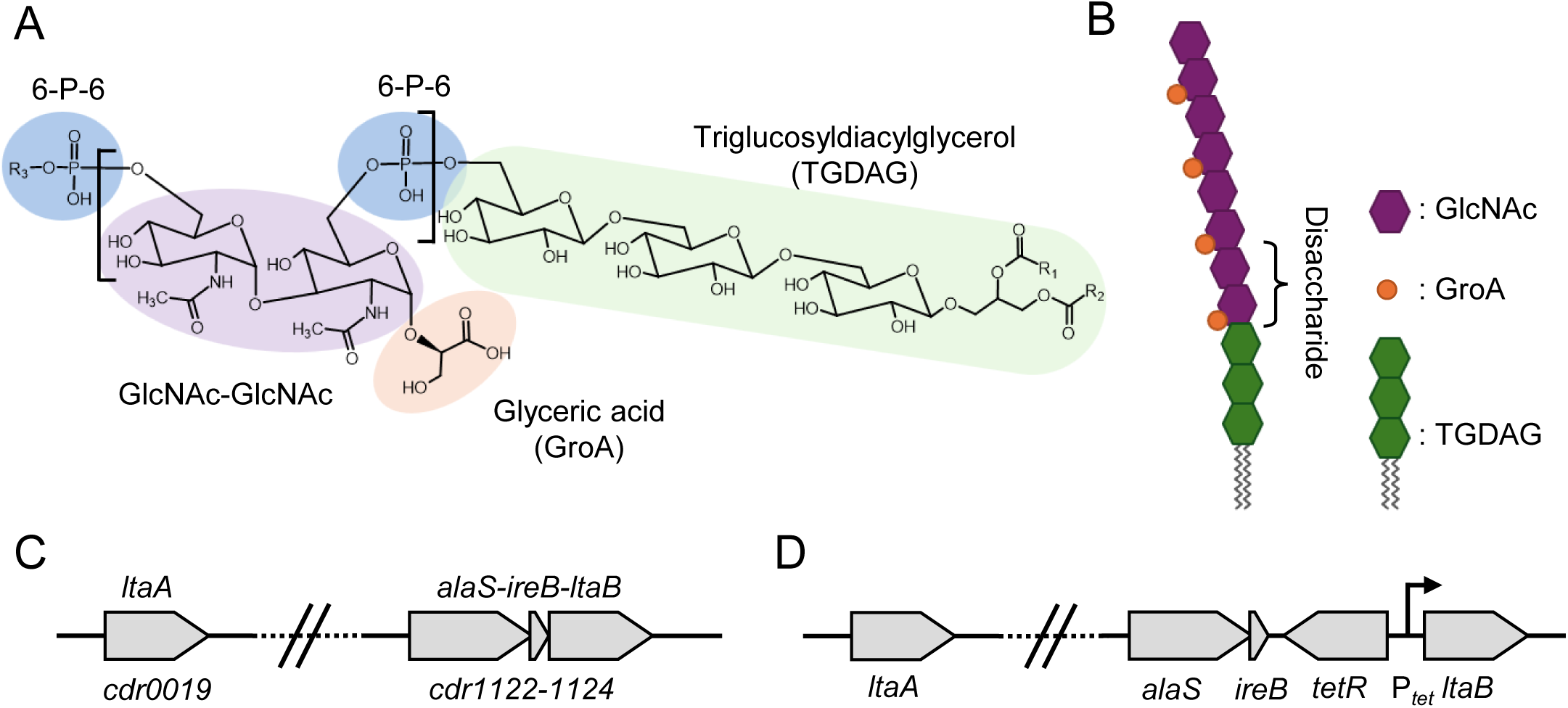
Structure of *Clostridioides difficile* lipoteichoic acid and genomic organization of *ltaA* and *ltaB*. **(A)** Chemical structure of the type V lipoteichoic acid (LTA) repeat and membrane anchor. The polymer contains repeating GlcNAc–GlcNAc–glyceric acid (GroA) units joined by 6-*P*-6 phosphodiester linkages and is anchored to the membrane by triglucosyldiacylglycerol (TGDAG). R_1_ and R_2_ denote lipid acyl chains and R_3_ denotes the extending chain of repeating GlcNAc(GroA)-GlcNAc subunits. **(B)** Schematic representation of *C. difficile* LTA. Purple hexagons represent GlcNAc, orange circles represent GroA, and green hexagons represent the TGDAG glycolipid anchor. The bracket indicates one disaccharide repeat unit. **(C)** Genomic organization of *ltaA* (*cdr0019*) and *ltaB* (*cdr1124*) in *C. difficile* strain R20291. *ltaB* is located immediately downstream of *alaS* and *ireB* within the *cdr1122–cdr1124* locus, whereas *ltaA* is encoded at a separate chromosomal locus. The interrupted line indicates the genomic separation between the loci. **(D)** LtaB depletion strain. *tetR* and P*_tet_* were inserted between *ireB* and *ltaB,* producing P*_tet_*-*ltaB*, in which *ltaB* expression depends on aTc.

LTAs are extracellular polymers of Gram-positive bacteria that are classified by the chemical composition of their repeating units and membrane anchors (13, 14). Type I LTA, found in organisms such as *Staphylococcus aureus, Bacillus subtilis*, and *Listeria monocytogenes*, consists of glycerol phosphate (GroP) units linked to a glycolipid anchor (13, 14). *S. aureus* uses LtaS to synthesize LTA by transferring GroP from phosphatidylglycerol to the growing polymer (15–18). *L. monocytogenes* employs two synthases, LtaP and LtaS, for polymer initiation and elongation (19, 20). *B. subtilis* encodes four synthases (LtaS, YvgJ, YfnI, and YqgS), which contribute differentially to LTA synthesis under distinct physiological conditions (18, 21–24).

In contrast to the relatively simple GroP polymer of type I LTA, types II-V contain more complex glycan repeats and rely on less well-defined biosynthetic pathways (5, 25). Type II and III LTAs in *Lactococcus garvieae* and *Clostridium innocuum* contain galactose repeats attached via glycolipids, although their biosynthetic genes remain unidentified (13, 14, 26). *Streptococcus pneumoniae* produces a type IV LTA with the same repeating structure as its WTA (27–30); a shared lipid precursor is transferred to peptidoglycan or a glycolipid to form WTA or LTA, respectively (27–30). Type V LTA encompasses lipoglycans of Gram-positive bacteria, including *C. difficile* and *P. anaerobius*, with GlcNAc(GroA)-GlcNAc repeats attached to glycolipid anchors (4, 14, 31, 32). This structural diversity, together with the limited knowledge of the corresponding biosynthetic enzymes, underscores why the type I LTA synthases provide an incomplete model for type V LTA synthesis in *C. difficile*.

Although the precise physiological functions of LTA remain incompletely understood, loss of LTA or disruption of LTA synthesis causes severe growth and morphological defects in many organisms. In *S. aureus* and *L. monocytogenes*, loss of LtaS severely impairs growth at physiological temperatures and results in defective septation and cell separation (18, 20, 33). In *B. subtilis*, a mutant lacking all four LTA synthases is viable but has pronounced cell division and morphological defects. Simultaneous loss of both LTA and WTA is lethal, suggesting functional overlap between these polymers (21, 24). These studies support an important role for LTA in maintaining cell envelope integrity, particularly during cell division.

Despite LTA’s importance for cell envelope function, the proteins responsible for LTA synthesis in *C. difficile* have not been identified. We report two previously uncharacterized proteins, LtaA and LtaB, that are needed for LTA synthesis. LtaA and LtaB share structural similarity with known LTA synthases and glycosyltransferase C superfamily members. Loss of LtaA alone does not detectably affect growth or morphology but alters apparent LTA polymer length, whereas loss of LtaB eliminates detectable LTA and causes severe growth and shape defects. Depleting LtaB in the absence of LtaA is lethal and results in production of abnormally short LTA species. Together, these findings identify LtaA and LtaB as components of the *C. difficile* LTA biosynthetic pathway and provide new insight into type V LTA synthesis in Gram-positive bacteria.

## Results

### Bioinformatic Identification of Two Putative LTA Synthases

We hypothesized that *C. difficile* LTA synthesis enzymes might resemble those in other organisms, despite the unique LTA structure. BLASTP search of strain R20291’s genome, ribotype 027, for homologs of *B. subtilis* or *S. aureus* LtaS yielded no significant hits, implying that *C. difficile* uses evolutionarily different enzymes for LTA synthesis (34).

To identify more distantly related candidates, we turned to HHpred, which uses hidden Markov models to detect remote structural homology (35). This analysis identified two largely uncharacterized *C. difficile* proteins, CDR20291_0019 and CDR20291_1124, with predicted structural similarity to LtaS from *B. subtilis*. Both proteins had highly significant HHpred scores (E values <4.7 x 10^-15^), consistent with meaningful structural similarity. Based on the functional data described below, we named these proteins LtaA (CDR20291_0019) and LtaB (CDR20291_1124). Pairwise alignment revealed that LtaA and LtaB are closely related, sharing 63% amino acid identity and 82% similarity (Fig. S1).

Topology prediction using SignalP and DeepTMHMM indicated that both LtaA and LtaB contain an N-terminal cleavable signal peptide followed by a large extracellular domain and a C-terminal membrane-embedded region containing eleven predicted transmembrane helices (Fig. S2) (36, 37). BLASTP searches of public databases failed to identify homologs with annotated function; however, HHpred analysis revealed that the N-terminal extracellular domains of LtaA and LtaB share structural similarity with type I LTA synthases, putative sulfatases, and glycosylphosphatidylinositol (GPI) ethanolamine phosphate transferases, which generate lipid anchors for cell-surface proteins in eukaryotes (Table S1) (38). The C-terminal transmembrane domains of LtaA and LtaB showed similarity to multiple members of the glycosyltransferase C (GT-C) superfamily (Table S1). GT-C enzymes are integral membrane proteins characterized by multiple transmembrane helices and catalyze the synthesis or transfer of complex polysaccharides using lipid-linked substrates (39). Structural homologs included WaaL, the O-antigen ligase responsible for attaching O-antigen to lipid A in Gram-negative bacteria, and MrdB, which transfers peptidoglycan precursors from a lipid carrier to the growing cell wall (40, 41). Because *C. difficile* LTA consists of repeating disaccharide subunits and LtaA/LtaB resemble GT-C enzymes, these proteins may participate in the transfer or assembly of lipid-linked LTA precursors.

### Loss of Both LtaA and LtaB is Lethal

Transposon sequencing (TN-seq) data predict that *ltaA* is nonessential and *ltaB* is essential in *C. difficile* (42, 43). We previously showed CRISPR interference (CRISPRi) of *ltaB* resulted in decreased viability and morphological defects (43). To assess the roles of these genes experimentally, we first constructed an *ltaA* deletion using CRISPR-Cas9. Strains in Fig. 2A, C-G carry the denoted plasmids for complementation experiments; phenotypes of plasmid-free deletion strains are shown in Fig. 2B and Fig. S3. Growth of the Δ*ltaA* mutant was indistinguishable from wild type on both solid medium and in liquid culture (Fig. 2B; Fig. S3A,C) and there were no detectable defects in cell length, sinuosity, or septa per cell (Fig. 2C-G).

**Figure 2.**
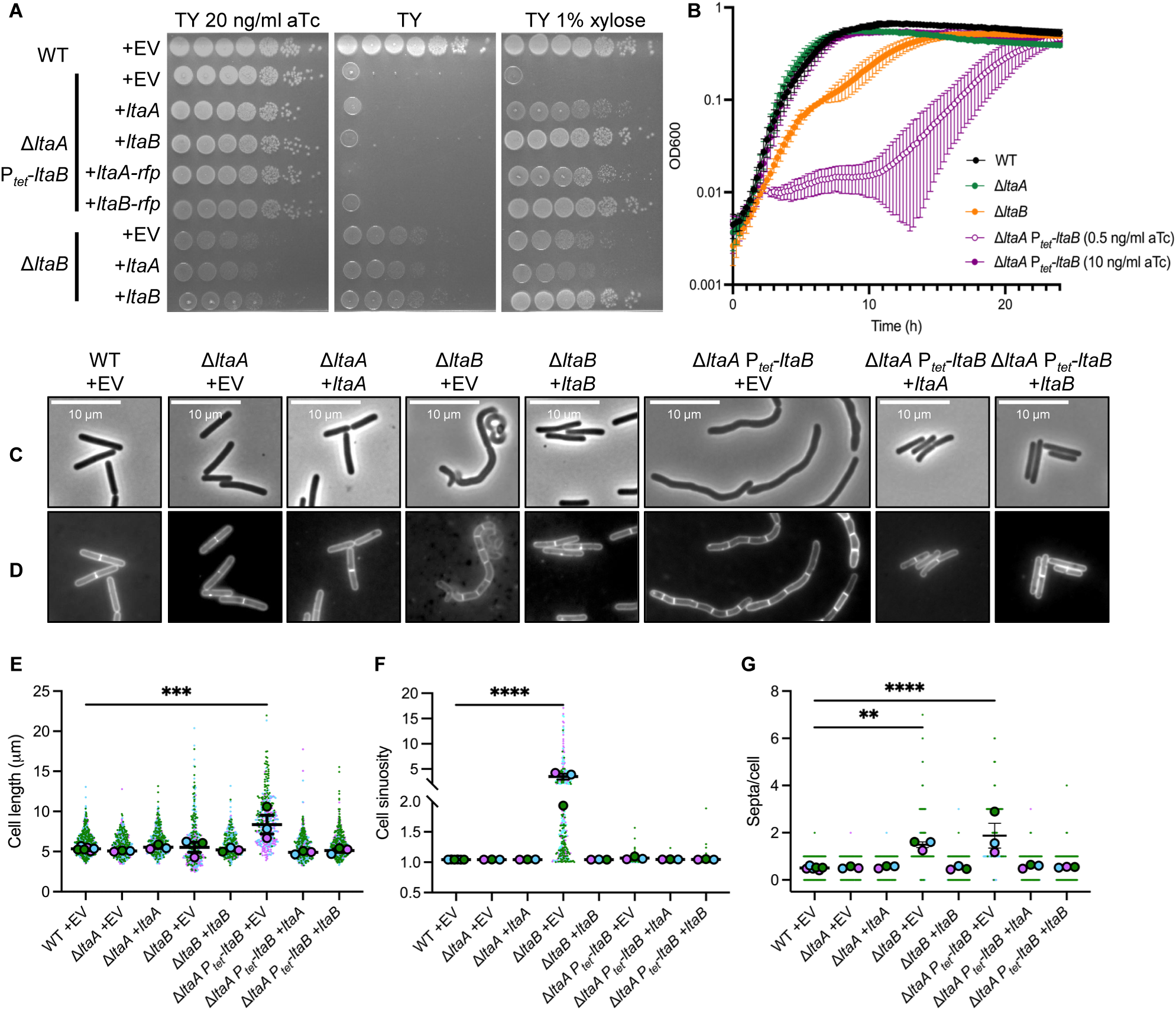
LtaA and LtaB are required for growth, morphology, and viability in *C. difficile*. Loss or depletion of *ltaB* causes severe defects in growth, cell morphology, and division, whereas loss of *ltaA* alone has no detectable effect. **(A)** Spot-titer viability assays comparing wild-type and mutant strains grown under inducing (with Anhydrotetracycline (aTc)) or noninducing (without aTc) conditions for *ltaB* expression, or with complementation from a xylose-inducible promoter where indicated. **(B)** Growth curves in TY broth measured as OD_600_ over time. Points and error bars represent the mean ± SD from four biological replicates. **(C)** Phase-contrast and **(D)** membrane fluorescence micrographs. Strains were grown overnight in TY with aTc (5 ng/mL for Δ*ltaA* P_tet_-*ltaB*; 1 ng/mL for P_tet_-*ltaB*) and subcultured 1:50 into TY with aTc for depletion conditions (1 ng/mL for Δ*ltaA* P_tet_-*ltaB*; 0.5 ng/mL for P_tet_-*ltaB*) or inducing conditions (10 ng/mL for both strains). Cells were sampled at OD_600_ ≈ 0.5 and stained with the membrane dye FM4-64. Scale bars, 10 µm. Images are representative of three independent experiments. **(E)** Quantification of average cell length based on three biological replicates, with at least 300 cells measured per condition. Filled circles depict the mean value from each replicate; bars and error bars represent the mean ± SD across replicates. **(F)** Cell sinuosity, defined as the ratio of medial axis length to pole-to-pole distance. **(G)** Average number of septa per cell, quantified from FM4-64–stained cells. Statistical significance in (E–G) was assessed by one-way ANOVA with Dunnett’s multiple-comparison test. ns, not significant; **p < 0.01; ***p < 0.001; ****p < 0.0001. **Strains used: (A & C-G)** WT + EV = BZ153; Δ*ltaB* EV = EW211; Δ*ltaB* + *ltaA* = EW213; Δ*ltaB* + *ltaB* = EW223; Δ*ltaA* P*_tet_-ltaB* + EV = CDE5031; Δ*ltaA P_tet_-ltaB + ltaA* = CDE5033; Δ*ltaA P_tet_-ltaB + ltaB* = CDE5032; Δ*ltaA P_tet_-ltaB + ltaA-rfp* = CDE5233; Δ*ltaA P_tet_-ltaB + ltaB-rfp* = CDE5231; Δ*ltaA* + EV = CDE5048; Δ*ltaA* + *ltaA* = CDE5050. **(Β)** WT = R20291; Δ*ltaA* = CDE5015; Δ*ltaB* = EW169; P*_tet_-ltaB* = CDE5011; Δ*ltaA P_tet_-ltaB* = CDE5029

*ltaB* is encoded as the terminal gene in an operon containing *alaS* and *ireB*, which encode an essential alanyl-tRNA synthetase and a regulatory phosphoprotein, respectively (Fig. 1C). Although *ltaB* is predicted to be essential, we generated a Δ*ltaB* mutant using CRISPR-Cas9. However, the deletion mutant exhibited a severe growth defect, likely explaining its classification as essential by Tn-Seq (42, 43) (Fig. 2A,B; Fig. S3A). In addition, Δ*ltaB* cells exhibited severe morphological abnormalities, including increased cell curvature, chaining, and aberrant septation (Fig. 2C-G). Expression of *ltaB in trans* restored both growth and normal cell morphology, confirming that these phenotypes were specifically attributable to loss of LtaB (Fig. 2A,C-G; Fig. S3D).

Given the sequence similarity between LtaA and LtaB, we asked if these proteins might have partially redundant functions. We could not construct a Δ*ltaA* Δ*ltaB* double mutant, suggesting that simultaneous loss of both genes is lethal. To test this, we engineered a depletion strain where *tetR* and a tetracycline-repressible promoter (*P*_tet_) were inserted between the end of *ireB* and the start of *ltaB*, generating a Δ*ltaA P*_tet_-*ltaB* strain (Fig. 1D). This strain grew comparably to wild type in the presence of inducing concentrations of anhydrotetracycline (aTc) but exhibited a severe growth defect under non-inducing or minimally inducing conditions (Fig. 2A,B; Fig. S3A). In contrast, a *P*_tet_-*ltaB* strain had a modest growth defect under non-inducing conditions, consistent with low-level leaky expression from the *P*_tet_ promoter (Fig. 2A; Fig. S3A,B).

To further test the functional overlap between LtaA and LtaB, we introduced plasmids expressing either *ltaA* or *ltaB* under the control of a xylose-inducible promoter (*P*_xyl_) into the Δ*ltaA P*_tet_-*ltaB* depletion strain. Expression of *ltaB* fully restored wild-type growth and colony morphology, while expression of *ltaA* partially rescued growth and morphology, yielding cells and colonies similar to those produced by the Δ*ltaB* mutant (Fig. 2A,C-G; Fig. S3E). In contrast, expression of *ltaA* in the Δ*ltaB* mutant failed to restore viability or normal morphology (Fig. 2A,C-G), indicating that LtaA alone cannot substitute for LtaB.

We used time-lapse microscopy to characterize the terminal phenotype of LtaB depletion in the Δ*ltaA* background. We observed that loss of LtaA combined with depletion of LtaB caused cells to lose rod shape, become curved, bulge, and ultimately lyse (Movies S1 and S2). These results indicate that simultaneous loss of LtaA and LtaB is lethal. Although LtaA can partially compensate for reduced LtaB activity, LtaB performs the primary role required for normal cell growth, shape, and survival.

### LtaB is Essential for LTA Synthesis

To determine if LtaA and LtaB are required for LTA synthesis in *Clostridioides difficile*, we examined LTA production in deletion and depletion strains using two complementary approaches: immunoblotting with an anti-*C. difficile* LTA antibody and metabolic labeling using a fatty acid analog with an alkyne modification that allows copper-catalyzed cycloaddition (“click chemistry”) with a fluorescently labeled azide (44).

We first analyzed LTA production by immunoblotting following partial purification of LTA from cell extracts (4, 45). A robust LTA signal was detected in wild-type cells and in the Δ*ltaA* mutant, whereas LTA was undetectable in the Δ*ltaB* mutant (Fig. 3A). Serial dilution of the wild-type LTA extract showed that the immunoblot could detect a 10-to 25-fold reduction in signal, indicating that LTA in the Δ*ltaB* mutant was less than approximately 4–10% of the wild-type level under these conditions (Fig. S4A). When the Δ*ltaA P*_tet_-*ltaB* strain was grown under depletion conditions, no LTA was detected, while induction of *ltaB* expression restored LTA production (Fig. 3A). LTA produced by both Δ*ltaA* and Δ*ltaA P*_tet_-*ltaB* strains under inducing conditions appeared modestly shorter than LTA from wild-type, suggesting that LtaA influences the length distribution of LTA polymers. In contrast, the *P*_tet_-*ltaB* strain grown under inducing conditions produced LTA that was consistently longer and more abundant than LTA from wild-type (Fig. 3A). Because LtaB abundance was not measured directly, we cannot determine whether this altered LTA profile reflects differences in LtaB expression or activity.

**Figure 3.**
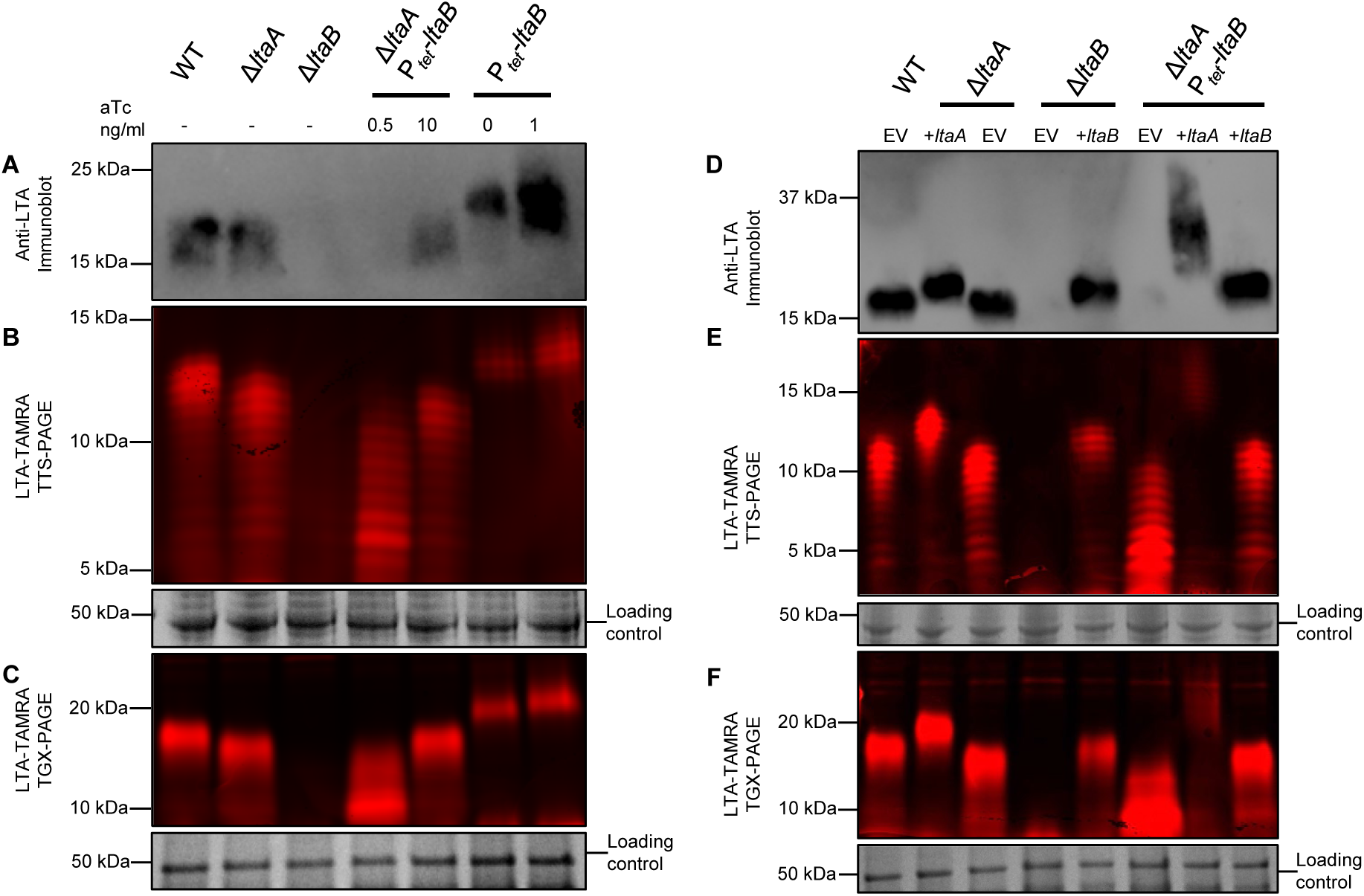
LtaB is required for lipoteichoic acid synthesis, whereas LtaA influences polymer length. Loss of *ltaB* eliminates detectable lipoteichoic acid. Loss of LtaA alters the apparent LTA polymer-length distribution without altering the apparent LTA concentration. **(A)** Immunoblot detection of LTA extracted from wild-type and mutant strains grown under inducing or depleting conditions for *ltaB* expression. LTA was detected using a recombinant human IgA anti-LTA antibody. **(B, C)** Detection of LTA metabolic labeling and in-gel fluorescence. Extracts were separated on Tris-Tricine (TTS-PAGE) or Tris-Glycine (TGX-PAGE) gels. Note that Tris-Tricine gels resolve low-molecular-weight LTA species. After fluorescence scanning, gels were stained with Coomassie blue to visualize the major S-layer protein as a loading control. **(D)** Immunoblot detection of LTA produced by Δ*ltaB* and Δ*ltaA Ptet-ltaB* strains expressing *ltaA* or *ltaB* from a xylose-inducible promoter. **(E,F)** Metabolic labeling and in-gel fluorescence of LTA from the same strains expressing xylose-induced *ltaA* or *ltaB*. Note that expression of *ltaB* restores LTA abundance and polymer length, whereas *ltaA* expression partially restores polymer length but not wild-type profiles. All images are representative of at least three independent experiments. Δ*ltaA Ptet-ltaB* strains in (D–F) were grown under depletion conditions. **Strains used: (A-C)** WT = R20291; Δ*ltaA* = CDE5015; Δ*ltaB* = EW169; P*_tet_-ltaB* = CDE5011; Δ*ltaA* P*_tet_-ltaB* = CDE5029; **(D-F)** WT + EV = BZ153; Δ*ltaA* + EV = CDE5048; Δ*ltaA* + *ltaA* = CDE5050; Δ*ltaB* + EV = EW211; Δ*ltaB* + *ltaB* = EW223; Δ*ltaB* + *ltaA* = EW213; Δ*ltaA* P*_tet_-ltaB* + EV = CDE5031; Δ*ltaA* P*_tet_-ltaB* + *ltaA* = CDE5033; Δ*ltaA* P*_tet_-ltaB* + *ltaB* = CDE5032.

Cairns et al. previously demonstrated that recognition of *C. difficile* LTA by monoclonal antibodies, including the 4D6 antibody used here, requires at least three phosphorylated, glycerated disaccharide repeat units in the context of the core and lipid anchor (46). Therefore, we considered that reduced immunoreactivity might reflect altered LTA structure or polymer length rather than a complete loss of LTA. To address this concern, we assayed lipid-linked LTA using a modified metabolic labeling technique previously used to study lipoproteins in *C. difficile* (44). Briefly, cells were grown in the presence of myristic acid alkyne, which is incorporated into lipids, collected by centrifugation, resuspended in buffer and lysed by sonication. Lysates were treated with a red-fluorescent TAMRA azide dye under conditions that promote cycloaddition to the alkyne by click chemistry” (44). After derivatization, labeled polymers were separated by SDS-PAGE and visualized by in-gel fluorescence (47).

When labeled extracts were separated on Tris-Glycine gels, LTA appeared as a broad red-fluorescent band migrating between the 15 and 20 kDa molecular weight markers of a protein standard (Fig. 3C). LTA from the Δ*ltaA* mutant was similar in size and abundance, whereas no LTA was detected in extracts of the Δ*ltaB* mutant (Fig. 3C). The sensitivity of LTA detection by metabolic labeling was similar to that of immunoblotting as indicated by serial dilutions (Fig. S4B). Importantly, the fluorescent bands were not sensitive to treatment with Proteinase K, consistent with their assignment as LTA rather than lipoproteins (Fig. S4C).

In contrast to the immunoblotting results, metabolic labeling detected the presence of a short LTA species in the Δ*ltaA* P*_tet_*-*ltaB* strain grown under depletion conditions (0.5 ng/ml aTc in Fig. 3A,C). Separation of the metabolically labeled samples on Tris-Tricine gels (TTG), which improves resolution of low-molecular-weight polymers, revealed a ladder of LTA polymers of varying lengths (Fig. 3B). Wild-type cells produced a ladder consistent with a distribution of LTA polymers containing approximately 10–12 repeating disaccharide units, assuming that successive bands correspond to the addition of individual repeat units, consistent with a previous report on LTA length in *C. difficile* 630 (4). Δ*ltaA* cells produced a modestly shorter LTA, and Δ*ltaA* P*_tet_*-*ltaB* cells grown under depletion conditions produced very short LTA polymers.

We next tested whether expression of *ltaA* or *ltaB* could restore LTA synthesis in mutant backgrounds. Expression of *ltaB* from a xylose-inducible promoter restored LTA production in the Δ*ltaB* mutant, as detected by both immunoblotting and metabolic labeling (Fig. 3D-F). In contrast, expression of *ltaA* in the Δ*ltaA* mutant produced longer-than-wild-type LTA, consistent with a role in polymer length control (Fig. 3D-F), whereas expression of *ltaB* in the Δ*ltaB* mutant restored normal LTA length (Fig. 3D-F). Expression of *ltaA* in the Δ*ltaA P*_tet_-*ltaB* depletion strain produced LTA species that were substantially longer than wild type. This difference was especially apparent by anti-LTA immunoblotting, where signal intensity was greater than by in-gel fluorescence of LTA-TAMRA, likely because antibody signal scales with polymer length whereas lipid labeling reflects anchor abundance. These experiments establish LtaB as the primary determinant of LTA production in *C. difficile*, while LtaA influences polymer length. The detection of short LTA species under conditions of limiting LtaB activity is likely due to low levels of LtaB that are insufficient to support normal growth and morphology.

### Residues Important for LtaB Function

Because LtaB is required for detectable LTA production and normal growth in *C. difficile*, we sought to identify residues important for its function. An AlphaFold3 model of LtaB lacking its N-terminal signal peptide predicted a protein with two domains: a large extracellular domain and a C-terminal multi-pass transmembrane domain (Fig. 4A). Confidence scores were high throughout most of the structure, supporting its use for structure-guided mutagenesis (Fig. 4A).

**Figure 4.**
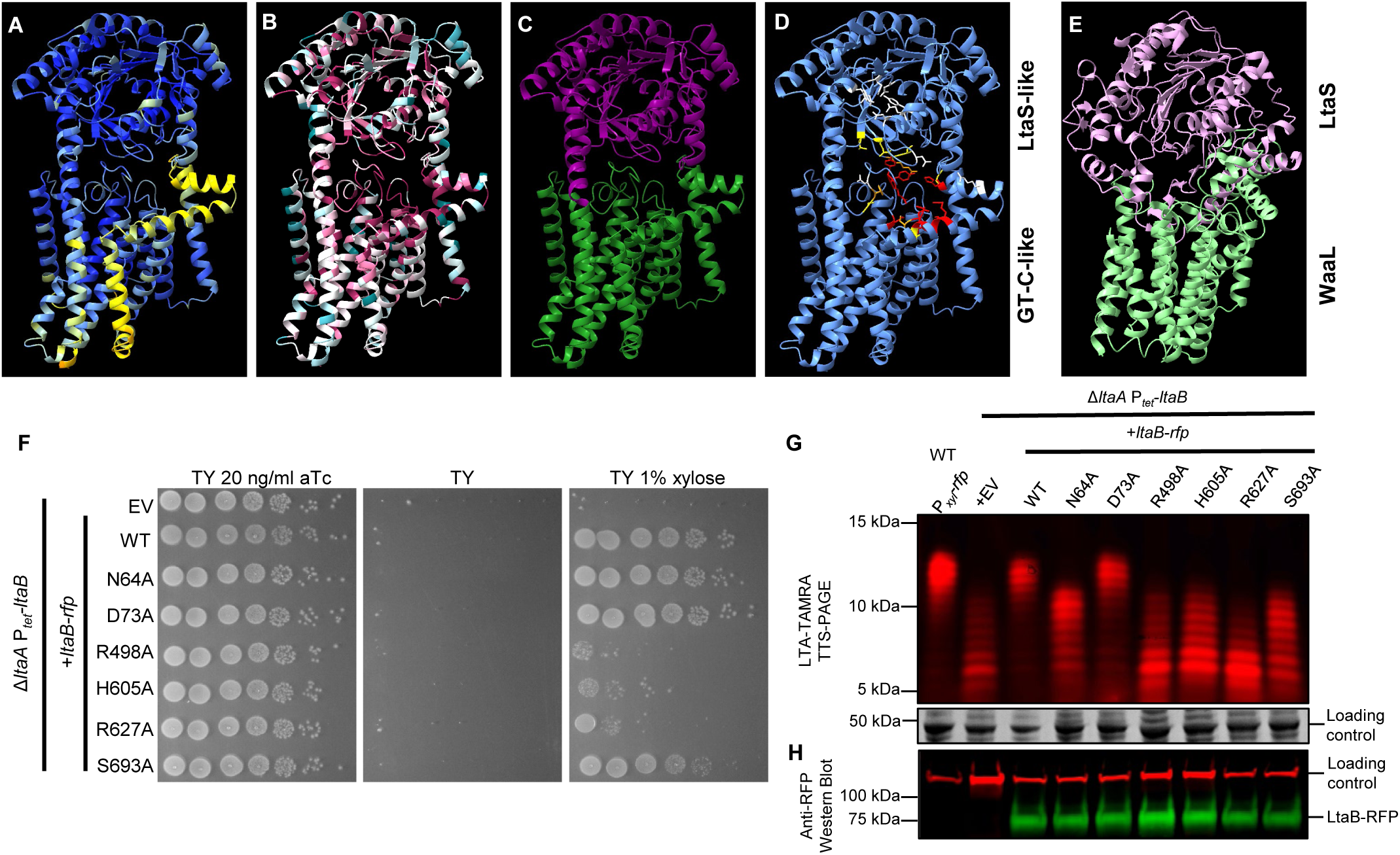
Conserved residues in LtaB are required for LTA synthesis and viability. Structure-guided mutational analysis identifies residues within the transmembrane glycosyltransferase C–like domain of LtaB that are essential for lipoteichoic acid synthesis and cell viability. **(A)** AlphaFold3 structural model of LtaB lacking its signal peptide, colored by predicted model confidence. Dark blue >90 (highly accurate), light blue 89-70 (modeled well), and yellow 69-50 (low confidence, caution). **(B)** ConSurf analysis of LtaB residues based on alignment of 67 homologs from the *Peptostreptococcaceae*, mapped onto the AlphaFold3 structure. ConSurf conservation scores are colored from turquoise (variable) through white (intermediate) to maroon (highly conserved). **(C)** Predicted domain organization of LtaB, highlighting the extracellular LtaS-like domain (purple) and the C-terminal multitransmembrane GTC-like domain (green). **(D)** Locations of residues selected for alanine substitution, colored according to their effects on growth and LTA synthesis. **Red** = severe viability defect and production of short LTA; **Orange** = partial viability defect and production of intermediate-length LTA; **Yellow** = no detectable viability defect and production of intermediate-length LTA; **White** = no detectable effect on viability or LTA synthesis. **(E)** Structural comparison of the LtaB AlphaFold3 model with the type I LTA synthase LtaS from *S. aureus* (PDB 2W5T; lavender) and the O-antigen ligase WaaL from *C. metallidurans* (PDB 7TPJ; light green). LtaS and WaaL were aligned with LtaB in ChimeraX using MatchMaker. **(F)** Viability of Δ*ltaA P*_tet_*-ltaB* strains expressing individual LtaB alanine-substitution mutants under xylose-inducing conditions. **(G)** Metabolic labeling of LTA synthesized by strains expressing LtaB mutant proteins, resolved on Tris-Tricine gels. **(H)** Immunoblot detection of LtaB-RFP fusion proteins demonstrating comparable accumulation of wild-type and mutant variants. All viability and LTA synthesis assays are representative of at least three independent experiments. **Strains used:** WT + P*_xyl_*-*rfp* = AP441; Δ*ltaA P_tet_-ltaB +* EV = CDE5031; Δ*ltaA* P*_tet_-ltaB* + *ltaB*-*rfp* = CDE5231; Δ*ltaA* P*_tet_-ltaB* strains expressing individual LtaB-RFP alanine-substitution mutants N64A = BZ779; D73A = CDE5375; R498A = CDE5369; H605A = CDE5370; R627A = CDE5372; S693A = BZ832 (see Table 3).

To identify conserved regions likely to be important for function, ConSurf was used to identify and map sequence similarity of 67 LtaB homologs from the *Peptostreptococcaceae* family onto the LtaB AF3 structure (Table S2) (48–51). This analysis revealed clusters of highly conserved residues within both the LtaS-like extracellular domain and the predicted glycosyltransferase C (GT-C)-like transmembrane domain (Fig. 4B; Fig. S5). Structural comparison of the AlphaFold3 model of LtaB with the crystal structure of the type I LTA synthase LtaS (PDB 2W5T) from *S. aureus* and the cryo-EM structure of the O-antigen ligase WaaL (PDB 7TPJ) from *Cupriavidus metallidurans* revealed partial overlap with catalytic regions of both enzymes (52, 53) (Fig. 4C,E).

Based on conservation and structural alignment, 30 residues were selected for mutational analysis (Fig. 4D; Fig. S1). These residues were substituted individually with alanine in a fusion of LtaB to an mCherry red fluorescent protein and expressed from a xylose-inducible plasmid (*P_xy_*_l_::*ltaB-rfp*). The LtaB-RFP fusion fully complemented the growth defect of the Δ*ltaA P*_tet_-*ltaB* strain and restored LTA synthesis, indicating that the tagged protein is functional (Fig. 2A; Fig. 4F,G). All mutant LtaB-RFP variants accumulated to comparable levels, as confirmed by immunoblotting with anti-RFP antisera, indicating that the observed phenotypes were not due to protein instability (Fig. 4H; Fig. S6).

Most alanine substitutions within the extracellular LtaS-like domain of LtaB supported viability and synthesis of normal length LTA (D35, N37, D68, D73, Y77, E91, N171 and Y233) (Fig. 4F,G; Fig. S6B,D,F; Fig. S7; Table S3). However, substitutions at a subset of residues within the LtaS-like domain (N64, L293, T311, and T312) rescued the viability defect of the Δ*ltaA P*_tet_-*ltaB* strain but resulted in production of intermediate-length LTA (Fig. 4F,G; Fig. S6B,D,F; Fig. S7; Table S3). These residues cluster near the interface between the extracellular and transmembrane domains, suggesting that this region may influence LTA polymer length or processivity rather than directly catalyzing LTA synthesis.

We found that nearly all mutations in the GT-C-like domain affected LTA synthesis (Fig. 4F,G; Fig. S6B,D,F; Fig. S7; Table S3). LtaB-RFP variants with mutations of R313, S490, Y491, D492, S604, R612, and N632 complemented the viability defect of the Δ*ltaA P*_tet_-*ltaB* strain but produced LTA of intermediate length. In contrast, mutations in R498, Y499, Y500, N504, H605, R627, K628, M631, N691, D692, and S693 were unable to fully complement the viability defect of the Δ*ltaA P*_tet_-*ltaB* strain and produced short LTA similar to the empty vector control (Fig. 4F,G; Fig. S6B,D,F; Fig. S7; Table S3). Most of the residues required for both viability and LTA production surround a highly conserved pocket within the GT-C-like domain (Fig. 4D, shown in red). These findings indicate that the GT-C-like domain is required for LTA synthesis, while the extracellular LtaS-like domain may influence LTA polymer length.

### LtaB Homologs in Other Species

To investigate the phylogenetic distribution and functional conservation of LtaB, we searched the June 2025 release of Fast.genomics, which has a curated main database that includes 7,312 genera of bacteria & archaea and uses mmseq2 for identification of homologs (54). It also has order-level databases with up to ten genomes per species. This analysis identified LtaB homologs primarily within the phyla Bacillota, Actinomycetota, and Armatimonadota (Table 1). There is particularly strong conservation within the order *Peptostreptococcales*, with LtaB homologs present in all examined members of the Peptostreptococcaceae, including *C. difficile* (Table 2; Fig. S8).

**Table 1.** Distribution of LtaB homologs across bacterial phyla. Homologs of *C. difficile* LtaB were identified using MMseqs2 searches of the June 2025 release of the Fast.genomics database. For each phylum with hits, the table lists the number of genomes examined, the number of genomes with at least one LtaB homolog, the total number of homologs identified, and the percentage of genomes with hits.

| Phylum | #Genomes | # Genomes with Hits | # of Hits | % Genomes with Hits |
|---|---|---|---|---|
| Actinomycetota | 809 | 123 | 142 | 15% |
| Armatimonadota | 63 | 14 | 25 | 22% |
| Bacillota | 1300 | 282 | 326 | 22% |

**Table 2.** Distribution of LtaB homologs among families of the order *Peptostreptococcales*. For each family, the table lists the number of genomes examined, the number of genomes with at least one LtaB homolog, the total number of homologs identified, and the percentage of genomes with hits, based on MMseqs2 searches of the Fast.genomics database.

| **Family** | **#Genomes** | **# Genomes with Hits** | **# of Hits** | **% Genomes with Hits** |
|---|---|---|---|---|
| Acidaminobacteraceae | 10 | 0 | 0 | 0% |
| Anaerovoracaceae | 103 | 0 | 0 | 0% |
| Caminicellaceae | 4 | 4 | 4 | 100% |
| Filifactoraceae | 17 | 4 | 4 | 24% |
| Natronincolaceae | 19 | 19 | 20 | 100% |
| Peptostreptococcaceae | 182 | 182 | 296 | 100% |
| T1SED10-28 | 1 | 0 | 0 | 0% |
| Thermotaleaceae | 10 | 10 | 11 | 100% |
| Tindalliaceae | 4 | 0 | 0 | 0% |

To test whether LtaB homologs from other species are functionally conserved, we selected representative homologs from closely related and more distantly related organisms and assessed their ability to complement loss of *ltaB* in *C. difficile*. We included members of the Peptostreptococcaceae family, although we were unable to obtain *P. anaerobius,* which produces an LTA similar to *C. difficile’s* LTA (31). Instead, we used an LtaB homolog from the closely related *Peptostreptococcus* sp. strain MV1. We also included *Paraclostridium bifermentans* which encodes three LtaB homologs: RS04615, RS06385, and RS06390, which are ∼50% identical to *C. difficile* LtaB (Table S4). As more distantly related examples, we selected LtaB homologs from members of the *Brevibacillaceae* family, including *Brevibacillus agri*, *Brevibacillus borstelensis*, and *Brevibacillus laterosporus* which share ∼25% identity (Table S4).

We expressed each homolog from a xylose-inducible promoter in the Δ*ltaA P*_tet_-*ltaB* background and tested complementation by monitoring LTA synthesis and viability. Among the three *P. bifermentans* homologs, RS04615 failed to restore growth and did not support LTA synthesis (Fig. 5A,C). In contrast, RS06385 partially restored growth, while RS06390 restored viability nearly to wild-type levels, although with a small-colony phenotype (Fig. 5A). Strains expressing RS06385 and RS06390 produced LTA polymers that were longer than wild-type *C. difficile* LTA, as detected by both metabolic labeling and immunoblotting (Fig. 5C,D).

**Figure 5.**
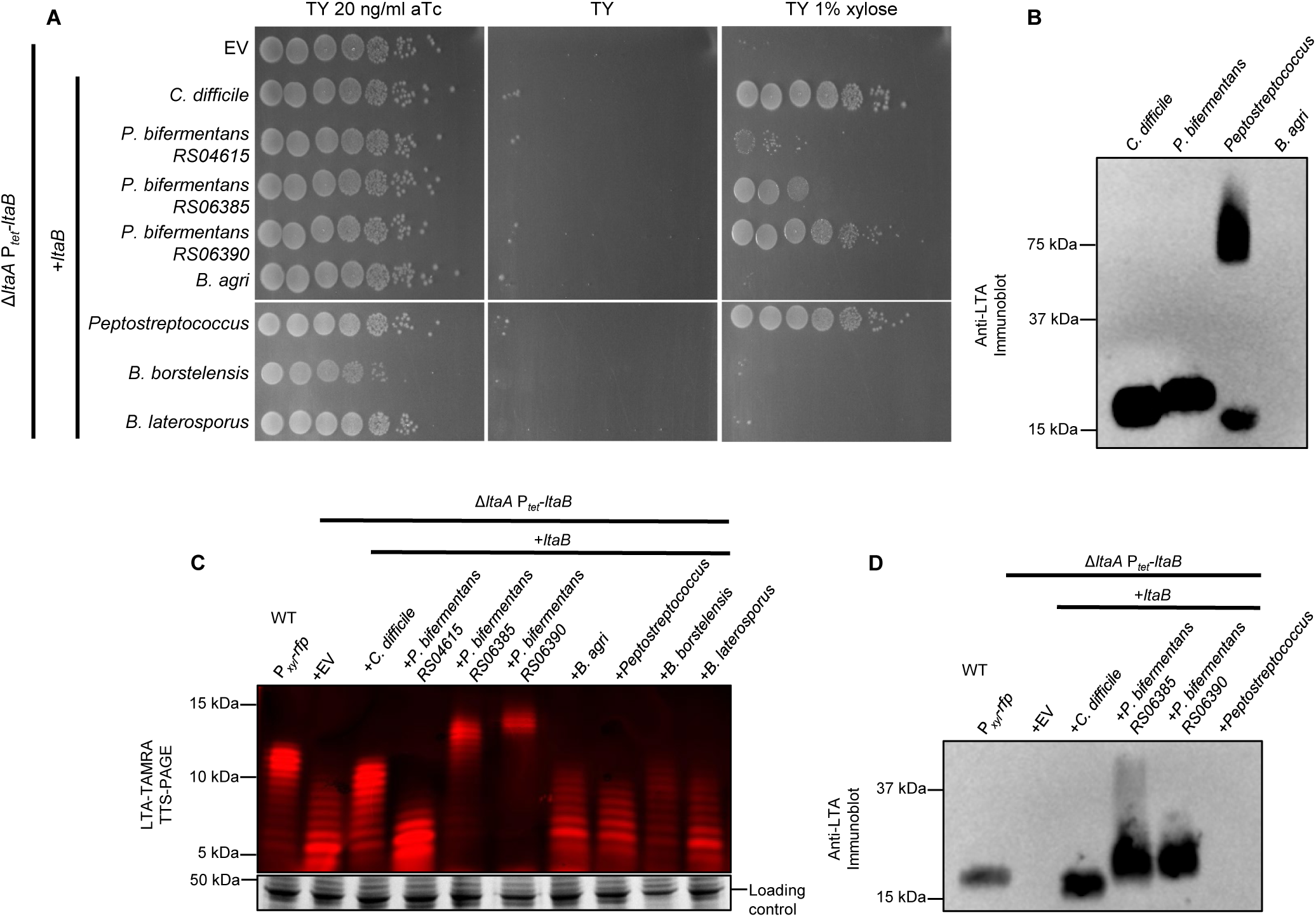
LtaB homologs from closely related species partially substitute for *C. difficile* LtaB. LtaB homologs from members of the *Peptostreptococcaceae* family partially or fully restore viability and/or LTA synthesis in a *C. difficile* LtaB depletion background, whereas homologs from more distantly related species do not. **(A)** Spot-titer viability assays of the Δ*ltaA P_tet_-ltaB* depletion strain expressing the indicated LtaB homologs from a *P_xyl_* expression vector. **(B)** Immunoblot detection of LTA extracted directly from the indicated species. **(C)** Metabolic labeling of LTA synthesized by Δ*ltaA P_tet_-ltaB* strains expressing the indicated LtaB homologs, visualized following click chemistry and Tris-Tricine gel electrophoresis. After fluorescence scanning, gels were stained with Coomassie blue to visualize the major S-layer protein as a loading control. **(D)** Immunoblot analysis of LTA produced by some of the strains shown in (C). The *P. bifermentans* homologs that improved viability also supported production of LTA species with altered polymer length relative to wild type; in contrast, the *Peptostreptococcus* sp. strain MV1 homolog restored viability without detectable LTA by either assay. All results are representative of at least three independent experiments. **Strains used: (A, C-D)** WT + P*_xyl_-rfp =* AP441 (P*_xyl_-rfp*); Δ*ltaA* P*_tet_-ltaB +* EV = CDE5031 (EV); Δ*ltaA* P*_tet_-ltaB* + *ltaB* (*C. difficile*) = CDE5032; Δ*ltaA* P*_tet_-ltaB* + *ltaB* (*P. bifermentans* RS04615, RS06385, RS06390) = CDE5414-CDE5416; Δ*ltaA* P*_tet_-ltaB* + *ltaB* (*Peptostreptococcus* sp. MV1) = CDE5418; Δ*ltaA* P*_tet_-ltaB* + *ltaB* (*B. agri, B. borstelensis, B. laterosporus*) = CDE5417, CDE5530, CDE5531 (see Table 3). **(B)** *C. difficile* R20291; *Paraclostridium bifermentans* ATCC 638; *Peptostreptococcus sp.* Strain MV1 HM-1102; *Brevibacillus agri* DSM 6348 (ATCC 51663).

Expression of the LtaB homolog from *Peptostreptococcus* sp. strain MV1 fully restored viability in the Δ*ltaA P*_tet_-*ltaB* strain but did not restore detectable LTA synthesis by either metabolic labeling or immunoblotting (Fig. 5A,C,D). This result was unexpected because all other homologs that improved growth also restored detectable LTA synthesis. The MV1 homolog therefore appears to restore viability without producing a detectable *C. difficile*-like LTA. In contrast, none of the *Brevibacillaceae* homologs tested restored viability or LTA production in *C. difficile* (Fig. 5A,C,D). Because we did not directly assess protein production, failure to complement could reflect insufficient expression, reduced protein stability, or lack of enzymatic activity.

To assess whether these organisms produced LTAs structurally similar to *C. difficile* LTA, LTAs were isolated directly from representative species and tested for immunoreactivity with the anti-*C. difficile* LTA antibody. We detected LTA in extracts from *P. bifermentans* and *Peptostreptococcus* sp. strain MV1 but not from B. agri (Fig. 5B). *Peptostreptococcus* sp. strain MV1 produced an additional high-molecular-weight species that migrated above the primary LTA ladder (Fig. 5B), suggesting an alternative or modified surface polymer.

## Discussion

As a type V LTA, the lipoteichoic acid of *Clostridioides difficile* differs substantially from the LTAs of well-studied Gram-positive organisms. Rather than a polymer of glycerol phosphate units, *C. difficile* LTA consists of a polysaccharide chain of repeating GlcNAc-GlcNAc-glyceric acid (GroA) subunits joined by 6-*P*-6 phosphodiester bonds and anchored to the membrane by the glycolipid TGDAG (4). The proteins involved in the synthesis of this distinctive LTA had not previously been identified despite the importance of LTA for cell envelope integrity and physiology in many bacteria. Here, we identified two proteins, LtaA and LtaB, that together are required for LTA synthesis in *C. difficile* and establish them as the first characterized proteins of the type V LTA synthase family.

### LtaA and LtaB Contain GT-C-like Domains Important for LTA Biogenesis

Sequence analysis, structural modeling, and mutational studies indicate that LtaA and LtaB belong to the glycosyltransferase C (GT-C) superfamily. GT-C enzymes are integral membrane proteins with multiple transmembrane helices that catalyze the transfer of sugar moieties from lipid-linked donors to acceptor substrates, often in the periplasmic or extracellular space (39). Well-characterized GT-C enzymes include WaaL, which ligates O-antigen to lipid A in Gram-negative bacteria, and MrdB (also called RodA), which transfers peptidoglycan precursors from undecaprenyl-pyrophosphate to the growing cell wall (40, 41).

LtaA and LtaB possess predicted transmembrane domains with structural similarity to members of the GT-C superfamily, particularly WaaL-like proteins (Fig. 4). Mutations at selected conserved residues in the LtaB GT-C-like domain substantially impair LTA production and viability, consistent with an important functional role for this region (Fig. 4B; S6). Whether these residues participate directly in catalysis remains to be established.

These findings suggest that LtaA and LtaB define a previously unrecognized subclass of GT-C-like enzymes involved in type V LTA synthesis. Unlike canonical type I LTA synthases, which use phosphatidylglycerol as a substrate, LtaA and LtaB likely act on a disaccharide-based repeat unit linked to a lipid carrier such as undecaprenol, as in O-antigen biosynthesis. The identity of this lipid-linked precursor and the enzymes that synthesize it remain unknown.

### LtaA and LtaB Play Distinct but Overlapping Roles in LTA Synthesis

Our data demonstrate that LtaB is the primary synthase required for LTA production in *C. difficile*. The absence of LtaB abolishes production of LTA (at least as detectable by currently available methods) and results in severe growth and morphological defects, including cell curvature, chaining, aberrant septation, and lysis. In contrast, loss of LtaA alone does not affect growth or cell morphology but results in formation of a shorter LTA polymer. Depletion of LtaB in a Δ*ltaA* background is lethal, indicating that LtaA and LtaB together perform an essential function.

LtaA influences apparent LTA polymer length in a manner that depends on its expression level. Loss of LtaA results in shorter LTAs, whereas expression of *ltaA* from a xylose-inducible promoter produces longer LTAs than those observed in wild type. This elongation phenotype is even more pronounced when LtaA is expressed under conditions of reduced LtaB activity in the Δ*ltaA P*_tet_-*ltaB* background. These findings indicate that LtaA and LtaB function in the same pathway but perform distinct roles. While LtaB is required for LTA production and normal growth, LtaA modulates polymer length in a dosage-dependent manner and can partially compensate for reduced LtaB activity. These distinct phenotypes raise the question of how LtaA and LtaB cooperate during type V LTA biosynthesis.

### Models for type V LTA synthesis in *C. difficile*

Based on our data, we propose two hypothetical models for LTA synthesis in *C. difficile* (Fig. 6). Both models assume that additional enzymes generate a lipid-linked precursor containing the GlcNAc(GroA)-GlcNAc repeat motif. The first model, analogous to type I LTA synthesis in *L. monocytogenes*, proposes distinct primase and polymerase activities for LtaA and LtaB (Fig. 6A). Under this model, one enzyme initiates synthesis by transferring a disaccharide subunit to the glycolipid anchor TGDAG, whereas the other adds further subunits. The second model, analogous to O-antigen synthesis, proposes that a polymer assembled on a lipid carrier is transferred to TGDAG (Fig. 6B). The structural similarity between LtaB and WaaL supports this possibility but does not establish a ligase reaction.

**Figure 6.**
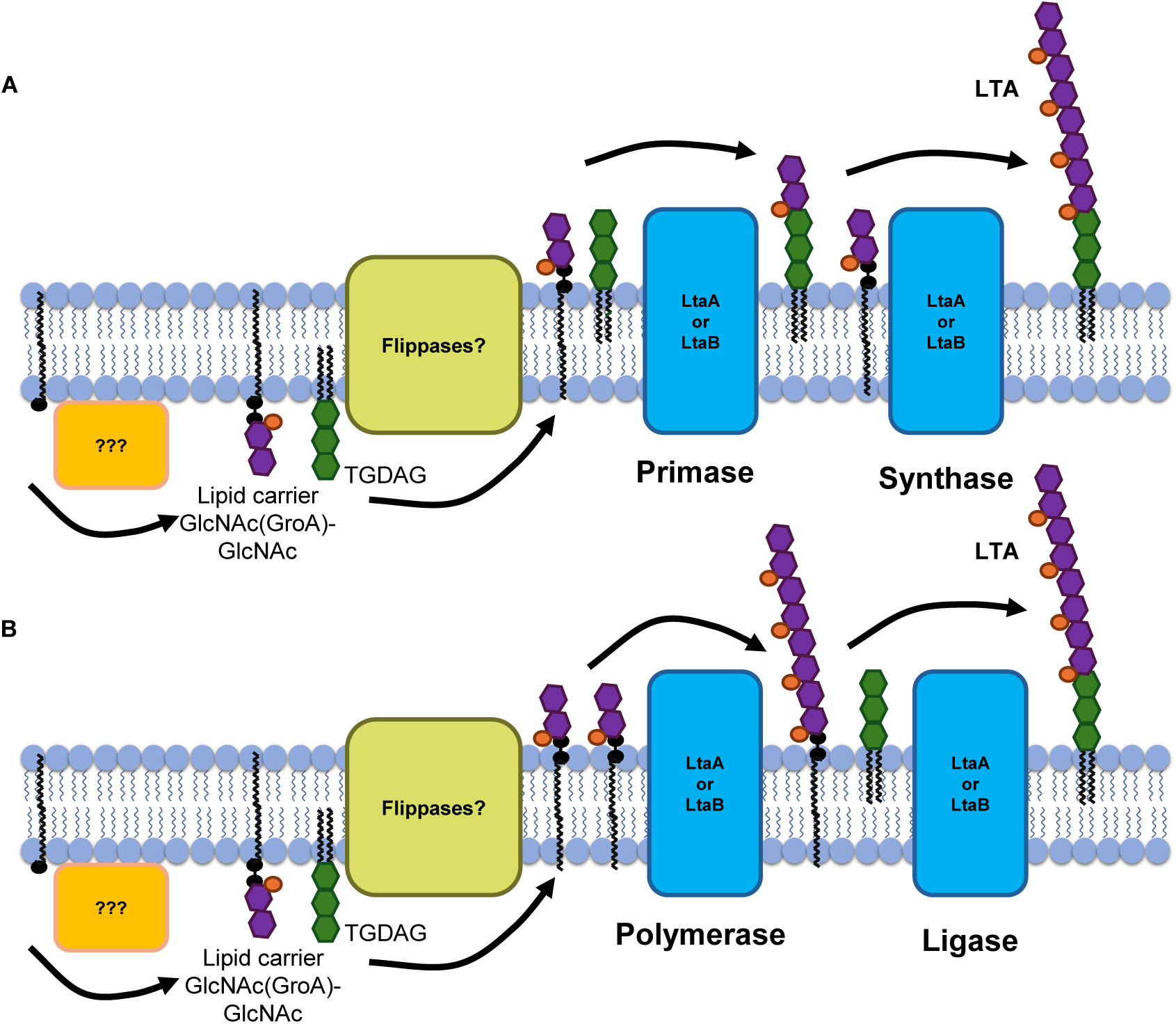
Models for type V lipoteichoic acid synthesis in *C. difficile*. Two non-mutually exclusive models are proposed to explain the roles of LtaA and LtaB in type V lipoteichoic acid (LTA) synthesis and cell envelope homeostasis in *Clostridioides difficile*. Black circles represent phosphates, and wavy lines represent the lipid carrier undecaprenyl (Und); GlcNAc(GroA)-GlcNAc repeat units attached to it are shown as lipid-linked precursors. **(A)** In a primase– polymerase model, LtaA and LtaB are proposed to perform distinct sequential functions during LTA synthesis, with one enzyme initiating transfer of a GlcNAc(GroA)-GlcNAc repeat unit to the glycolipid anchor triglucosyldiacylglycerol (TGDAG) and the other catalyzing polymer elongation. This model is consistent with the shortened LTA observed in strains lacking LtaA and the essential role of LtaB for robust LTA production. **(B)** In a ligase-based model analogous to O-antigen synthesis, the LTA polymer is assembled on a lipid carrier and subsequently transferred *en bloc* to TGDAG by LtaA or LtaB. The strong structural similarity of LtaB to WaaL-family glycosyltransferase C enzymes supports this possibility and might explain why LtaB is essential for LTA synthesis. In both models, the synthesis of the lipid-linked GlcNAc(GroA)-GlcNAc precursor and its transport across the membrane require additional, currently unidentified enzymes.

Our data do not distinguish between these models or identify the individual enzymatic reactions performed by LtaA and LtaB. Altering LtaA expression changes the observed LTA length distribution, but this phenotype does not establish that LtaA directly catalyzes or regulates polymer elongation. More broadly, the proposed lipid-linked GlcNAc(GroA)-GlcNAc precursor, its assembly mechanism, and its transport across the membrane remain hypothetical. The current findings therefore support distinct functional contributions of LtaA and LtaB to type V LTA biogenesis rather than assignment of the proteins to specific biochemical steps. Identifying the relevant intermediates and determining the reactions each protein catalyzes will be essential to define the mechanism of LTA biosynthesis in *C. difficile*.

### Why Is LTA Essential?

An important unanswered question is why LTA synthesis is required for *C. difficile* viability. Strains lacking or depleted for the major LtaB synthase exhibit severe morphological defects, including increased septation, chaining, and impaired cell separation, phenotypes that resemble those observed in LTA synthase mutants from other Gram-positive bacteria (55–57). These similarities support the idea that LTAs play a conserved role in coordinating cell wall synthesis and cell division. The presence of *C. difficile* chains containing multiple septa in our mutants suggests that cell division initiates but daughter-cell separation is compromised.

One possibility is that disruption of LTA biogenesis affects the availability of lipid-linked intermediates required for other envelope biosynthetic pathways. Alternatively, in other organisms, LTAs have been proposed to directly or indirectly influence the activity or localization of peptidoglycan hydrolases (58–60), which contribute to cell division, cell separation, and cell morphology. However, the identities and abundances of the proposed intermediates remain undetermined. We therefore cannot distinguish a requirement for the mature LTA polymer from effects on precursor metabolism or other functions of LtaA and LtaB.

Unexpectedly, the LtaB homolog from *Peptostreptococcus* sp. strain MV1 restored viability to the Δ*ltaA* P*_tet_*-*ltaB* strain despite the absence of detectable LTA by either immunoblotting or metabolic labeling. One possible explanation is that the MV1 homolog produces an LTA species that differs sufficiently from *C. difficile* LTA to escape detection by our assays. Another option is that LtaB’s essential function is linked, at least in part, to the proper metabolism of lipid-linked LTA precursors. In this model, loss of LtaA and LtaB leads to accumulation of a lipid-linked intermediate, sequestering a carrier molecule required for other essential cell-envelope biosynthetic pathways, such as peptidoglycan synthesis. Because undecaprenyl phosphate and related lipid carriers are shared among multiple biosynthetic pathways, disruption of LTA synthesis could indirectly impair cell growth by limiting carrier availability for these processes (8, 61, 62). Expression of the MV1 homolog may relieve this bottleneck by consuming, transferring, or otherwise processing the lipid-linked precursor, thereby regenerating the carrier without producing a *C. difficile*-like LTA polymer. This model offers a simple explanation for how viability can be restored without detectable LTA synthesis and suggests that proper turnover of lipid-linked intermediates may contribute to the essentiality of the LTA biosynthetic pathway. Nevertheless, direct evidence linking LTA synthesis to carrier depletion is currently lacking, and future studies will be required to determine whether the essential role of LtaA and LtaB reflects a requirement for the mature LTA polymer itself, proper metabolism of lipid-linked intermediates, or both.

### Distribution and evolution of type V LTA synthesis

LtaB homologs are predominantly found in organisms from the Bacillota, Actinomycetota, and Armatimonadota phyla. LtaB is present in all examined members of the family *Peptostreptococcaceae*, including *C. difficile*. In the fast.genomics dataset all *Peptostreptococcaceae* encode at least one LtaB homolog, with most encoding 2-3 LtaB homologs. This suggests conservation of an LtaB-related biosynthetic function, potentially involving type V LTA synthesis. In fact, we detected LTAs in both *Paraclostridium bifermentans* and *Peptostreptococcus* sp. strain MV1 that cross-react with our anti-LTA antibodies. In addition, LtaB homologs from *P. bifermentans* partially or fully complemented the loss of LtaB in *C. difficile*. The inability of the *Brevibacillaceae* homologs to complement may reflect functional divergence or a lack of expression in *C. difficile*.

LtaA and LtaB are the first identified enzymes required for type V LTA synthesis. LtaB is required for normal LTA production and growth, whereas LtaA partially compensates for reduced LtaB activity and influences the apparent LTA polymer-length distribution. These proteins have distinct but partially overlapping functions in type V LTA biogenesis, although their individual enzymatic activities remain to be determined.

## Materials and Methods

### Bacterial Strains, Media, and Growth Conditions

Bacterial strains are listed in Table 3. *C. difficile* strains used in this study were all derived from strain R20291. *C. difficile* and *P. bifermentans* were grown in tryptone-yeast (TY) medium consisting of 3% tryptone, 2% yeast extract, and 2% agar for solid medium, supplemented as needed with thiamphenicol (10 µg/mL; Thi10). *Peptostreptococcus* sp. strain MV1 was grown in Reinforced Clostridial HiVeg Broth (3.8%; HIMEDIA) with 2% agar for solid medium. All anaerobic strains were grown at 37 °C in an anaerobic chamber (Coy Laboratory Products) in an atmosphere of ∼2% H₂, ∼5% CO₂, and ∼93% N₂.

**Table 3.**
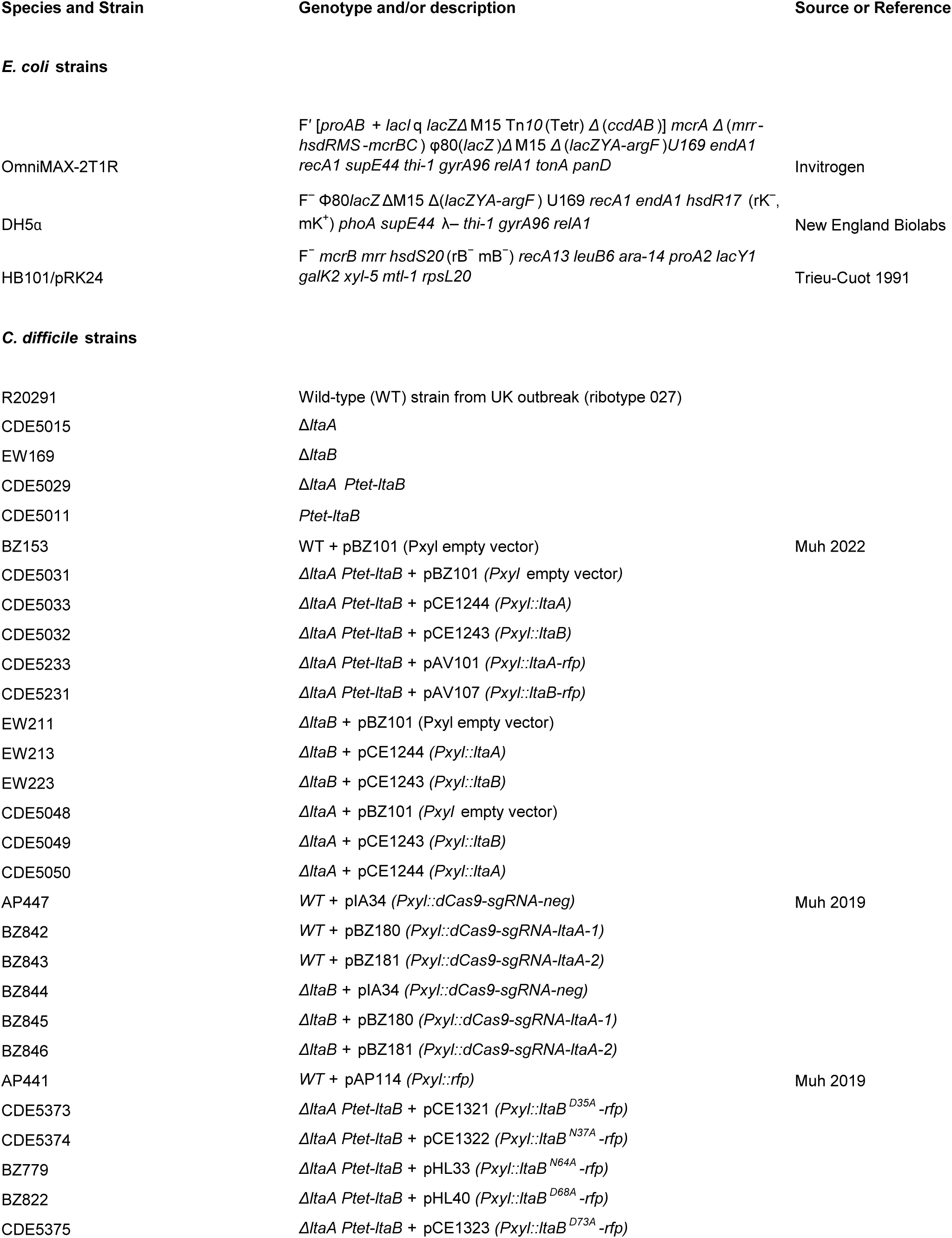
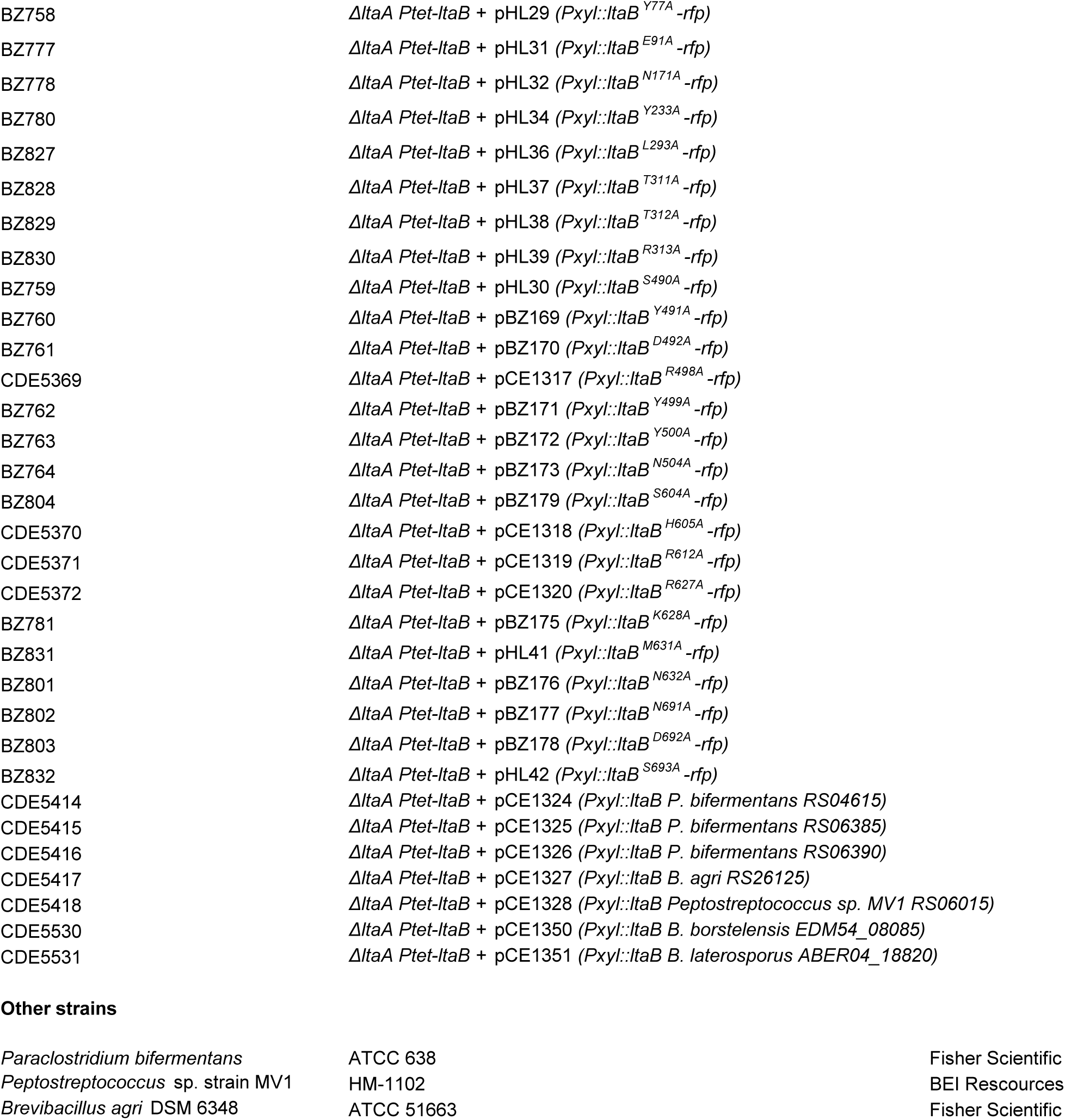
Bacterial strains used in this study. Strains of *E. coli* used for plasmid construction and conjugation, and *C. difficile* strains derived from R20291, are listed with their genotypes or descriptions and their sources or references.

*E. coli* and *Brevibacillus* strains were grown aerobically in lysogeny broth (LB; 1% tryptone, 0.5% yeast extract, 0.5% NaCl) at 37 °C, supplemented as needed with chloramphenicol (10 µg/mL; Cam10) or ampicillin (100 µg/mL; Amp100).

Anhydrotetracycline (aTc) was used to induce gene expression from the P*_tet_* promoter. Unless otherwise indicated, overnight cultures of Δ*ltaA* P*_tet_*-*ltaB* strains were grown in TY containing 5 ng/mL aTc and subcultured 1:50 into TY containing 1 ng/mL aTc for depletion conditions or 10 ng/mL aTc for inducing conditions. Overnight cultures of P*_tet_*-*ltaB* strains were grown in TY containing 1 ng/mL aTc and subcultured 1:50 into TY containing 0.5 ng/mL aTc for depletion conditions or 10 ng/mL aTc for inducing conditions. For strains harboring Pxyl plasmids, xylose was used at 1% (w/v) for induction.

### Plasmid and Bacterial Strain Construction

Plasmids used in this study are listed in Table 4, with additional details provided in Table S5. Plasmids were constructed by isothermal assembly (NEBuilder HiFi DNA Assembly; New England Biolabs). Regions amplified by PCR were verified by DNA sequencing. Oligonucleotide primers were synthesized by Integrated DNA Technologies and are listed in Table S6. All plasmids were propagated in *E. coli* DH5α or OmniMax.

**Table 4.**
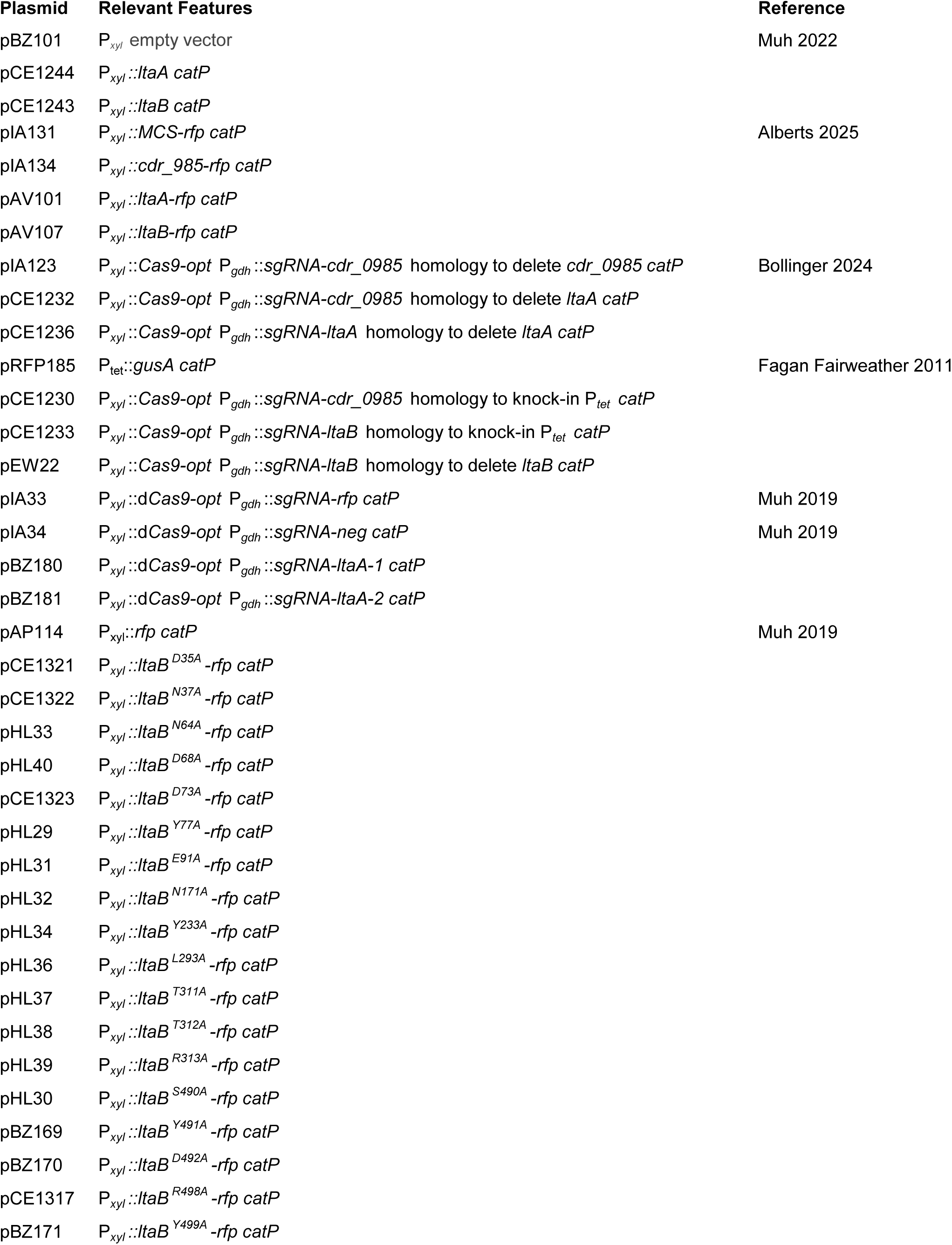
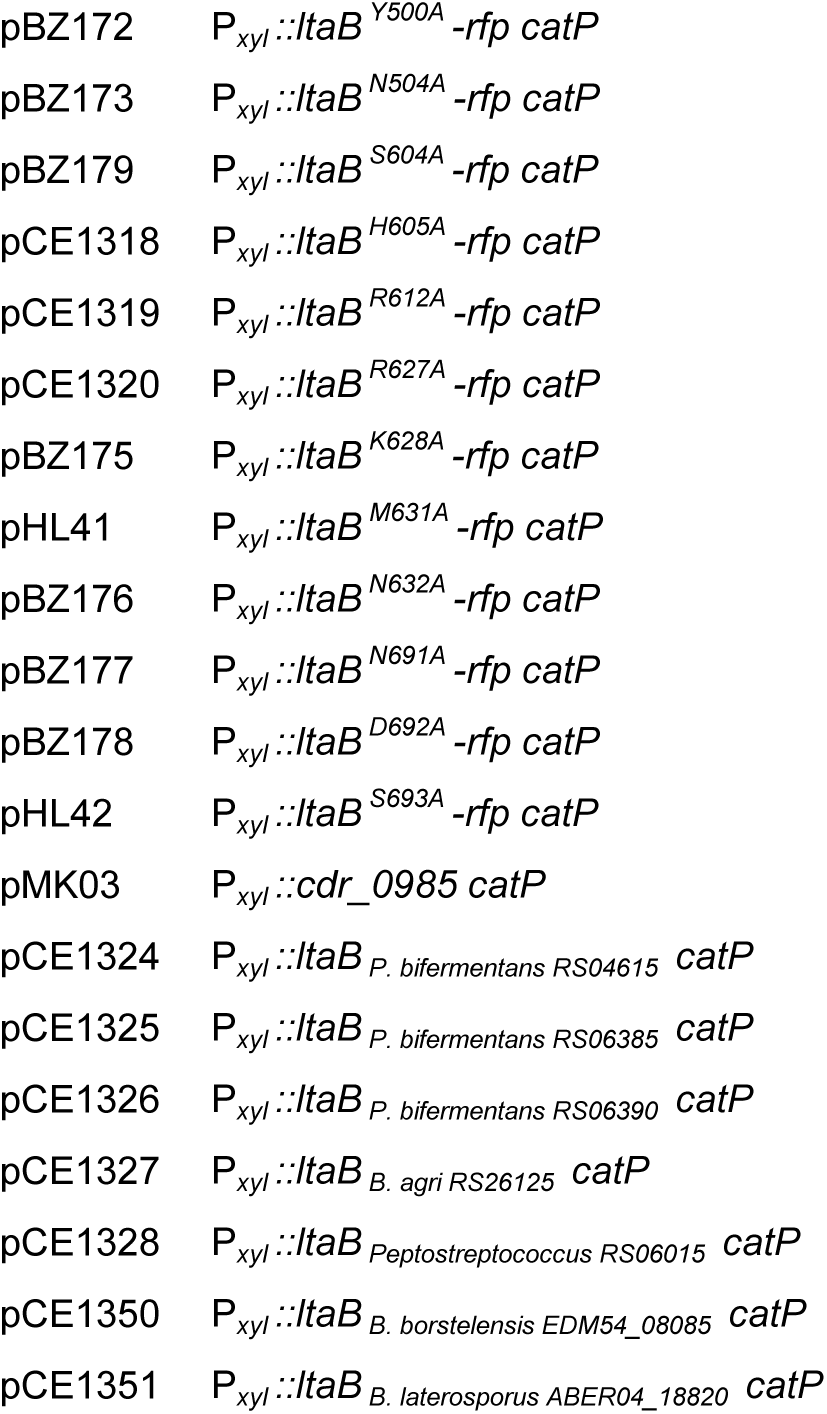
Plasmids used in this study. Plasmids are listed with their relevant features and references.

Genes for xylose-inducible expression in *C. difficile* were amplified by PCR and cloned into plasmids pAP114 (SacI/BamHI) or pMK03 (PmeI/KpnI) (43, 63). CRISPRi constructs were generated by cloning sgRNAs into pIA34 (MscI/NotI) (63). RFP-tagged LtaB constructs were generated by cloning *ltaB* into pIA134 (PmeI/KpnI). CRISPR–Cas9 genome editing plasmids were designed as previously described (64, 65).

Plasmids were introduced into *C. difficile* by conjugation using *E. coli* HB101 containing pRK24, and transconjugants were selected as described previously (65, 66). Successful mutagenesis was confirmed by PCR.

### Viability Assays

Viability was assessed using a spot titer assay. Ten-fold serial dilutions were prepared from overnight cultures, and 5 µL of each dilution was spotted onto TY agar supplemented as indicated. Plates were incubated anaerobically at 37 °C for approximately 18 hours prior to imaging.

### Microscopy

Overnight cultures of *C. difficile* were subcultured to an OD_600_ of 0.05 in TY (supplemented with thi10, 1% xylose, and aTc when needed) and allowed to grow until an OD_600_ of 0.5-0.7. Cells were immobilized using thin 1% agarose pads. Phase-contrast micrographs were recorded on an Olympus BX60 microscope equipped with a 100× UPlanApo objective (numerical aperture, 1.45). Micrographs were acquired using a Hamamatsu Orca Flash 4.0 V2 + complementary metal oxide semiconductor camera. Excitation light was generated with an X-Cite XYLIS LED light source. Membranes were stained with the lipophilic dye FM4-64 (Life Technologies) at 10 µg/mL. Cells were imaged immediately without washing. Red fluorescence was detected with the Chroma 49008 filter set (538-582 nm excitation filter, 587 nm dichroic mirror, and a 590-667 nm emission filter). The MicrobeJ plug-in from the Fiji image analysis package was used to measure cell length and sinuosity (67, 68). Cell sinuosity is the ratio of the cell length along its medial axis to the distance between the poles of a cell. A cell with a sinuosity value of 1 is a perfectly straight rod, whereas curvy cells have sinuosity values larger than 1. At least 300 cells from three independent experiments were used for quantification.

### Time-Lapse Microscopy

Time-lapse microscopy was performed essentially as described previously (69–71). Briefly, exponentially growing cells were spotted onto anaerobic agarose pads prepared in TY medium supplemented with 0.1% (w/v) L-cysteine. Pads were sealed anaerobically using GeneFrame chambers and imaged at 37 °C on a Nikon Ti2 Eclipse inverted microscope equipped with a 100× oil objective (NA 1.45) and an Andor Sona camera. Images were acquired every 5 minutes for 5–6 hours and analyzed using Nikon NIS-Elements software.

### LTA Extraction

LTA was extracted using a modified hot phenol method (4, 45). Overnight cultures were subcultured to an OD_600_ of 0.05 into 200 ml of the appropriate media and grown to mid-log phase, harvested, and normalized by OD_600_. Cell pellets were resuspended in 1 mL of 4% (v/v) phenol in water and incubated for 30 min at room temperature, followed by centrifugation at 16,000 g for 1 min. Pellets were washed once with 1× phosphate-buffered saline (PBS; 137 mM NaCl, 2.7 mM KCl, 10 mM Na2HPO4, and 1.8 mM KH2PO4, pH 7.4), resuspended in 1 mL water, and incubated at 95 °C for 30 min. After centrifugation (3,000 × g, 10 min), pellets were resuspended in 1 mL of 45% (v/v) phenol in water and incubated at 68 °C for 30 min. Samples were centrifuged at 3,000 × g for 10 min, and the aqueous phase was collected. The remaining phenol phase was re-extracted with 600 µL water at 95 °C for 30 min, centrifuged, and the aqueous phase was pooled with the first extract. Combined aqueous fractions were dialyzed against 1 L deionized water at 4 °C using 6 kDa MWCO (Spectrum) tubing for ∼24 h with at least one water change.

### Anti-LTA Antibody

A recombinant anti-LTA IgA antibody was produced using the variable heavy (VH) and variable light (VL) chain domain sequences from the anti-LTA antibody 4D6, which were obtained from Cairns et al. (46). The 4D6 VH gene fragment was synthesized (IDT) and fused to the human IgA2m1 constant region (GenBank AAT74071.1) using overlap extension PCR, along with the tissue plasminogen activator (TPA) signal peptide (residues MDAMKRGLCCVLLLCGAVFVSPAGA). Similarly, we synthesized the 4D6 VL gene fragment (IDT) and fused it to the constant region of human light chain λ1 (UniProt P0DOX8), along with the murine immunoglobulin kappa (IgGκ) light chain signal peptide sequence (residues METDTLLLWVLLLWVPGSTG). The resulting fusions were Electra cloned into expression vector pD2610v13 (ATUM). Resulting expression constructs were co-transfected into Expi293 (Gibco: A14527) using the ExpiFectamine 293 transfection kit (ThermoFisher: A14525) per the manufacturers protocol. Five days following transfection, the supernatant was harvested and purified using Capture-Select IgA affinity resin (Thermo Scientific: 194288010). Eluted protein was buffer exchanged into PBS pH 7.4 and purified using Superdex 200 Increase 10 300 Size Exclusion Chromatography (SEC) (Cytiva) on an AKTA Pure FPLC (Cytiva). Fractions containing 4D6 IgA were pooled and maintained in PBS until use.

### Immunoblot Analysis

For LTA detection, purified LTA extracts were diluted 1:2 in 2X Laemmli sample buffer. Samples were separated on 4–20% Tris-Glycine TGX gels (Bio-Rad) and transferred to PVDF membranes. Membranes were blocked in 5% bovine serum albumin.

LTA was detected using a recombinant 4D6-IgA. The antibody was used at a 1:10,000 dilution, followed by a goat anti-human IgA horseradish peroxidase-conjugated secondary antibody (Invitrogen; 1:10,000). Detection was performed using ECL Prime (Amersham) and an Azure Sapphire Biomolecular Imager.

For detection of LtaB-RFP fusion proteins, whole-cell lysates were prepared by sonication and separated by SDS-PAGE. Blots were probed with anti-RFP antisera (1:10,000) and detected using IRDye 680LT goat anti-rabbit secondary antibody (LI-COR). Streptavidin-IR680LT was used to detect an endogenous biotinylated protein as a loading control. Blots were imaged on an Azure Sapphire Biomolecular Imager.

### Metabolic Labeling and Click Chemistry

Overnight cultures of *C. difficile* were subcultured in 4 mL TY to an OD_600_ of 0.05 and grown in the presence of thi10, 1% xylose, and aTc when indicated. At an OD_600_ of ∼0.3, myristic acid alkyne (25 µM; Vector Laboratories) was added, and cultures were incubated for ∼6 hours. Cells (1 mL of culture at an OD600 of 0.5) were harvested by centrifugation (16,000 × g, 1 min), washed once with 1× PBS, and resuspended in 300 µL of PBS containing 1% SDS. Lysates were prepared by sonication (Branson Sonifier 450; output 3; two cycles of 15 pulses), clarified by centrifugation (16,000 × g, 1 min), and transferred to fresh tubes.

Click chemistry was performed by incubating lysates with 100 µM CuSO₄, 125 µM THPTA (tris-hydroxypropyltriazolylmethylamine), 20 µM TAMRA-azide (5-isomer; Lumiprobe), and 1.2 mM ascorbic acid for 1 hour at room temperature with gentle rocking. Reactions were quenched with 4× fluorescence-compatible sample buffer (Invitrogen), and samples were separated on either 16.5% Tris-Tricine or 4–20% Tris-Glycine TGX gels (Bio-Rad). Labeled species were visualized using an Azure Sapphire Biomolecular Imager (Cy3 channel), followed by Coomassie Brilliant Blue staining to assess total protein loading.

For recovery of metabolically labeled LTA following extraction, cells were labeled with myristic acid alkyne as described above and then processed using the hot phenol LTA extraction procedure. We pooled the aqueous phases from the two phenol extraction steps and dialyzed them against deionized water. The dialyzed aqueous fraction was subjected to click chemistry with TAMRA-azide and analyzed by gel electrophoresis and in-gel fluorescence.

### Alphafold3 Modeling and ConSurf Analysis

LtaB was modeled without its signal peptide using AlphaFold3 with auto-seed settings on the AlphaFold3 server (72). The highest-ranked structure by Predicted Local Distance Difference Test (pLDDT) was used for all subsequent steps. The predicted LtaB structure was used as an input for ConSurf (48–51) with the following parameters: maximum number of homologs: 150, maximum redundancy: 95% identity, MSA building program: Multiple Alignment using Fast Fourier Transform (MAFFT), minimum identity: 35%, homolog selection: sample, E-value cutoff: 0.0001. ConSurf identified 67 high-quality unique hits from other *Peptostreptococcaceae* with a minimum of 35% identity with LtaB (Table S2). AlphaFold and ConSurf structures were rendered in ChimeraX version 1.5 using default settings (73). Structural comparisons between LtaB from *C. difficile*, LtaS (PDB 2W5T) from *S. aureus*, and WaaL (PDB 7TPJ) from *Cupriavidus metallidurans* were performed using the MatchMaker function in ChimeraX (73).

## Acknowledgements

This work was supported by Public Health Service Grants R01AI087834 and R01AI195554 to C.D.E. from the National Institute of Allergy and Infectious Diseases. B.R.Z. was supported by grant T32GM008365 and the University of Iowa Center for Biocatalysis and Bioprocessing. Thank you to Alex Valdez for constructing pAV101 and pAV107. The following reagent was obtained through BEI Resources, NIAID, NIH as part of the Human Microbiome Project: *Peptostreptococcus* sp., Strain MV1, HM-1102. We thank members of the Ellermeier and Weiss laboratories for helpful discussions.

---

**Figure S1. LtaA and LtaB are closely related.** Overall, LtaA and LtaB share 63% amino acid identity and 82% similarity across their full-length sequences, consistent with partially overlapping functions in LTA synthesis. Shown is an alignment of LtaA and LtaB from *Clostridioides difficile* strain R20291 generated using Clustal Omega (74). Identical residues are indicated by asterisks (*), strongly similar residues by colons (:), and weakly similar residues by periods (.). Residues of LtaB subsequently selected for alanine-substitution analysis are in bold and highlighted based on their effects on viability and LTA synthesis: **Red** = severe viability defect and production of short LTA; **Orange** = partial viability defect and production of intermediate-length LTA; **Yellow** = no detectable viability defect and production of intermediate-length LTA; **Bolded but not highlighted** = no detectable effect on viability or LTA synthesis.

**Figure S2. Predicted topology of LtaA and LtaB.** LtaA and LtaB share a conserved membrane topology consistent with membership in the glycosyltransferase C superfamily. **(A,B)** Topology predictions for LtaA (A) and LtaB (B) generated using DeepTMHMM, showing an N-terminal cleavable signal peptide (orange), a large extracellular domain (blue), and a C-terminal region containing multiple predicted transmembrane helices (red). **(C)** Schematic representation of the predicted topology of LtaA and LtaB. Both proteins are predicted to contain an extracellular LtaS-like domain followed by a GTC-like domain with 11 transmembrane helices.

**Figure S3. Requirement for LtaA and LtaB in growth and morphology.** Loss or depletion of LtaB, but not LtaA, results in severe growth and morphological defects. **(A)** Spot-titer viability assays of wild-type and mutant strains grown under the indicated inducing or depletion conditions. **(B–E)** Growth curves of the indicated strains monitored in liquid culture. **(F)** Time-lapse phase-contrast microscopy of Δ*ltaA* P*_tet_-ltaB* cells following depletion of LtaB, showing progressive loss of rod shape and cell lysis. Images are representative of three independent experiments. The growth curve is graphed as the mean ± sd of at least three experiments.

**Strains used: (Α)** WT = R20291; Δ*ltaA* = CDE5015; Δ*ltaB* = EW169; P*_tet_-ltaB* = CDE5011; Δ*ltaA* P*_tet_-ltaB* = CDE5029. **(B)** WT = R20291; P*_tet_-ltaB* = CDE5011. **(C)** Δ*ltaA* + EV = CDE5048; Δ*ltaA* + *ltaA* = CDE5050. **(D)** WT + EV = BZ153; Δ*ltaB* EV = EW211; Δ*ltaB* + *ltaA*= EW213. **(E)** Δ*ltaA* P*_tet_-ltaB* + EV = CDE5031; Δ*ltaA P_tet_-ltaB + ltaA* = CDE5033; Δ*ltaA P_tet_-ltaB + ltaB* = CDE5032 (Table 3).

**Figure S4. LTA Detection Methods. (A)** Sensitivity of LTA detection by immuno-blotting. LTA extracts from wild-type were diluted serially as indicated above the blot and analyzed as described in Methods. **(B)** Sensitivity of LTA detection by metabolic labeling. LTA from wild-type cells grown in the presence of myristic acid alkyne was extracted with hot phenol as described in Methods. Extracts were reacted with TAMRA-azide by click chemistry and serial dilutions were analyzed by in-gel fluorescence. After imaging, the gel was stained with Coomassie Brilliant Blue, revealing the absence of proteins in the hot phenol extracts. **(C)** The fluorescent band that runs at 15-20 kDa is not a lipoprotein. Wild-type cells grown in the presence of myristic acid alkyne were lysed by sonication. Lysates were treated with increasing amounts of Proteinase K as indicated, reacted with TAMRA-azide by click chemistry and analyzed by in-gel fluorescence. After imaging, the gel was stained with Coomassie Brilliant Blue.

**Strains used:** WT = R20291

**Figure S5. Conservation of amino acids in LtaB.** ConSurf was used to map a multiple sequence alignment of 67 LtaB homologs from various *Peptostreptococcaceae* (see Table S2) onto the AlphaFold3 structural model of *C. difficile* LtaB. Highly conserved residues cluster within the predicted glycosyltransferase C–like transmembrane domain.

**Figure S6. Effect of alanine substitutions in the GTC-like domain of LtaB on LTA synthesis. (A,C,E,G)** Immunoblot detection of LtaB-RFP alanine-substitution mutants expressed in the Δ*ltaA P_tet_-ltaB* background. **(B,D,F)** Metabolic labeling of LTA produced by the corresponding strains, resolved on Tris-Tricine gels. Mutations that impair viability are associated with the production of short LTA polymers. All images are representative of at least three independent experiments.

**Strains used:** WT + *rfp* = AP441; Δ*ltaA P_tet_-ltaB +* EV = CDE5029; Δ*ltaA* P*_tet_-ltaB* + *ltaB*-*rfp* = CDE5231; Δ*ltaA* P*_tet_-ltaB* strains expressing individual LtaB-RFP alanine-substitution mutants = (see Table 3).

**Figure S7. Effect of LtaB alanine substitutions on viability.** Specific conserved residues in LtaB are required for cell viability. Spot-titer viability assays of Δ*ltaA P_tet_-ltaB* strains expressing individual LtaB alanine-substitution mutants from a *P_xyl_* plasmid. Mutations within the conserved core of the GTC-like domain result in severe or complete loss of viability. Images are representative of at least three independent experiments.

**Strains used:** WT + *rfp* = AP441; Δ*ltaA* P*_tet_-ltaB* + EV = CDE5029; Δ*ltaA* P*_tet_-ltaB* + *ltaB-rfp* = CDE5231; Δ*ltaA* P*_tet_-ltaB* strains expressing individual LtaB-RFP alanine-substitution mutants = (see Table 3).

**Figure S8. Phylogenetic distribution of LtaB homologs among families within the order *Peptostreptococcales* and closely related lineages.** Tips and inner ring are colored according to family designation as indicated in the key. Filled-in squares in the outer ring denote the presence of an LtaB homolog identified using MMseqs2 searches of the Fast.genomics database.

**Movie S1. Live cell microscopy WT.** Phase-contrast movie of WT *C. difficile* R20291 grown on TY 1% agarose.

**Movie S2. Live cell microscopy of LtaB depletion in the absence of LtaA. Phase-contrast**movie of Δ*ltaA* P*_tet_*-*ltaB* grown on TY 1% agarose.

**Table S1. HHpred analysis of LtaA and LtaB from *Clostridioides difficile*.** Representative structural homologs identified for the N-terminal and C-terminal regions of LtaA and LtaB are listed with their functional annotations and HHpred probabilities.

**Table S2. LtaB homologs used for ConSurf conservation analysis.** Protein sequences used for ConSurf analysis of LtaB were identified from members of the family *Peptostreptococcaceae* and related lineages using homology searches. The table lists the UniProt accession number, organism name, and protein sequence used in the analysis. Homologs were selected according to the criteria described in the Materials and Methods and were used to generate the multiple sequence alignment from which residue conservation scores were calculated and mapped onto the AlphaFold3 model of LtaB (Fig. 4 and Fig. S5).

**Table S3. Phenotypic classification of LtaB alanine-substitution mutants.** Residues were grouped according to their ability to complement growth of the Δ*ltaA P_tet_-ltaB* strain and restore LTA synthesis. White = no detectable effect on viability or LTA synthesis; Yellow = no detectable viability defect and production of intermediate-length LTA; Orange = partial viability defect and production of intermediate-length LTA; Red = severe viability defect and production of short LTA.

**Table S4. Sequence identity of LtaA and LtaB homologs from representative bacterial species.** Protein identifiers, source organisms, protein lengths, and percent amino acid identity to C. difficile LtaA and LtaB are shown.

**Table S5. Plasmids used in this study.** Listed are all plasmids constructed or utilized in this work. Relevant genetic features are indicated for each plasmid, including inducible expression constructs, CRISPR-Cas9 genome editing plasmids, CRISPR interference (CRISPRi) plasmids, and LtaB-RFP alanine-substitution derivatives. Parent vectors, restriction enzymes used for plasmid linearization, primers used for amplification of inserts or mutagenic fragments, PCR template DNA, assembly method, and original references are provided where applicable.

**Table S6. Oligonucleotides used in this study.** Oligonucleotides were used for cloning, homology-arm amplification, CRISPR-Cas9-mediated genome editing, CRISPR interference (CRISPRi), promoter insertion, and generation of LtaB alanine-substitution derivatives. The table lists the oligo identifier, nucleotide sequence, and the purpose of each oligonucleotide. Mutagenic primers used in structure-guided analysis of LtaB are indicated by the corresponding amino acid substitution.

## References

1. Centers for Disease Control and Prevention (U.S.). 2019. Antibiotic resistance threats in the United States, 2019. Centers for Disease Control and Prevention (U.S.).

2. Kirk JA, Banerji O, Fagan RP. 2016. Characteristics of the *Clostridium difficile* cell envelope and its importance in therapeutics. Microb Biotechnol 10:76–90.

3. Ganeshapillai J, Vinogradov E, Rousseau J, Weese JS, Monteiro MA. 2008. *Clostridium difficile* cell-surface polysaccharides composed of pentaglycosyl and hexaglycosyl phosphate repeating units. Carbohydrate Research 343:703–710.

4. Reid CW, Vinogradov E, Li J, Jarrell HC, Logan SM, Brisson J-R. 2012. Structural characterization of surface glycans from *Clostridium difficile*. Carbohydrate Research 354:65–73.

5. Anwar F, Vedantam G. 2022. Surface-displayed glycopolymers of *Clostridioides difficile*. Current Opinion in Microbiology 66:86–91.

6. Calabi E, Ward S, Wren B, Paxton T, Panico M, Morris H, Dell A, Dougan G, Fairweather N. 2001. Molecular characterization of the surface layer proteins from *Clostridium difficile*. Molecular Microbiology 40:1187–1199.

7. Fagan RP, Albesa-Jové D, Qazi O, Svergun DI, Brown KA, Fairweather NF. 2009. Structural insights into the molecular organization of the S-layer from *Clostridium difficile*. Molecular Microbiology 71:1308–1322.

8. Willing SE, Candela T, Shaw HA, Seager Z, Mesnage S, Fagan RP, Fairweather NF. 2015. *Clostridium difficile* surface proteins are anchored to the cell wall using CWB 2 motifs that recognise the anionic polymer PSII. Molecular Microbiology 96:596–608.

9. McBride SM, Sonenshein AL. 2011. The dlt operon confers resistance to cationic antimicrobial peptides in Clostridium difficile. Microbiology (Reading, England) 157:1457–65.

10. Lacotte P-A, Denis-Quanquin S, Chatonnat E, Le Bris J, Leparfait D, Lequeux T, Martin-Verstraete I, Candela T. 2023. The absence of surface D-alanylation, localized on lipoteichoic acid, impacts the *Clostridioides difficile* way of life and antibiotic resistance. Frontiers in Microbiology 14.

11. Ma Z, Pequegnat B, Hodgins D, Sagermann M, Ganeshapillai J, Arar S, Dawson J, Frame NW, Wheadon S, Chow H, Arroyo L, Monteiro MA. 2026. *Clostridioides difficile* water-soluble zwitterionic lipoteichoic acid wsPS-III: structure, synthesis, and immunogenicity. Front Immunol 17:1901286.

12. Woods EC, Nawrocki KL, Suárez JM, McBride SM. 2016. The *Clostridium difficile* Dlt Pathway Is Controlled by the Extracytoplasmic Function Sigma Factor σ V in Response to Lysozyme. Infection and Immunity 84:1902–1916.

13. Schneewind O, Missiakas D. 2014. Lipoteichoic acids, phosphate-containing polymers in the envelope of gram-positive bacteria. J Bacteriol 196:1133–1142.

14. Percy MG, Gründling A. 2014. Lipoteichoic Acid Synthesis and Function in Gram-Positive Bacteria. Annu Rev Microbiol 68:81–100.

15. Jorasch P, Wolter FP, Zähringer U, Heinz E. 1998. A UDP glucosyltransferase from Bacillus subtilis successively transfers up to four glucose residues to 1,2-diacylglycerol: expression of *ypfP* in *Escherichia coli* and structural analysis of its reaction products. Molecular Microbiology 29:419–430.

16. Kiriukhin MY, Debabov DV, Shinabarger DL, Neuhaus FC. 2001. Biosynthesis of the Glycolipid Anchor in Lipoteichoic Acid of *Staphylococcus aureus* RN4220: Role of YpfP, the Diglucosyldiacylglycerol Synthase. J. Bacteriology 183:3506–3514.

17. Gründling A, Schneewind O. 2007. Genes Required for Glycolipid Synthesis and Lipoteichoic Acid Anchoring in *Staphylococcus aureus*. J Bacteriol 189:2521–2530.

18. Gründling A, Schneewind O. 2007. Synthesis of glycerol phosphate lipoteichoic acid in *Staphylococcus aureus*. Proc Natl Acad Sci USA 104:8478–8483.

19. Uchikawa K, Sekikawa I, Azuma I. 1986. Structural studies on lipoteichoic acids from four *Listeria* strains. J Bacteriol 168:115–122.

20. Webb AJ, Karatsa-Dodgson M, Gründling A. 2009. Two-enzyme systems for glycolipid and polyglycerolphosphate lipoteichoic acid synthesis in *Listeria monocytogenes*. Mol Microbiol 74:299–314.

21. Wörmann ME, Corrigan RM, Simpson PJ, Matthews SJ, Gründling A. 2011. Enzymatic activities and functional interdependencies of *Bacillus subtilis* lipoteichoic acid synthesis enzymes. Molecular Microbiology 79:566–583.

22. Jervis AJ, Thackray PD, Houston CW, Horsburgh MJ, Moir A. 2007. SigM-Responsive Genes of *Bacillus subtilis* and Their Promoters. J Bacteriol 189:4534–4538.

23. Eiamphungporn W, Helmann JD. 2008. The *Bacillus subtilis* σ^M^ regulon and its contribution to cell envelope stress responses. Molecular Microbiology 67:830–848.

24. Schirner K, Marles-Wright J, Lewis RJ, Errington J. 2009. Distinct and essential morphogenic functions for wall- and lipo-teichoic acids in *Bacillus subtilis*. EMBO J 28:830–842.

25. Shiraishi T, Yokota S, Fukiya S, Yokota A. 2016. Structural diversity and biological significance of lipoteichoic acid in Gram-positive bacteria: focusing on beneficial probiotic lactic acid bacteria. Biosci Microbiota Food Health 35:147–161.

26. Koch HU, Fischer W. 1978. Acyldiglucosyldiacylglycerol-containing lipoteichoic acid with a poly(3-O-galabiosyl-2-O-galactosyl-sn-glycero-1-phosphate) chain from *Streptococcus lactis* Kiel 42172. Biochemistry 17:5275–5281.

27. Seo HS, Cartee RT, Pritchard DG, Nahm MH. 2008. A New Model of Pneumococcal Lipoteichoic Acid Structure Resolves Biochemical, Biosynthetic, and Serologic Inconsistencies of the Current Model. J Bacteriol 190:2379–2387.

28. Denapaite D, Brückner R, Hakenbeck R, Vollmer W. 2012. Biosynthesis of Teichoic Acids in *Streptococcus pneumoniae* and Closely Related Species: Lessons from Genomes. Microbial Drug Resistance 18:344–358.

29. Gisch N, Kohler T, Ulmer AJ, Müthing J, Pribyl T, Fischer K, Lindner B, Hammerschmidt S, Zähringer U. 2013. Structural Reevaluation of *Streptococcus pneumoniae* Lipoteichoic Acid and New Insights into Its Immunostimulatory Potency. Journal of Biological Chemistry 288:15654–15667.

30. Fischer W, Behr T, Hartmann R, Peter-Katalinić J, Egge H. 1993. Teichoic acid and lipoteichoic acid of *Streptococcus pneumoniae* possess identical chain structures: A reinvestigation of teichoid acid (C polysaccharide). European Journal of Biochemistry 215:851–857.

31. Stortz CA, Cherniak R, Jones RG, Treber TD, Reinhardt DJ. 1990. Polysaccharides from *Peptostreptococcus anaerobius* and structure of the species-specific antigen. Carbohydrate Research 207:101–120.

32. Fischer W. 1994. Chapter 10 Lipoteichoic acids and lipoglycans, p. 199–215. In New Comprehensive Biochemistry. Elsevier.

33. Oku Y, Kurokawa K, Matsuo M, Yamada S, Lee B-L, Sekimizu K. 2009. Pleiotropic Roles of Polyglycerolphosphate Synthase of Lipoteichoic Acid in Growth of *Staphylococcus aureus* Cells. J Bacteriol 191:141–151.

34. Altschul SF, Gish W, Miller W, Myers EW, Lipman DJ. 1990. Basic local alignment search tool. Journal of Molecular Biology 215:403–410.

35. Söding J, Biegert A, Lupas AN. 2005. The HHpred interactive server for protein homology detection and structure prediction. Nucleic Acids Research 33:W244–8.

36. Teufel F, Almagro Armenteros JJ, Johansen AR, Gíslason MH, Pihl SI, Tsirigos KD, Winther O, Brunak S, Von Heijne G, Nielsen H. 2022. SignalP 6.0 predicts all five types of signal peptides using protein language models. Nat Biotechnol 40:1023–1025.

37. Hallgren J, Tsirigos KD, Pedersen MD, Almagro Armenteros JJ, Marcatili P, Nielsen H, Krogh A, Winther O. 2022. DeepTMHMM predicts alpha and beta transmembrane proteins using deep neural networks. bioRxiv 10.1101/2022.04.08.487609.

38. Kinoshita T. 2016. Glycosylphosphatidylinositol (GPI) Anchors: Biochemistry and Cell Biology: Introduction to a Thematic Review Series. J Lipid Res 57:4–5.

39. Alexander JAN, Locher KP. 2023. Emerging structural insights into C-type glycosyltransferases. Current Opinion in Structural Biology 79:102547.

40. Ruan X, Loyola DE, Marolda CL, Perez-Donoso JM, Valvano MA. 2012. The WaaL O-antigen lipopolysaccharide ligase has features in common with metal ion-independent inverting glycosyltransferases. Glycobiology 22:288–299.

41. Taguchi A, Welsh MA, Marmont LS, Lee W, Sjodt M, Kruse AC, Kahne D, Bernhardt TG, Walker S. 2019. FtsW is a peptidoglycan polymerase that is functional only in complex with its cognate penicillin-binding protein. Nat Microbiol 4:587–594.

42. Dembek M, Barquist L, Boinett CJ, Cain AK, Mayho M, Lawley TD, Fairweather NF, Fagan RP. 2015. High-Throughput Analysis of Gene Essentiality and Sporulation in *Clostridium difficile*. mBio 6:e02383–14.

43. Alberts ME, Kurtz MP, Müh U, Bernardi JP, Bollinger KW, Dobrila HA, Duncan L, Laster HM, Orea AJ, Pannullo AG, Rivera-Rosado JG, Torres FV, Ellermeier CD, Weiss DS. 2025. Analysis of essential genes in *Clostridioides difficile* by CRISPRi and Tn-seq. J Bacteriol e00220-25.

44. Charlton TM, Kovacs-Simon A, Michell SL, Fairweather NF, Tate EW. 2015. Quantitative Lipoproteomics in *Clostridium difficile* Reveals a Role for Lipoproteins in Sporulation. Chemistry C Biology 22:1562–1573.

45. Cox AD, St. Michael F, Aubry A, Cairns CM, Strong PCR, Hayes AC, Logan SM. 2013. Investigating the candidacy of a lipoteichoic acid-based glycoconjugate as a vaccine to combat *Clostridium difficile* infection. Glycoconj J 30:843–855.

46. Cairns CM, van Faassen H, St Michael F, Aubry A, Henry KA, Rossotti MA, Logan SM, Hussack G, Gisch N, Hogendorf WFJ, Pedersen CM, Cox AD. 2020. Development and Characterization of Mouse Monoclonal Antibodies Specific for *Clostridiodes (Clostridium) difficile* Lipoteichoic Acid. ACS Chem Biol 15:1050–1058.

47. Broncel M, Serwa RA, Ciepla P, Krause E, Dallman MJ, Magee AI, Tate EW. 2015. Multifunctional Reagents for Quantitative Proteome-Wide Analysis of Protein Modification in Human Cells and Dynamic Profiling of Protein Lipidation During Vertebrate Development. Angew Chem Int Ed 54:5948–5951.

48. Glaser F, Pupko T, Paz I, Bell RE, Bechor-Shental D, Martz E, Ben-Tal N. 2003. ConSurf: Identification of Functional Regions in Proteins by Surface-Mapping of Phylogenetic Information. Bioinformatics 19:163–164.

49. Landau M, Mayrose I, Rosenberg Y, Glaser F, Martz E, Pupko T, Ben-Tal N. 2005. ConSurf 2005: the projection of evolutionary conservation scores of residues on protein structures. Nucleic Acids Research 33:W299–W302.

50. Ashkenazy H, Abadi S, Martz E, Chay O, Mayrose I, Pupko T, Ben-Tal N. 2016. ConSurf 2016: an improved methodology to estimate and visualize evolutionary conservation in macromolecules. Nucleic Acids Res 44:W344–W350.

51. Yariv B, Yariv E, Kessel A, Masrati G, Chorin AB, Martz E, Mayrose I, Pupko T, Ben-Tal N. 2023. Using evolutionary data to make sense of macromolecules with a “face-lifted” ConSurf. Protein Science 32:e4582.

52. Ashraf KU, Nygaard R, Vickery ON, Erramilli SK, Herrera CM, McConville TH, Petrou VI, Giacometti SI, Dufrisne MB, Nosol K, Zinkle AP, Graham CLB, Loukeris M, Kloss B, Skorupinska-Tudek K, Swiezewska E, Roper DI, Clarke OB, Uhlemann A-C, Kossiakoff AA, Trent MS, Stansfeld PJ, Mancia F. 2022. Structural basis of lipopolysaccharide maturation by the O-antigen ligase. Nature 604:371–376.

53. Lu D, Wörmann ME, Zhang X, Schneewind O, Gründling A, Freemont PS. 2009. Structure-based mechanism of lipoteichoic acid synthesis by *Staphylococcus aureus* LtaS. Proc Natl Acad Sci USA 106:1584–1589.

54. Price MN, Arkin AP. 2024. A fast comparative genome browser for diverse bacteria and archaea. PLoS One 19:e0301871.

55. Flores-Kim J, Dobihal GS, Bernhardt TG, Rudner DZ. 2022. WhyD tailors surface polymers to prevent premature bacteriolysis and direct cell elongation in *Streptococcus pneumoniae*. eLife 11:e76392.

56. Kasahara J, Kiriyama Y, Miyashita M, Kondo T, Yamada T, Yazawa K, Yoshikawa R, Yamamoto H. 2016. Teichoic Acid Polymers Affect Expression and Localization of dl-Endopeptidase LytE Required for Lateral Cell Wall Hydrolysis in *Bacillus subtilis*. J Bacteriol 198:1585–1594.

57. Kanampalliwar A, Shah M, Park Y, Jeong B, Lawson PA, Bell M, Fesko EM, Sainato A, Sainato D, Walker S, Bae T. 2025. The role of lipoteichoic acid in *Staphylococcus aureus* cell wall integrity. bioRxiv 10.1101/2025.01.16.633316.

58. Hesser AR, Matano LM, Vickery CR, Wood BM, Santiago AG, Morris HG, Do T, Losick R, Walker S. 2020. The length of lipoteichoic acid polymers controls *Staphylococcus aureus* cell size and envelope integrity. J Bacteriol 202:e00149–20.

59. Flores-Kim J, Dobihal GS, Fenton A, Rudner DZ, Bernhardt TG. 2019. A switch in surface polymer biogenesis triggers growth-phase-dependent and antibiotic-induced bacteriolysis. eLife 8:e44912.

60. Sassine J, Sousa J, Lalk M, Daniel RA, Vollmer W. 2020. Cell morphology maintenance in *Bacillus subtilis* through balanced peptidoglycan synthesis and hydrolysis. Sci Rep 10:17910.

61. van Heijenoort J. 2007. Lipid intermediates in the biosynthesis of bacterial peptidoglycan. Microbiol Mol Biol Rev 71:620–635.

62. Chu M, Mallozzi MJG, Roxas BP, Bertolo L, Monteiro MA, Agellon A, Viswanathan VK, Vedantam G. 2016. A *Clostridium difficile* Cell Wall Glycopolymer Locus Influences Bacterial Shape, Polysaccharide Production and Virulence. PLoS Pathog 12:e1005946.

63. Müh U, Pannullo A, Weiss D, Ellermeier C. 2019. A xylose-inducible expression system and a CRISPRi-plasmid for targeted knock-down of gene expression in *Clostridioides difficile*. J Bacteriol 201:e00711–18.

64. Kaus GM, Snyder LF, Müh U, Flores MJ, Popham DL, Ellermeier CD. 2020. Lysozyme Resistance in *Clostridioides difficile* Is Dependent on Two Peptidoglycan Deacetylases. J Bacteriol 202:e00421–20.

65. McAllister KN, Bouillaut L, Kahn JN, Self WT, Sorg JA. 2017. Using CRISPR-Cas9-mediated genome editing to generate *C. difficile* mutants defective in selenoproteins synthesis. Sci Rep 7:14672.

66. Müh U, Ellermeier CD, Weiss DS. 2022. The WalRK Two-Component System Is Essential for Proper Cell Envelope Biogenesis in *Clostridioides difficile*. J Bacteriol 204:e00121–22.

67. Schindelin J, Arganda-Carreras I, Frise E, Kaynig V, Longair M, Pietzsch T, Preibisch S, Rueden C, Saalfeld S, Schmid B, Tinevez J-Y, White DJ, Hartenstein V, Eliceiri K, Tomancak P, Cardona A. 2012. Fiji: an open-source platform for biological-image analysis. Nat Methods 9:676–682.

68. Ducret A, Quardokus EM, Brun YV. 2016. MicrobeJ, a tool for high throughput bacterial cell detection and quantitative analysis. Nat Microbiol 1:16077.

69. Ribis JW, Nieto C, DiBenedetto NV, Mehra A, Kuhn P, Dong Q, Nagawa I, El Meouche I, Aldridge BB, Dunlop MJ, Tamayo R, Singh A, Shen A. 2025. Unique growth and morphology properties of Clade 5 *Clostridioides difficile* strains revealed by single-cell time-lapse microscopy. PLoS Pathog 21:e1013155.

70. Kurtz MP, Müh U, Weiss DS, Ellermeier CD. 2026. CenIR, an essential BlaIR-family regulatory system in *C. difficile*. J Bacteriol 208:e0019626.

71. Bollinger KW, Müh U, Brannen PB, Popham DL, Weiss DS, Ellermeier CD. 2026. Carboxypeptidase activity drives L,D-transpeptidase essentiality during vegetative growth and sporulation in *Clostridioides difficile*. mBio e0158026.

72. Abramson J, Adler J, Dunger J, Evans R, Green T, Pritzel A, Ronneberger O, Willmore L, Ballard AJ, Bambrick J, Bodenstein SW, Evans DA, Hung C-C, O’Neill M, Reiman D, Tunyasuvunakool K, Wu Z, Žemgulytė A, Arvaniti E, Beattie C, Bertolli O, Bridgland A, Cherepanov A, Congreve M, Cowen-Rivers AI, Cowie A, Figurnov M, Fuchs FB, Gladman H, Jain R, Khan YA, Low CMR, Perlin K, Potapenko A, Savy P, Singh S, Stecula A, Thillaisundaram A, Tong C, Yakneen S, Zhong ED, Zielinski M, Žídek A, Bapst V, Kohli P, Jaderberg M, Hassabis D, Jumper JM. 2024. Accurate structure prediction of biomolecular interactions with AlphaFold 3. Nature 630:493–500.

73. Pettersen EF, Goddard TD, Huang CC, Meng EC, Couch GS, Croll TI, Morris JH, Ferrin TE. 2021. ChimeraX: Structure visualization for researchers, educators, and developers. Protein Science 30:70–82.

74. Sievers F, Wilm A, Dineen D, Gibson TJ, Karplus K, Li W, Lopez R, McWilliam H, Remmert M, Söding J, Thompson JD, Higgins DG. 2011. Fast, scalable generation of high-quality protein multiple sequence alignments using Clustal Omega. Molecular Systems Biology 7:539.

