## Supplementary material for "Identification of two novel lipoteichoic acid synthases essential for viability in *Clostridioides difficile*": Table S1

**Table S1. HHpred analysis of LtaA and LtaB from *Clostridioides difficile***

| Region | Representative HHpred hit | Functional annotation | LtaA Prob. (%) | LtaB Prob. (%) |
|---|---|---|---|---|
| N-terminal | 8DI0_C | Arylsulfatase / sulfatase-fold protein | 99.7 | 99.7 |
| N-terminal | cd16024 | GPI ethanolamine phosphate transferase | 99.7 | 99.7 |
| N-terminal | cd16015 | Lipoteichoic acid synthase family | 99.5 | 99.6 |
| N-terminal | 4UOR_C | Lipoteichoic acid synthase | 99.5 | 99.5 |
| N-terminal | 5I5D_A | YejM-family inner membrane protein | 99.5 | 99.5 |
| C-terminal | 7TPG_B | WaaL O-antigen ligase, Cupriavidus metallidurans (GT-C) | 93 | 94 |
| C-terminal | 9RMK_A | Oligosaccharyltransferase; membrane glycosyltransferase (GT-C) | 86.8 | 82.4 |
| C-terminal | PF13425 | WaaL-family O-antigen ligase; membrane glycan ligase (GT-C) | 58.5 | 61.2 |
| C-terminal | 25HS_A | FtsW Peptidoglycan-associated integral membrane protein (GT-C) | 44.3 | 50.4 |
| C-terminal | 8TJ3_B | Peptidoglycan glycosyltransferase MrdB/RodA (GT-C) | 44.4 | 41.2 |
