## Supplementary material for "Identification of two novel lipoteichoic acid synthases essential for viability in *Clostridioides difficile*": Table S2

**Table S2. LtaB homologs used for ConSurf conservation analysis**

| Accession | Scientific Name | Sequence |
|---|---|---|
| UPI001D125D26 | Clostridioides sp. ZZV13-5731 | EEKGKVIFIDMNRTNLGSMLKVPILKSELEKRGHIALMNIRGDQGTDDRRSYASMGAGTRADVTTEDYINFQKVDQDTAKTFEVATGQKAKEINNLSI<br>NRSINENLEYGEYGASLGSLGKTLSENGLKTSVIGNADIIENNQLVKNRNLCLVAMDQYGRIDSGNIDDINKKDSKMPFGIATDYNKLLDETKKSYKN<br>SDAIFVELGDTYRLDVYRLNLNDNTYAAMKKNIVKNIDKYLKEVFSMAGENDTVYISSAFPSDLDYKNKRRLSPIIKFNGEGKGLLESATTRKAGIVANL<br>DVGVDILNEFGLKNDEMVGRSYSLVDREDNANYLQSLYKKIVSISSIRSSIINGFVGVISASWVIAMFAIFFRNYMPHKERVFKVLKEFIKLGIIMPLSFM<br>LAPLMNFKTPITISLGIFVTTLVLYLLGRKLFKDNDIKHMGFFATLTILVIVIDSVFGTYLMQNNIMSYDAVIGARYYGVGNEYEGVTIASAVFGLAVLLHYK<br>KVSKPLVVIFSLIILITSAYPSMGANVGGAISECVAYLLFIMLIYDVKLDIKKIALLAISAVVVVCAFAALDIISGSESHLGMFVQRIFLNGPGEILQTFGRKIQ<br>MNMQLAQTSVWVNILLVGIGVIAVLIFKPSRHFRKIMNNFPILFKGFIASMVGCIITLLVNDSGIVAASTASIYILIPLIIISINMIIFEKNN |
| UPI00215A9007 | Romboutsia lituseburensis | DEKGKVIFINMNRTNLVSMLHIQDLDKEVQERGYVGLMNIRGDKGTDDRRSYASMGAGGRANVSTEQNINFQAADKDSAKLFEAATGKKAKAINDL<br>TVNKSINENIEKGQYGSILGSLGQTLSENGFKTAVLGNSDIVENGELVKNRNIGLVAMDHNGRIDAGNVDDINIKDFTMPYGIRTDYDKLINETKKYYK<br>ENDVLFVDLGDTYRLDNYRLNLNEKTYSKMKHSIGKNIDKYLKEVFDMVGEKDTVYISSAFPSDLDYKNKRRLSPIIKFSGEGKGILSSATTRREGIVAN<br>LDVGVDILNEFGLKNETMVGRTYNLVEKEDNVKYVLDEFEKIVSISNIRATVVNGFVGVISASWVIAMILILLRNYLPHKEKVFIILKEFIKLGIIMPLSFLLA<br>PLFNFKQPMTITAGILFTTLVLYILGRALFKDDDIKQMGFFAFITIGVITIDSAFGTFLMQNNIMSYDAIIGARYYGVGNEYEGVTIACAVFALAVLLNYKKIP<br>KWLVVVLSIVILITSAYPSMGANVGGAISECVAYLLFILLIYDMKLDFKKVILLGFAAVAVVAIFAFLDIVSGSESHLGMFVQQILLNGPMTVIQTFTRKIQ<br>MNIKLAQTSVWVNILLAGIAIIAVFIFRPSKHFRDIAKKYPALFKGFVASMVGCIVTLLVNDSGIVAASTASIYILIPVIIISINMIIFGDIN |
| A0A2P2BWR6 | Romboutsia hominis | DEHGKVIFINMNRTNLNTMLSVPTLKKELKNRGYVALMNIRGDGGTDDRRSYASMGAGGRVNVSAQEYIDFQELTKENGKVFKSVTGKEPKAINNID<br>INKSINDNIEYGQYGSMLGALGQTLSENNLKTAVLGNSDKIENGELVKNRNIGLVAMDEYGRIDGGNIDNINVKDESMPYGIRTDYNKLIKETQKYYN<br>ESDAIFVDLGDTYRLDQYRLFLNDNTYKSMKANIVKNINKYLKEVFNMVGENDVVYISSAFPNDLDYKNRRRLSPIIKFEGNKKGLLSSATTRREGIVA<br>NIDVGVDILNEFGLKNKNMVGRSYELIQKDDNIGYIQHEFVKMVSVSDIRANVVNTFVGIVSASWVIGMIAILLRKYVPYKERVFNVLKELIKLGIIMPLA<br>LLLAPVFNFKTPMTITAGIVLTTLILYILGRKLFKDDIKQMGFFALVTIIVIVIDSVFGTYLMRNNIMSYDAIIGARYYGIGNEYEGVTIASAVFAMAVLLNYKK<br>IPKWLVALMSIVILITSAYPSMGANVGGAISESVAYLLFILLIYDVKLDFKKVILLGVAAVAVVVAFAVLDVVSGSESHLSVFVNQILVNGPGAILQTFGRKI<br>QMNMKLAQTSVWVNILLAGIFIIAMFIFKPPRHFRDIAKKYPILFKGFIASMVGCIVTLLVNDSGIVAASTASIYILIPIIIISINMIIFNKED |
| UPI001AE1E881 | Metaclostridioides mangenotii | EEKGKVVFINMNRTNLVSMRKVPTLKKELDNRGYIALMNIRGDKGTNDRRSYASMGAGGRANVTkDEDVINFGNVSKENAKIFEAATGKKAKGINYL<br>NVNRSINDNFENGEFGSTLGKMGQTLSDNNLKTAVIGNSDTLENGETVKNRDLCLLAMDEYGRIDMGNVDDINIKDTSMPYGISTDYEKLTNETKEY<br>YSKSDALFVDLGDAYRLDAYKGSLNEDTYESMRKNVDEKIDNYLKEVFNMVGENDVVYVVSGFPSNLDYKNKRRLSPVVKFEGTGKGLLTSSTTRQ<br>NGIVANLDIGVDILDEFGIKSDSMVGRSFSNVDKEENVEYLLGEYEKIVSISNIRGTIINIFVSVVSVSWIIGTVAILFRNRFRHKEKVFNVLKELIKLGIIMP<br>LSFMIAPLMNFKTPMGILLGIFAVALGLYVIGRVFIKDDLKHMGFFAGVTIALIVIDSMFGTFMMKNNIMSYDAIVGARYYGVGNEYEGVTIASAIFALSV<br>LMNYKKIPKWSVIVISLIILITSAYPSMGANVGGAISECVAYLLFILLIYDVKIDFKKIILLGLSAVVVVAVFAFLDIKTGSGSHLGAFVQQIMVNGPGEIIQTF<br>VRKIQMNVKLAQTSVWVNILLAGIAIIAVLIFRPSKHFKKVMVDYPIIFKGFISSMVGCIITLLVNDSGIVAAATASIYILIPIVIISINMIIFNKEN |
| UPI001C10A6BD | Metaclostridioides mangenotii | EEKGKVVFINMNRTNLVSMRKVPTLKKELDNRGYIALMNIRGDKGTNDRRSYASMGAGGRANVTKDDDvINFGNVNEENAKIFEAATGKKAKGINYL<br>NVNRSINDNFENGEFGSTLGNMGQTLSDNNLKTAVIGNSDTIENGEVVVNRDLCLLAMDQYGRIDMGNVDNINKKDTSMPYGISTDYDKLLKETKD<br>YYSKSDALFVDLGDAYRLDAYKGSLNEETYESMRDNVDKKIDKYLKEVFNMVGENDVVYVVSGFPSNLDYKNKRRLSPVVKFEGAGKGLLTSSTTRQ<br>NGIVANLDIGVDILNEFGLKSETMVGRSLSTNDKEENIEYLLGEYEKIVSISNIRGTIINIFVSVVSISWVIAAVAILFRNRFKHKEKVFNVLKEFIKLGIIMPL<br>SFMIAPLLNFKTPMGILLGIFAVAVVLYIIGRLFIKDDLKHMGFFAGLTIALIVIDSVFGTFMMKNNIMSYDAIVGARYYGVGNEYEGVTIASAIFALSVLM<br>NYKKIPKWTVIVISLIILITSAYPSMGANVGGAISECVAYLLFILLIYDIRIDLKKVLLLGLSAVVVVAVFAFLDLKTGSGSHLGGFVHQIMINGPGEIIQTFV<br>RKIQMNVKLAQTSVWVNILLAGIGIIAVLIFKPSKHFRKVMKDYPIIFKGFVSSMVGCIITLLVNDSGIVAAATASIYILIPIVIISINMIIFNKEN |

| | | |
|---|---|---|
| A0A1M5K4S3 | Asaccharospora irregularis DSM 2635 | EEKGKVIFISMDRTNLVSMMHIQALKNELEYRGYIGLMNIRGDKGTDDPRSYASMGAGCRANLmVKDEYIDFTNVDVDAAKTFEVATGKKAKKINYLT<br>INKSINENFENGEYGATLGSLGQTLSENGLKTAVIGNADIIQNGELIKNRNIGLIAMDNYGRIDAGNVDDIDKKDPYMPFGISTDYDKLIKETKKYYSEN<br>DAIFVELGDTYRLDMYRLALNEDTHAKMKHKIGKNISKYLDEVFKMVGENDTVYITSPFPSDLDYKNRRRLSPIIKIKGEGKGILTSSTTRREGIVANLDV<br>GVDILTELGLKNDVMIGRSLKNIEKEDNSDYLLGEFEKIVSISNIRAGVINTFVGIISASWVIAMFAVIFRKRIPNNERVFPVLKEFIKLGIIMPLSFLLSPIM<br>NLKTPLTISAGIILTTIIFYIIGKKFFKDDLKHMGFFAGLTILLITIDSIFGTYLMQNNIMSYDAIIGARYYGIGNEYEGITIASAIFSLAVLLNYKRIPKWLVAVFS<br>GIILIVSAYPSMGANVGGAISECVAYLLFLLLIFDVKLDFKKFLLLGLAAVGVVVLFAVLDIVSGSESHLAVFTNQILLEGPSAIIQTFTRKIQMNLKLAQTS<br>VWVNILLVGIAIIAGLIFKPSKHFKNISDRYPVIFKGFVASMVGCIVTLLVNDSGIVAASTASIYILIPIIIMSINMIIFEDK |
| UPI0004B23C52 | Faecalimicrobium dakarense | DEKGKVIFIDMNRTNLINMLKIPSLENELSTRGYIGLMNIRGDKGTDDRRSFASMGAGGRANVSTEMNINFEEINEDTKATFKAATGKSPKKINDLTIN<br>QSINENIENGQYGSVLGSLGQTLSDNKLKVALLGNSDIIENGELVKNRNLGLVCMDQYGRIDAGNIDNINTKNSSMPFGIQTDYDKLIKETKENYKSS<br>DALFIELGDTYRLDLYKGYLNDHTFASMKKNVYKNIDKYLKEVFNMVGENDTVYIASGFPSDIDYKNKRRLSPIIKFKGDKKGLVLSATTRREGIVANIDI<br>GVDILNEFGLKNDAMVGRSYSLVEKDDNVEYLSDEFEKIVTIADIRANVINGFVSVISISWVVAMCLIFIRNKITNQEKVFSVLKEFIKLGIIMPLSFLVAPI<br>LNLQTPVTITCGILLTTLAFYILGRWLFKEDLKQMGFFAFLTILLITVDSVFGTYLMQNNIMSYDAIIGARYYGIGNEYEGVTIASAVFALAVLLNYKKIPKW<br>LVIVFSLIILITSAYPSMGANVGGAISECVAYLLFILLIFDVKMDFKKVVLLGLAAVAVVAIFAALDIVSGSESHLGLFVKQIFMNGPEAIIQTFGRKIQMNM<br>KLAQSSVWVNILLAGIGIIGVLIFRPSRHFRTIAKKYPILFKGFVASMVGCIVTLLVNDSGIVAAATASIYILIPLIIISINMMIFDDK |
| UPI00215B6EB9 | Romboutsia lituseburensis | DKNGKVIFINMNRTNLASMLHIKSLEKEVQNRGYVGLMNIRGDKGTDDRRSYASMGAGRRANVSTEQNINFQSVDNDSAKLFETATGQKSKSINDL<br>TINKSINENIENGQYGSTLGSLGQTLSENGLKTAVLGNSDIVENGELIKNRNIGLVAMDQYGRIDSGNIENINVKDNTMPYGIRTDYNKLIKETKKYYEE<br>NDALFVDLGDTYRLDNYRLNLNEKTYSNMKSKIGNNIDKYLKEVFNMIDENDTVYISSAFPSDLDYKNKRRLSPIIKFNGDGKGVLSSATTRREGIVAN<br>LDVGADILNTFGFENDEMVGKEYNLIKKDGNIKFVLDEYEKIVSVSKIRPNVVNTFVSIVSASWVIAMVILLFKDRVKYKEKVFKVLKEFIKLGFILPISFFIA<br>PIFNFKTPETIVFGIVLTTLILYIIGIKLFKNDLKHMGYFAVISVLAIVIDSILGSYLMKNSIMSYDAIIGARYYGMGNEYQGIVIGSSIFGIAVLLHYNKISKWL<br>AAISCFVILITTASPTMGANVGAAISECMAFLLLVLLVFDVKLDFKKIILIGLAGVGIVFAFAVIDIISGSESHLSGFVNQILLDGPSAIIQTFSRKIQMNIKLA<br>QSSAWVNILLAGIAIIAIFIFKPSKHFRNIANKYPFIFKGFIACIVGCFITLLVNDSGIVAAATTSIYILIPLLIISINMIIFNEK |
| A0A1G9Q2K8 | Romboutsia lituseburensis DSM 797 | DQNGKVIFINMNRTNLASMLHIKSLDEEVQNRGYIGLMNIRGDKGTDDRRSYASMGAGRRANVSTEQNINFQSVDDHSSKLFETSTGQKSKSINDL<br>TINKSINENIANGQYGSTLGSLGQTLSENGLKTAVLGNSDIVENGELIKNRNIGLVAMDQYGRIDSGNIDNINIKDDTMPYGIRTDYNKLIKETKKHYEK<br>NDVLFVDLGDTYRLDNYRLNLNEETYSNMKSKIGKNIDKYLKEVFDMIDEKDTVYISSAFPSDLDYKNKRRLSPIIKFNGEGKGVLSSATTRREGVVAN<br>LDVGADILNTFGLKNEAMVGKEYNLISKNDNIKFVSDEYEKIVSVSKIRPNVVNTFVSVVSASWVIAMVILLFKDRVHHKEKVFKILKEFIKLGFILPISFFI<br>APIFNFKTPETIVIGIVLTTLILYIIGRKLFKNDLKHMGYFAVISVLAIVIDSILGSYLMKNSIMSYDAIIGARYYGMGNEYQGIVIGSSIFGIAVLLHYNKISKW<br>LAAISCFVILITTASPIMGANVGAAISECMAFLLLVLLVFDVKLDFKKIILIGLAGVAIVFAFAAIDIISGSESHLSGFVNQILLDGPNAIIQTFSRKIQMNIKLA<br>QSSAWVNILLAGIAIIAIFIFKPSKHFRNIAKKYPFIFKGFIACIVGCFITLLVNDSGIVAAATTSIYILIPLLIISINMIIFNEK |
| A0A371J7Q2 | Romboutsia weinsteinii | QGKVIFINMTRTSLDNMLKISSLKEELDNRGHVGLMNIRGDKGTDDRRSFASMGAGGRANVSTEDLINFQETDKNEAATFKAATGQKAKAINDLTIN<br>NSIHENKDKGQYGSTLGSLGQTLSDNNLKTAVLGNSDIIENGELVKNRNIGLVAMDEHGRIDAGNIDDINVEDLSMPYGIRTDYDKLTKETKKYYNES<br>NAIFVDLGDTYRLDMYRLNLNEKTHGQMRSKIAKNIDKYLEEVFNMIGENDTVYITSAFPSDLDYKNKKRLSPIIKFSGEGKGVLASPTTRRDGIVANLD<br>VGVDILSEFGLENENMVGRKYDLVDKSDNITFLSDEYEKIVTISSIRSNVVNSFVGVVSASWIIGMFAVFFRNYIPNKKKVFTVLKEFIKLGLILPLSFLIVA<br>IFNFKTPMSISLGILATTIAFYLIGRLLFKDDIKHMGFFAVVTILAISIDCAFGTYLMKNSIMSYDAIIGARYYGMGNEYQGIAIGSAIFGLAVLLQYKKIPKW<br>LAVVSSIFILITSASPAMGANVGAAISESVAYLLFILIIFNVKLDFKKVILIGLAAAGVVFAFAAFDMISGSESHLAGFVEQILVNGPGTIIQTFSRKIQMNLK<br>LAQTSVWVNILLAGVAIIGVLIFRPSRHFRKIAREYKFIFDSFIATMVGCAITLLVNDSGIVAASTASIYILIPLIIISMNMIIFEDK |
| A0A371IWC9 | Romboutsia maritimum | EKGKVLFINMNRTNLINMQNIPFLNKELKTKGYVGLMNIRGDKGTDDRRSYATMGAGRRASVSTENYIKFENVDEKSAKAFKAATGKDAKGINNVSIN<br>RSINENELNGEYGAILGSLGQTLSDNGFKVSAIGNSDIIEDGKLEKNRNIGLIAMDKYGRIDSGNVDNINKEASNMPFKVATDYNKLIQETKKSYKETD<br>AIFVDLGDTYRLDMYRLSLNDNTYSSMKKKIYSNINTYLEEVFNMIGENDTVYITSTFPSDLDYKNKRRLSPIIKFKGDSKGILSSATTRRDGILANLDVG<br>VDVLSEFGLQNEDMVGKKYNLVDRKDNISYLLDEYQKIVSISDVRSSVVNSFVGVISASWIIAMIAVIFRNYMPHKEKVFIILKEFIKLGMIMPLSFLIVGI<br>FNFKTQMSIAIAIVIATLILYVIGRKLFKDDLKNMGFFAVITVLLVVIDAVLGTYLMKNSIMSYDPLNGARYYGVGNEYQGIIIGGAIFGSAILLNYKKIPKW<br>VTIVSSLVILITSASPAMGANVGSAISESVAYLLFILIIFDVKIDFKKVILLGIAAVVVVVSFATFDILSGTQSHLSGFVNQILTDGPGAIIETFTRKIQMNLKLA<br>QTSVWVNILLAGVGIIAVLIFKPSRHFRRIANKYPFIFKGFIASMIGCMVTLLVNDSGVVAASTTAIYILIPLVIISINMMIFNEK |

| | | |
|---|---|---|
| UPI0011DD9B03 | Terrisporobacter glycolicus | EEGKVIYISMNRANIKDLQRIPVLKEKLSKSGYIGLMNTRGDQGNDDARSYASIGAGCRANVVNDEAISFENLNDENSKIYKASTGENPKGINNLGIN<br>KSINNNIESGSYGSTLGLLGQTLSDNNKSVALLGNSDIVENNQLKKIRNTGLICMDTSGRVDSGNVDNITVKDLNMPYGLRTDYEKLTAETKTYYKSS<br>DALFVELGDTYRLDEYKLNLNENTYNKTRKQIYSYINTYLKEVFNMIGENDVVYIVSESPSTLDYNNKRRLAPVIKFSDNKKGLLESATTRRDGIIANLDL<br>GVDILSEFGLKNKAMIGRTLNNIEKDDNINYLSSQYEKIVSINNIRATVVNTFVGIVSASWVIAMLALLCKDKLPHKEKIFVILKELIKLGLIMPLTLLLSPIF<br>NFKSQLGLVSGVVIMTILLYLSGRLLFKDNDIKQMGYYAFLTIVLIVLDSVMGTPLMKNCIMSYDAIVGARYYGVGNEYEGVTIASAVFALSVLLNYKKIP<br>KWFTILLSVVILITSAFPSMGANVGGAISECIAYLLFIMLIFDVKLDFKKIMILGISAVVLVLIFAGLDLMLGMESHLGVFTQQILQEGPQAIFNTFGRKISM<br>NLKLAKSSVWVNILLVGIAVIGIFIFRPSKHMKNISNKYPMIFKGFIASMVGCLITLLVNDSGIVAAATASIYILIPILIISINMLIFNDK |
| A0A1C5M9I7 | A0A1C5M9I7_9CLOT | QGKVIFINMNRTSLASMLQISSLKEELDNRGYIGLMNIRGDKGTDDRRSLATMGAGRRANVSTESFINFRESNDNSSKVYEASTNAKAKSINDLTINRS<br>LNENAEKGEYGAVLGSLGQTLSENNLKTALIGNSDTIENENLVRNRNLGLVAMDQYGRIDAGNIDNINIEDLSMPYGIRTDYNKLLEETKNQYKESDA<br>IFVDIGDTYRLDKYRLNLNEATYSATKSKILNYIDEYLKEVFSIVNENDTIYIVSAFPSDIDYLNRRRLSPIIKLSNEGKGLLASATTRRDGIVANLDIGVDIL<br>NEFGLENEDMVGKSFTTIDKDDNTTYLLDEYEKIVSISNIRSTIINTFVGVVSVSWVIAMFAVLFRNYIPNKEKVFKVLKEFIKLGFILPLAFLVVGIFNFKT<br>QVSISLGITGTTILLYILGRILFKDDIKHMGFFAGITILAISMDCVFGTYLMKNSIMSYDAIIGARYYGMGNEYQGIAIGSAIFALAVLLQYRKIPKWLAAVAS<br>VLTLLTTASPSMGANVGAAISECVAYLVFLLIIFDVKLDFKKVFLILLAAAGVVFAFAAFDIISGSESHLSGFVKQILANGPITIIQTFSRKIQMNLKLAKTTV<br>WVNILLVGVAIVGIMIFRPSKHFRNLANKYKYIFDSFIATLVGCFITLLVNDSGIVAAATACIYILIPLIIISINMIIFEDE |
| UPI001D16E2B1 | Terrisporobacter mayombei | EKGKVIYISMNRANVNDLQRIPVLKEKLSKSGYIGLMNTRGDQGNDDARSYASIGAGCRANVVDNEVISFENLTDENGKIYKASTGEDPKGINNLFVN<br>KSINNNIESGSYGSTLGLLGQTLSDNNKKVALLGNSDIVENNQLIKIRNAGLICMDTLGRVDSGNVDDITVKDLNMPYGLRTDYDKLTADTKTYYESS<br>DALFVELGDTYRLDEYKLNLNENTYNKTRKQIYSYINTYLKEVFNMIDENDVVYIVSEFPSTLDYNNKRRLAPVIKFSDNKKGLLESATTRRDGIIANLDL<br>GVDILNEFGLKNEAMIGRTLNNIEKDDNINYLSSQYEKIVSINDIRSTVVNTFVGIVSASWVIAMLALLCKDKLPHKEKIFVILKELIKLGLIMPLTLLLSPIF<br>NFKSQLGLVSGVVIMTILLYLSGRLLFKDNDIKQMGYYAFLTIALIVLDSVIGTPLMKNCIMSYDAIVGARYYGVGNEYEGVTIASAVFALSVLLNYKKMP<br>KWFTIVLSLVILITSAFPSMGANVGGAISECIAYLLFIMLIFDVKLDFKKIIILGLSAVVLVLIFAGLDLMLGMESHLGVFTQQILQEGPQAIFNTFGRKISM<br>NLKLAKSSVWVNILLVGIAVIGIFIFRPSKHMKNISNKYPMIFKGFIASMVGCLITLLVNDSGIVAAATASIYILIPILVISINMIIFNDE |
| A0A943AQC7 | A0A943AQC7_9FIRM | ESGKVIFINMNRTTIKSMQDIPSLKAELEKRGYMGLMNIRGDKGTDDKRSLASMGAGGRANLASDSYINFKEATKENATIYKSSTGKTPKKINDLSINQ<br>SLNENEANGQYGSTLGSLGQTLSDNNFKVAVLGNSDTVENGELKENRNICLIAMDNYGRVADGNIEDINIEDDTMPFGIRADYDKLTKETKSLYENN<br>DALFVDLGDTYRLDQYKGFLNEHTYSKMKKTIHNNISKYLESVFNMVGDNDVVYIASSFPSKLAYKNKERLSPIIKFKGNEKGLLSSSTTRRDGIVANID<br>VGVDILNEFGLENKSMVGRAYSLIQKDDNVDFLSYELEKMATISNIRSTVVNTFVGVVSVSWVIGMVAILFRNRIPNKDKVFNVIKEFIKLGIIMPLVFLL<br>APIFNFKTPVSMTIGIIITTLALYLLGRVLFKDDLKQMGFFALITILVIVIDCIFGTYLMKNNIMSYDAIIGARYYGVGNEYEGVSIASPIFAFAILLNyNKKLPK<br>WSIIIASIVILITSAYPTMGANVGGAISQTAAYLLFIMLIFDVKLDLKKVILIGISVVAVVGAFAFFDIVSGSESHLGLFVQQIFLNGPSAIIQTFARKIGMNVK<br>LAQTSVWVNILLAGIFIIGIFIIKPPRQFRMIAKRYPMIFKGFIASMVGCIVTLLVNDSGIVAASTASIYILIPLIIISINMLVPENK |
| A0A0B3W2P0 | Terrisporobacter othiniensis | EEGKVIYISMNRTNINDLQKIPVLQEKLKNSGYIGLMNTRGDQGNDDARSYATIGAGTRANVVNNEVISFENLTDENGKLYKTATGEEPKGINNISINK<br>SINNNAEKGSYGSTLGLLGQTLSDNNKKVALLGNSDIVENNQLIKIRNTGLIAMDTLGRVESGNVDNISVKDLSMPYGLRTDYNKLITDTKTYYESNDA<br>LFIELGDTYRLDEYKSNLNENTYNKTKKQIYSHINTYLKEVFNMVNENDVIYIVSEFPSTLDYNNKRRLAPVIKFSDNKKGLLESATTRRGGIIANLDLGV<br>DILNEFGLKNEAMIGRALSSIEKDDNIDFLSGEYKKIVSINNIRSTVVNTFVGIVSTSWVIAMLALLFKDKLPNKEKVFIVLKELIKLGLIMPLTLLLSPIFNF<br>KSQLGLVSGVVIMTIILYLSGRILFKNNDIKQMGYYALLTILLIVIDSVIGTYFMKNCIMSYDAIVGARYYGIGNEYEGVTIASAVFALSVLLNYKKMPKWLT<br>IVLSVIILITSAFPSMGANVGGAISECVAYLLFIMLIFDVKLDLKKIMILGLAAVVLVLVFAGLDLVLGMESHLGVFTQQIIQEGPQAIFNTFGRKISMNLKL<br>AKSSVWVNILLVGIAVIGIFIFRPSKHMKNISNKYPMIFKGFIASMVGCLITLLVNDSGIVAAATASIYILIPILIISINMIIFNDK |

| | | |
|---|---|---|
| A0A1V1I2V3 | Romboutsia ilealis | ESGKVIFINMNRTTLESMQDIHSLKTELEKRGYMGLMNIRGDQGTDDNRSLASMGAGGRVNLASDSYINFKEATKENTIIYKAVTGQSSKKINDLAIN<br>QSLNENEAKGQYGSTLGSLGQTLSDNNFKVAVLGNSDTVENDEVKENRNICLIAMDNYGRVEDGNIDDINIEDDTMPFGIRTDYDKLTKETKALYEN<br>NDALFVDLGDTYRLDQYKGYLNEKTYSKMKKTIYNNISQYLKSVFDMVGENDVVYIASSFPSKLAYKNKERLSPIVKFKGNKKGLLSSSTTRRDGIVANI<br>DVGVDILNEFGLKNEAMVGRAYSLIQQDDNLDFLSHELEKMATVSSIRANVVNTFVGVVSVSWVIGMVAILFRNRIPNKEKVFGVLKEFIKLGIIMPLV<br>FLLAPIFNFKTPVSMTIGIIITTLVLYLLGRLLFKDDLKQMGFFALVTIVVIVVDCMSGSYLMKNSIMSYDAIVGARYYGVGNEYEGVSIASPIFAFAILLNy<br>NKKLPKWSIVIASILILITSAYPTMGANVGGAISQTAAYLLFIMLIFDIKLDFKKVILICISVVGVVGIFAFLDIVSGSESHLGLFVQQILLNGPSTIIQTFARKI<br>SMNVKLAQTSVWVNILLAGIFIIGIFIIKPPKQIRMIAKRYPIIFKGFIASMIGCMVTLLVNDSGIVAASTASIYVLIPLIIISINMLVLENK |
| UPI001D16B73A | Terrisporobacter mayombei | EEGKVIYISMNRSNINDLQRIPVLNEKLNKSGYIGLMNTRGDQGNDDARSYASIGAGSRANVISNGPISFVNLTEESGLIYKSATGENPKEINNLDINA<br>SININVEQGSYGSTLGVLGQTLSDHNKKMALLGNSDIVENNNLVKIRNAGLICMDTSGRVYSGNVDNINTKDLSMPYGIRTDYEKLIKETKSYYKDND<br>ALFIELGDTYRLDKYKSNLNEKTYNKIKKNIYTNVNTYLSEVFNMIGKNDVVYIVSESQSDLDYNNKRRLSPVIKFSDYKKGLLSSATTRRDGIIANLDLG<br>VDILNEFGLSNDIMIGRTFDYIEKDDNIKYLSNQYEKIVSINNIRSTLVNIFVGIVSVSGVMAMLALLFKDKLPYKDKVFIVLKEFIKLGIIMPLALLSAPIFN<br>AKTQMGIIISVIAMTILIYLSGRLLFKHSDIKQMSYYALLTIGLVVFDSIIGTFFMKYSVMSYDAIVGARYYGVGNEYQGVIIGSAIFGLAVLLDNKRIHKWL<br>VGALFVVILITTASPFMGANVGAAISECVAFLIFMMLIYGVKLDLKKIIMVGLAAFLLVLGFAGLDLLMGTESHLGAFVMEILREGPSAIILTFSRKISMNLK<br>LARSTVWVNILLTGIAVIGIFIFRPSRHMKNISNKYPMIFKGFIASMVGCLIALLVNDSGIVAAATASIYILIPILIISINMIIFNDK |
| UPI00047F4213 | Metaclostridioides mangenotii | EEKGKVIFINMNRTNLINMRKIPILENEIDRRGYLAYMNIRGDKGTNDRRSYASMGAGGRANVSKiEDPIIFNKTNKEDAEKFKAVTGKDAKEINYLNVN<br>RSINENYENGEFGSSLGVLGQTLSDNNLKTAVIGNSDIVDNGKLLENRDLCLMAMDQYGRIDMGNIDDINKKDLSMPFGISTDYDKLLNETKKYHS<br>QSDVVYVDLGDTYRLDLYRNSLNDNSFNEMRDNIEKKIDDYLTQVFKMAGKNDIIYIASGFPSDIDFKNKRRLTPIVKFKGDGKGLISSATTRRNGITA<br>NLDVGVDILNEFGLKNKLMVGKPFNLIDKNNNVEYLLGEYEKIVSITNVRATIINTFVSVISISWIIATVAILIRNRIPNKEKVFNILKELIKLGILMPLSFMIAT<br>LFNFKSQIGILFGIFVVASILYLLGRVLFKDDLKHMGFFSGATILLIVLDSILGTYMMKNNIMSYDAIVGARYYGIGNEYGGVIIGSSIFAISVLMSYKEIPK<br>WTVILASLVILITCAYPSMGASVGGAISECVAYLLFIFLIFNMKIDLKRILILGFSAVAVVAIFAYLDIVSGAESHLSGFVQQIMLKGPGQIIQTFTRKIGMNI<br>KIAQTSVWVNILLAGIGIMAVLIFKPSKHFKRVMIEYPIIFKGLVSSVVGCVVALLVNDSGIVAAATASIYIFIPVVIISINMIIFNESN |
| A0A6H1X129 | A0A6H1X129_9FIRM | EDKKVIFINMDRTNLENMMKMPFLKEKLSKEGYIGLMNIKGDRGTDDKRSYATIGAGRRANVANDEYINFAQRTDENAKVYESAVGEKAKAINDLTIN<br>MSLSANKTGEYGAVLGSLGQTLSENNIKTAVLGNADIIVGDSITKNRNIALIAMDNYGRIPAGNIEDINIEDSSMPFGMRTDYDKLLNETKKYYNESDAI<br>FIELGDTYRLDKYKKNLNSNTYDKMRDSIYSNINNYIESVFKLVDENDVVYITSTFPSDFDYKNKRRLSPVVKFSKEynGKGILVSATTRRDGIIANLDLG<br>VDILNEYGLSNEHMVGRVINKIEKDDNIDYLSHEFDKIVSISKVRANIVNTFVIIIAASWIIGMLAVFARKKIPYKKKVFAIIKEFIKLGIIMPLAFLVAPMVNY<br>RTEFGIGSSIIITTIVLYIIGRILFKDDIKNMGFFSLITIILIAIDSIFGTYLMQNNIMSYDAMIGARYYGIGNEYEGITIASAVFGFAVLLQYKKIPKWLTIISLFVIL<br>ITSAYPSMGANVGGAISEIVAYLLFIMLIFDVKLDLKKVVILGLSAVGIVSIFAVLDLVTGSQSHLGGFVSQILVEGPSVIINTFGRKIQMNLKLAQTSIWV<br>NILLAGIVIIGVLIFRPSEHFKRIAKKYPVVFKGFIASMVGCVVTLLVNDSGIVAASTASIYILIPIIIMSINMIIFE |
| UPI001D0F7B11 | unclassified Clostridioides | GKVIFIDMNRTSMNNMLRIKSLREELENRGYIGLMNIRGDKGSDDRRSYASMGAGGRANVANEEDINFESSSKDRNIVFESVTGKKAKGINDLTINM<br>SINENLNFGEYGSTLGSLGQSLSENELKTSILGNADIVEGGQLVKNRNLCLTAMDEYGRIPYGNVDNINKKDLSMPYGISTDYTKLVAETKEAYKNSD<br>VVFVELGDTYRLDLYKPNLNENTYKNMKDKIEKNIDEYLKNIFSMVGENDTVYIASAFPSDLDYKNKRRLSPIIKLSGNGKGLLSSSTTRREGIVTNLDV<br>GVDILDSFDLKNPNMVGRKYELINRDDNKEYLMDEYQKIVSISSIRSVILNGFVNVVFLSWIVAMIAILFRKHISKKHKETTyfILKELLKIGIVMPLSFMITP<br>IMNFKTPVAISLGIIIITLTIYLISKVLIKNDLKNILFFTGLTILIMVIDSALGAYLMKSNVMSYDCIIGARYYGVGNEYQGVAIGSAIFTFAILLTYKKIPKWSVIV<br>FSLAILITSASPIMGANVGSAISECVAFLLFILLIYDVKIDFKKVILLGIAVLLVLGVFVTIDIVSGSNSHLGMFVKDIYFNGPGEIIQTFSRKIEMNLKLAQTS<br>AWVNILLAGIGVILVLMLTQIRYFKQLINEYPIVFKGFIASVVGCFVTLLVNDSGIVSSATAFIYVIVPMIILSINMTALKE |
| UPI00129DAE47 | Romboutsia faecis | DESKKTIFINMDRTSLESMLEIPAIKDELVKRGYIGLMNIRGDKGTDDKRSYASIGAGRKANVAADEYINFVENTEMNSKLYESSVGEKPKAINDITINMS<br>LAASKQGEYGAILGTLGQTLSDNNINYSVLGNADMQVNDQLVKNRNIGLLAMDNYGRIPSGNIEDINIEDSSMPFGIRTDYDKLITETNKYYNNSDAL<br>FIELGDTYRLDNYRMNLNSKTLGQMRTAIYSNIDKYLREVFSMVDEDDVIYITSVFPSESDYKNKRRLSPVVKLSGDGKGILASSTTRRDGIISNIDLGV<br>DILNEFGLSNELMSGRVLSFIDKDDNIGYLSHEYDKIVSISKTRSDIINTFVIIIAASWVIGMLAVFARNKIPYKEKVFTVIKELIKLGIIMPLAFLVAPLVNYR<br>TEFSIGTTIIISTLVLYVLGKVLFKDDIKNMGFFAFLTIMIIAIDSVFGTYLMQNNIMSYDALIGARYYGVGNEYEGVTIASAIFGFSVLLNYKKIPKWLAAVA<br>LFIILITSAHPAMGANVGGAISESIGYLLFVMLIFDIKIDFKKVIILGLSAIAVVGVFALFDLVSGSQSHLGAFVNQILSEGPSAIISTFARKIHMNIKLAHTSV<br>WVNILLAGIAVIGVLIFRPSEHFKRIALRYPIIFKGFIASMVGCVVALLVNDSGIVAASTASIYILIPIIIMSINMIIFEDE |

| | | |
|---|---|---|
| UPI00035C6455 | Terrisporobacter glycolicus ATCC 14880 = DSM 1288 | EEGKVIYISMNRSNINDLQGIPVLKEKLSKSGYIGLMNTRGDQGNDDERSYASIGAGGRANVISNGPISFENLTKENGLIYKSATGKNPKEINNLNINA<br>SININIEQSSYGSTLGVLGQTLSDNNKKMALLGNSDIIENNNVIKIRNIGLVCMDTFGRVYSGNVDNINIKDLSMPYGIRTDYEKLIKETKSYYKDNDAL<br>FIELGDTYRLDKYKSNLNEKTYNKIKTNIYTNVNNYLSEVLNMVNENDVVYIVSEFQSNLDYNNKRRLSPVIKFSGHKKGLLSSATTRRDGIIANLDLGV<br>DILKEFGLSNDIMIGKTFDYIEKDDNIKYLSNQYEKIVSINNIRSTLVNIFVGIVSVSGLMAMIALLFKDKLPYKDKVFIVLKEFIKLGMVMPLALLSSPIFNA<br>KTQMGIIISVIAMTILIYLSGRLLFKHSDIKQMSYYALLTVGLVVFDSIIGTFFMKYSVMSYDAIVGARYYGVGNEYQGVIIGSSIFGLAVLLNNKRMHKWL<br>VGALFIVILITTAAPFMGANVGAAISECIAFLIFMMLIYDVKLDLKKIIIVVLTAFLLVLGFAGLDILMGTESHLGAFVMEILREGPSAIILTFSRKISMNLKLAK<br>STVWVNILLTGIAVIGIFIFRPSRHMKNISNKYPMIFKGFIASIVGCLITLLVNDSGIVAAATASIYILIPILIISINMMIFNNK |
| UPI0006BC02E8 | Clostridioides difficile | GKVVFIDMNRTSMSNMLRIKSLREELDNRGYIGLMNIRGDKGSDDRRSYASMGAGGRANVANEEDINFESSSKDRDIVFKAATGKNAKGINNLTINK<br>SINENLNFGEYGSVLGSLGQSLSDNGLKTSVLGNADIIENGQLIKNRNLCLVAMDEYGRIPNGNVDNINKKDLSMPYGISTDYNKLITETREAYKNND<br>VVFVELGDTYRLDLYKPNLTEETYASIKNNIEKNIDVYLKNVFSMVGENDTVYISSAFPSDLDYKNKRRLSPIIKLNGNGKGLLSSSTTRREGIVTNLDVG<br>AEILNNFDIKNQNMVGKKYELINRDDNKEYLMDEYQKIVSVSSIRSIILNGFVSIVFLSWIVAMILILFRKHISknHKETIFFILKEFLKIGIVMPLSFMITPIM<br>NFKTPVAISLGIIIITLTIYLISKVLIKNDLKNMLFFTGLTILIMVIDSAFGSYLMKSNVMSYDCIIGARYYGVGNEYQGVAIGSAILTFAVLLTYKKIPKWSVIV<br>FSLVILITSASPIMGANVGSAISECVAFLLFILLIYNVKIDLKKVVLLGIAVLLVLGVFVAIDMISGSNSHLGMFVKEIYTNGPGEIIQTFSRKIEMNLKLAQT<br>SAWVNILLTGIGVILVLMINQIRYFKQLIDEYPIVFKGFIASIAGCFVTLLVNDSGIVSSATAFIYVIVPMIILSINLVALKE |
| UPI0010B6C248 | Clostridioides difficile | RGKVIFIDMNRTSMSNMLRIKSLREELDNRGYIGLMNIRGDKGSDDRRSYASMGAGGRANVANEEDINFESSSKDRNIVFESATGKSAKGINNLTINK<br>SVNENLNFGEYGSVLGSLGQSLSDNGLKASVLGNSDIIENGQLIKNRNLCLTAMDEYGRIPNGNVDTINKKDLSMPYGISTDYDKLIVETKEAYKNND<br>VVFVELGDTYRLDLYKPNLNEKTYENMKNKIEKNIDVYLKNIFSMVGENDTIYIASAFPSDLDYKNKRRLSPIVKLNGVGKGLLSSSTTRREGIVTNLDV<br>GVEILDNFNIKNQNMVGRKYELINRDDNKEFLMDEYQKIVSISSIRSTILNGFVSIVFLSWIVAMIAILFRKhiSKKYKETTFFILKELLKIGIVMPLSFMITPI<br>MNFKTPLAISLGIIIITLTIYLISKVLIKNDLKNMLFFTGLTIVIMVIDAGFGSYLMKSNVMSYDCIIGARYYGVGNEYQGVAIGSAIFTFAILLTYKNIPKWSVI<br>VFSLLILITSASPIMGANVGSAISECVAFLLFILLIYNVKIDFKKIVLLGIAVLFVLGAFVAIDMILGSNSHLGMFVKEIYFNGPGEIIQTFSRKIEMNLKLAQT<br>SAWVNILLTGIGVILALMINQIRYFKQLMDEYPIVFKGFIASIAGCLVTLLVNDSGIVSSATAFIYIIVPIITLSVNLTALKE |
| UPI001D0FD3AE | unclassified Clostridioides | GKVIFIDMNRTSMNNMLRIKSLREELEDRGYIGLMNIRGDKGSDDRRSYASMGAGGRANVANEEDINFESSSKDRNIVFESATGKKAKGINDLTINM<br>SINENLNFGEYGSTLGSLGQSLSENKLKTSVLGNADIVENGKLVKNRNLCLTAMDEYGRISYGNVDNINKKDLSMPYGISTDYTKLVAETKEAYKNSD<br>VVFVELGDTYRLDLYKSNLNEKTYKNMKDKIEKNIDEYLKNIFSMVGENDTVYIASAFPSDLDYKNKRRLSPIIKLNGNGKGMLSSSTTRREGIVTNLDV<br>GAEILNNFDIKNQNMVGKKYELINRDDNKEYLMDEYQKIVSVSSIRSTILNGFVSIVFLSWIVAMILILFRKHISknHKETIFFILKEFLKIGIVMPLSFMITPI<br>MNFKTPVAISLGIIIITLTIYLISKVLIKNDLKNMLFFTGLTILIMVIDSAFGSYLMKSNVMSYDCIIGARYYGVGNEYQGVAIGSAIFTFAILLTYKKIPKWSVI<br>VFSLVILITSASPAMGANVGTAISECVAFLLFILLIYNVKIDFKKVVLLGVAVLLVLGAFVAIDMISGSNSHLGMFVKEVYFNGPGEIIQTFSRKIEMNLKLA<br>QTSAWVNILLTGIGVILVLMINQIRYFKQLIDEYPIIFKGFIASIAGCFVTLLVNDSGIVSSATAFIYVIVAMIILSINLIVLKE |
| UPI001D0C8C8A | Clostridioides sp. ES-S-0108-01 | GKVIFIDMNRTSMSNMLKIKSLREELDGRGYIGLMNIRGDKGNDDRRSYASMGAGGRANVANEEDINFESSSKDRDIAFKSATGKSAKGINNLTINKS<br>INENLNFGEYGSVLGSLGQSLSDNGLKVSVLGNADIIENGQLIKNRNLCLVAMDEYGRIPNGNVDNINKKDLSMPYGISTDYNKLITETKEAYKNNDV<br>VFVELGDTYRLDLYKPNLTEKTYVNMKNSIEKNIDVYLKNVFSMVGENDTVYISSAFPSDLDYKNKRRLSPIIKLNGDGKGLLSSSTTRREGIVTNLDVG<br>VEILDNFNIKNQNMVGRKYEPISRDDNKEYLMDEYQKIVSISSIRSTILNGFVSIVFLSWIVAMIAILFRKHISknYKETTFFILKELLKIGIVMPLSFMITPIM<br>NFKTPLAISLGIIIITLTIYLISKVLIKNDLKNMLFFTGLTILIVVIDSALGSYLMKSNVMSYDCIIGARYYGVGNEYQGVAIGSSIFTFVILLTYKNIPKWSVIVF<br>SLAILITSASPIMGANVGSAISECVAFLLFILLIYNIKIDFKKIVLLGIAVLFVLGVFVAIDMISGSNSHLGVFVKEIYFNGPGEIIQTFSRKIEMNLKLAQTST<br>WVNILLTGIGVILVLMINQIRYFKQLLDEYPIVFKGFIASIAGCLVTLLVNDSGIVSSATAFIYVIVPMIILSINLIALKE |
| UPI001D0F84C4 | Clostridioides sp. ZZV15-6383 | GKVIFIDMNRTSMSNMLRIKSLREELDNRGYIGLMNIRGDKGSDDRRSYASMGAGGRANVANEEDINFEGSSKDKNINFESATGKSAKGINNLTINK<br>SINENLNFGEYGSVLGSLGQSLSDNGLKASVLGNSDIIENGQLIKNRNLCLTAMDEYGRIPNGNVDNINKKDLSMPYGISTDYNKLIVETKESYKNND<br>VVFIEIGDTYRLDLYKPNLNEKTYESMKDKIEKNIDVYLKNLFSMVGENDTVYISSAFPSDLDYKNKRRLSPIIKLNGNGKGLLSSSTTRREGIVTNLDVG<br>VEILQNFDIKNQNMVGKKYELINRDDNKEYLMDEYQKIVSISSIRSTILNGFVSIVFLSWIVAMIAILLRKHISKNHRETTffILKELLKIGIVMPLSFMITPIM<br>NFKTPLAISLGMIIITLTIYLISKVLIKNDLKNMLFFTGLTILILVIDSGFGSYLMKSNVMSYDCIIGARYYGVGNEYQGVAIGSAIFTFAILLTYKNVPKWSVII<br>FSLVILITSASPIMGANVGSAISECVAFLLFILLIYNVKIDFKKVVLLGIAVLLVLGVFVAIDMISGSNSHLGMFVKEIYFNGPGEIIQTFSRKIEMNLKLAQT<br>SAWVNILLTGIGVILVLMINQIRYFKQLIDEYPLIFKGFIASIVGCFVTLLVNDSGIVSSATAFIYIIVPMIILSINLT |

| | | |
|---|---|---|
| UPI00202CB53B | Paraclostridium ghonii | DDKGKIIVINMNRTSFEDLLSIDSLSRKMEKEGYIGLMNIKGDKGTDDRRSYASVGAGGKVTLPDQSLINFEQVNEDNAKAYKAKTGHNPKKINDMKI<br>NHSLSENLENGSYGSVLGSLGQTLSNHKLKSSVIGNADRVNNGELVLNRNIALMAMDNIGRVDDGNVDTINIKDNTMPYGIRTDYDKLKSETKKFYD<br>NSDVLFVELGDTYRLDDYKNYLNDKTYNDMKKNIYKNIDSYVEEVFKMVDENDTIYIMSAFPSKEDYKNQKRLSPIIKFEGSQKGVLKSATTRRAGVVG<br>NIDIGVDILSEFGLENEEMVGKKFESIDKDNNIEFINHEFDKMVSMYQVRTLIINTTVGIVAAAWIIcAILLVFCRDKLPKKKLIFNTLKEFIKLGILIPLAFIV<br>SPIFNFSTQASIALGIVGTLIVLYLIGKIFFKDDMKYMAFFSILTIIVVVVDSIFGTYLMKNNIMSYDAIIGARYYGIGNEYEGITIGCTIFAMSVLVNYKKIPK<br>WAVVVVSLIILITSAYPSMGANVGGAISESVAYTLFILLAFDVQMDFKKSILLSLVPVMIVFIFAVLDIVSKSQSHLSGFVNQILVNGPIAIIQTFTRKIQMN<br>VEIATSNILIVVALVVAGYMTKFIFKPVNHFKKLSDKYPYIFKGFVATTVGCIVTLIFNDSGIVAAGTASIYILVPILIMSVNMIVFDKE |
| UPI001F1EB3EF | Terrisporobacter glycolicus | EGKVIYISMNRSNINDLQRIPALKEKLSKSGYIGLMNTRGDQGNDDARSYASIGAGGRANVISNGPISFENLNKDNGLMYKSAIGKNPKGINNLDINA<br>SINVNIEKGSYGSTLGVLGQTLSDANKKIALLGNSDIVKNNNIVKIRNAGLICMDNFGRVNTGNVDNINTKDLSMPYGIKTDYEKLIKETKSYYKGNDA<br>LFIELGDTYRLDEYKSNLNEKTYKDIKKNIYANINNYLSEVFNMVGENDVVYIVSEFQSKLDYNNKRRLSPVVKFSGSKKGLLSSATTRRDGIIANLDLG<br>VDILKEFGLSSNIMIGRAFDHVEKSDNISYLSNQYEKIVSINSVRSTLVNVFVGIVSASGVIAMLALLFKNKLSYKDKVFTVLKEFIKLGLIMPLALLSAPIF<br>NAKTQTGIIISVVVISILIYLSGRLLFKNSDIKQMSYYALLTVGLVVFDSIIGTFFMKYSVMSYDAIVGARYYGVGNEYQGVIIGSAIFGLAVLLNKERIHKW<br>LVGVLFIVLLITTAAPFMGANVGAAISECVAFLIFMMLIYGVKLDLKKIIIVGFAALLLVLGFAGLDILMGTESHLGAFVMEILQKGPSVIILTFSRKISMNLK<br>LARSTVWVNILLTGIAVVGIFIFRPSKHMKNISNKYPMIFKGFIASMVGCLVTLLVNDSGIVAAATASIYILIPILIISINMIIFNDK |
| UPI000A178399 | Paraclostridium bifermentans | DDKGKAIVIDLNRTGLEDMLSIDSLSKKMEKEGYIGLMNIRGDRGTDDRRSYAAVGAGGKVTLPNQSLINFEEVNKGNAKAYKAETGHNPKKINDMTI<br>NHSLSENLENGSYGSVLGSLGQTLSNHKLRASVIGNADRVNNGELIKNRNIALMAMDNIGRVDDGNVDNINIKDNTMPYGIRTDYNKLKSETKKFY<br>DNSDVLFVELGDTYRLDDYKNYLNEKTYNNMKKNIYKNIDDYLQEVFKMVDENDTIYIMSAFPSNADYKNQKRLSPVIKFEGSQKGVLKSATTRREGV<br>VGNLDIGVDILAKFGLKNEKMVGRAFEEVNKEDNISFINHEFDKMVSMLGVRTPIVNTTVGIVAAAWIICrVLLIFFRKKLPKKELVFSILKEFIKLGVVIPL<br>AFIVSPIFNFATPTSITLGIFGTLLVLYLIGKIFFKDDIKYMAFFSIITIIVIVVDSIFGTHLMQNNIMSYDAIIGARYYGVGNEYEGITIGCAIFAMSVLVNYKKIP<br>KWIAVVASLIILITSAYPSMGANVGGAISESVAYTLFLLLAFDVKMDLKKSILLSLVPVIIVCIFAALDVVSKSQSHLSGFVNQISVNGPIAIIQTFTRKIQMN<br>VDIATSNILIVVALVVAGYMTKFIFKPANHFKKLSEKYPYIFKGFVATTVGCIVTLIFNDSGIVAAGTASIYILIPILVMSVNMMIFDKE |
| UPI001C1165DC | Intestinibacter bartlettii | GKVIYVSLNRTNLEHLLEIPIIKEKTSNSGYVALMTTRGDGSNTDQKSFASIGSGVRANVLDQDYTIFESLDKDGQAQFKAITGVKPWAINNLRINRDL<br>NQNSDYGSYGATLGTLGTTLNENGLKVALLGNSDIIKNGELKKIRNLALTCMDQYGRVESGNVDDINIDDNAMPYGISTDYKKLKEETKKYYENNDIT<br>FVELGDTYRLDEFRKNLNENTYASMKKRIYQRVNDYLKEVFDMVGENDTVYIVSEFQSNLDHLNKRRLAPVMKFTKDGYGLLESATTRNSGIVSNIDI<br>GVDILNEFGLKNELMIGKPFKTVQAENGIEKLLNDFKRIVSINSIRASVVNTYVGIVSASWVIAMIALLFKNKIPKKEKglgrdqIFKILKEFIKLGLIMPLAFLT<br>APIFQFKTQTGLVAGVLIATIIYYLIGSKLFKNDDIKQMGFYALLTIGLIIVDCVIGTPLMKTNIMSYDAINGARYYGVGNEYEGVTIASAVFALAVLLNYKKI<br>PKWLAVILSLVILVTSAMPSMGANVGGAISECVAYLLFILLIFDVKLDLKKIIMIGVATVLLVVVFAGLDFLMGTESHLAVFTNQIIQEGPGAIINTFGRKIQ<br>MNMKLAKSSVWVNILIVSIVIIGALIFKPSGQFKKIHDKYPMIFKGFMASFVGCIVTLLVNDSGIVAASTAAIYILIPLIIISINM |
| UPI0018AB3675 | Clostridium sp. 1001270J_160509_D11 | QQGKIIYISLNRTSLEHLLEIPIIQKKTSDSGYVALMTTRGDGSNSDQKSFASIGAGVRANVLDQDYTIFESLDKDGKEQFKAVTGATPWAINNLRINRD<br>LNQNSDYGSYGATLGTLGTTLNENGLKVALLGNSDIIKNGELKKIRNLALTCMDKYGRVKSGNVDDINKNDNTMPYGISTDYEKLKKETKKYYENSDI<br>TFVELGDTYRLDEFRKNLNENTYKSMQDEIYKRVSDYLTEVFDMVGENDTVYIVSEFQSNLDNLNKRRLAPVMKFTKDGYGLLESATTRNNGIVSNIDI<br>GVDILNEFGIKNELMIGKPFKTVQAENGIEKLTNDFKRIVSINSIRASVVNTYVGIVSASWVIAMIALLFKNKIPKKEKglgrdqIFKILKEFIKLGLIMPLAFLT<br>APIFQFKTQTGLVTGVLIATIIYYVIGSTVFKKDDIKQMGFYAFLTIGLIILDCIIGTPLMKSNIMSYDAINGARYYGVGNEYEGVTIASAVFALAVLLNYKKIP<br>KWLAVVLSIIILVTSAMPSMGANVGGAISECVAYLLFILLIFDVKLDLKKIIMIGVATVLVVVAFAGLDFLMGSESHLSVFTNQIIQEGPAAIINTFGRKIQM<br>NMKLAKSSVWVNILIVSIVIIGALIFKPSGYFKKIHDRYPMIFKGFMASFVGCIVTLLVNDSGIVAASTAAIYILIPLIIISINMSLFDKN |

| | | |
|---|---|---|
| UPI001655184F | Paraclostridium tenue | EKGKVILINMNKTSLEDMLSIDALNKKLNKEGYIGLMNIRGDRGTDDRRSYAAIGAGGKVTMPSQSLINFEEVNKDNGVAYKAETGHNPKKINDMTIN<br>YSLSENLENGSYGSKLGSLGQTLADKKIKTSVIGNADKVNNGELLKNRNIALMAMDNIGRVESGNVDNINIKDNTMPYGIRTDYDKLKNETKKFYDDS<br>EVVFIELGDTYRLDEYKNYLNDKTYKNMKNKIYKNIDSYLKEVFKTVNENDTVYIMSTFPSKKDYKEQKRLSPIIKFEGSGKGVLRSATTRRDGVVGNIDI<br>GVDILSKFGIENPQMVGKKLIKVDKKDNISFINHEFEKMVSMIGVRSTIVNTSVGIVAVAGVIAtILLLFFRDKLPKKEMIFSILKEFLKLGILIPLVFIVSPIFN<br>FSTKGSITMGILGTLVAFYIIGKKLFKDDIAYMGFFSILTIIVIVIDSMFGTHLMQNNIMSYDAIIGARYYGIGNEYEGITIGCAIFALSVLVNYKKIPNWFVVV<br>LALIILITSAYPSMGANVGGSISESVAYVLFILLIFDAKMDLKKYILLALVPVIIVFLFAALDIVSKSQSHLSVFVNQIMVDGPGAIIQTFVRKIQMNIDIATSN<br>VFIIIVFVIVGFISSFIFKPSRQFKKISDKYPYIFKGFVASTVGCIVTLLVNDSGIVAAGTTSIYILIPLLIMSINMMIFNKE |
| A0A1C5XMB5 | uncultured Clostridium sp. | NKGKVIYLSLNRTSLENLLEIPIIKEKTSNSGYVALMTTKGDGSNTDQKSFASIGSGVRANVLDEDYVKFETLNENEKTQFEAATGVKSWAIGNIKINRNI<br>NQNTDNGGYGATLGMLGTTLNKNNLKVALLGNSDIKKNDDLKKIRNLALVCMDENGRVQSGNVDNINKKDDTMPFGISTDYDKLKSETKKYYENN<br>DAIFVELGDTYRLDEYKNNLNENTYKSMKKKIYQNINDYLKEVFEMAGKNDTIYIVSEFESNLDYKTKRRLSPVIKFEKDGYGLLESSTTRNSGIVSNIDI<br>GVDIINKFGLKNELMIGKVFKNVQDENGIEKVLKDYERIVSINTIRASVVNSYVGIVSASWVIAMIAILFKNKIPKKEKglgrdqIFKVLKELIKLGLIMPLAFLT<br>APVFQFKTPAGLVSGVLIVTIIYYLIGYFLFRKDDIKQMGFYAFLTIGLIIIDCVIGTPLMKSGIMSYDAINGARYYGVGNEYEGVTIASAVFALAVLLNYKKI<br>SKWLAVVLSLVILITSAMPSMGANVGGAISECVAYLLFILLIFDVKLDLKKIIMIGVATVLVVVAFAGLDFLMGTESHLAVFTNQIIQEGPAAIFNTFGRKIQ<br>MNLKLAKSSVWVNILIVSIIIIGALIFKPSGYFKKIHDKYPMIFKGFMASFVGCIVTLLVNDSGIVAASTAAIYILIPIIIIGINMIIFDKE |
| A0A446IDW2 | Paraclostridium sordellii 8483 | EEKGKVILINMNRTSLDDMLSIQALSKKVENEGYIGLMNIRGDRGTDDRRSYAAIGAGGRVTMPSQSLINFEELNKDNGPAYKAETGHNPKKINDMTI<br>NYSLSENLENGSYGSKLGSLGQTLADKKIKTSVIGNADKVNNGELLKNRNIALMAMDNIGRVENGNVDNINIKDNTMPYGISTDYNKLKSDTKKFYD<br>NSDVLFVELGDTYRLDEYKDYLNEKTYKEMKLRIYNKINSYLEEVFKMANKNDTIYITSTFPSNEDYKNQKRLSPIIKFDGSSKGVLKSATTRREGVVGNI<br>DIGVDILSKFGLVNENMLGKKFIDVKKTNNVEFINHEFEKMVSMLGVRSTIVNTSVGIVAAAWVICtILLVFFRNKIPKKEVVFKVLKEIVKLGILLPLAFIL<br>APIFNLSTKAGITMGIVGTLIGLYIIGKIFFKKDMTYLMFFSLMTIIVIVIDSIFGTYLMQNNIMSYDAIIGARYYGVGNEYEGVTIGCAIFALSVLLNYKKIPR<br>WLVIIFALVILITSAYPSMGANVGGCISESVAYVLFILLIFDVKMDLKKSILLGLVPVILVGLFATLDIVSKSQSHLSGFVNQIMLNGPGAIIQTFTRKIQMN<br>VDIATSNIFIVIALIVAGFMSKFIFKPSRHFKKLADKYPYVFKGFVAIIVGCIVALIFNDSGIVAAGTGSIYILIPLLVMSVNMMIFDKE |
| A0A544QVA7 | Peptacetobacter hominis | DEKGKVIFINMSSTGLENMLSINAIEDRIQKEGYIGLMNIKGDQGNDAERGLASMGATGRATVGEVDekiSASFVNPDDREKKIFESATGKKAEAIINTEI<br>NVSVNENIEKGEFESTLGMLGQVMNENNLTTAVLGNADLYSEDGeiSLKNRAIGFLAMDNYGRVDAGNVDDINKEDLSMPYGVSTDYAKLKKETDK<br>LYKENDALFVDLGDTYRLDEYKPNLNEDSYAKMRAKVYSRINGYLEHVFNMAGENDTIYIVSAFPSNLDYINKKRLSPVIKFGGEGtsKGVLTSATTRR<br>DGIVANIDIGVDILNEFGLSNEMMLGRSLSEVNMDNNVEYLTSEYSKITSAALIRAGVVNGFVGVVSASWVIGLIAVLMSKRIPEKYRDglFKLLKELIKL<br>GIIMPLSFLLAPLLGLNTPVGMSIGVIAVALVIYAISRFAFKDDDIAQMGAVAAVTIAVIAIDSVFGTYLMKNNIMSYDAMVGARYYGIGNEYEGVTIGSAV<br>FAFAALLNYKRIPKWAVAVLSLVILITSAHPAMGANVGGAISESIAYLLFVMLIFNIRIDWKKVIVLGVAAAAVVCVFAYFDLSSGSPSHLGLFVQQIMLN<br>GPSEIVLTFTRKIQMNVQLAQTSVWVNILLVGIAIIAALIFKPTKHFKIIAHKYPFIFKGFVASMVGCIVTLFVNDSGIVAASTASIYILIPIMVITINMLFEKE |
| UPI001916EADB | Peptacetobacter hiranonis | RGKVIFINMSRTSLENMMNIKEIGNRVEKEGYVGLMNIKGDQGNDDARSLATMGATGRStIAQFDKKVAVDFVQPNVAEakEYEASTGKKAKKIVNM<br>EANRSSNENIEKGEFESTLGMVGQILSENGYKTSVIGNADYVDfDGQLVKNRNIAFAAMDNYGRVDDGNIDDINIKDNSMPFGISTDYKKLKAETKEL<br>YKNTDAIYVDLGDTYRLDEYKMNLNENTYAKMKAKTYSRINSYLKEVFSMVGENDTVYIVSAFPSNLDYSNKKRLSPVMKFSKGvsGKGVLTSATTRRD<br>GIIANMDIGVDILNHFGLSNEMMLGRTISNVEKENNIKFLQKEYSKINSAALIRAGVVNTFVGVVSASWVIGLIAVLLAKRiaPKYKDKLFTILKELIKFGII<br>MPLSFMLAPIFNFSGQVSMSLAVIGVTFLIYFIAKAMFKDDDLKQMGFIALVTIVLIAVDAVFGTYLMKNNIMSYDAMVGARYYGIGNEYEGVTIGSAVF<br>AFAVLLYYKKIPKWVAAASFGVILITSAYPSMGANVGGAISECAAYLLFVLLMYNVKVDWKKIVGIGLAVVAVVGVFAFMDLSSGSPSHLGLFVQQIML<br>NGPGEIVLTFTRKIQMNVKLAQTSVWVNILIVGIVIIAALIFKPNKHFRVIAGKYPMVFRGFVAAMVGCIITLLVNDSGIVAASTASIYILIPVMIITINMILNE<br>DK |

| | | |
|---|---|---|
| UPI000A617BE2 | Paraclostridium benzoelyticum | DSKHKTIVINLNRSNFSDFLEISTIKKELENRGYIGLMNIRGDQGTDDNRAYASIGAGGRVNTYNTDFSGFKNMDKDNSVIYESATGKKAKGINDVNIN<br>LSINDNLEKGKYGSTLGELGQALSDNKLRVSVLGNADIGTKPEELNRNIAAMAMDNYGRVASGNVDNIDIKDDKMPFGISTDYEKLKRETKKYYNES<br>DTLFVELGDTYRLDLYKKYLNEKTYQDMEEQIHKNIDSYLKEVFKLIGENDKVYIISAFPKDIDYKNQRRLSPVIKFDGNSKGLLTSATTRRDGIIGNSDI<br>GASILSDYNLTSEKMVGRKLENIKNDKNVDFIRSEYSKIVNMASVRSGIIQFFVNFVSIILILTIMLLTFRKKIKNKlsmkiDKVFYVFKELNKFGIIMPAAFLL<br>GPILNLNSKIGIYTSIGLMGIIIYGIGKLVFKTDFKQIGFYATLTILVILIDAVLGTYLMQNSVMSYNPIVGARYYGIGNEYEGIVIGSSIVAMTVLIMDKKLKKI<br>FVPIISVVIVFISSTPTMGANVGSAISATLALILFTLLVFNIKLDIRKIIFVFIAVGLVVFGFAAIDMISGSESHLALFINQILVDGPAAIIETFGRKISMNMKLM<br>QTSSWVYIIYIGIAGLIISKISCKKEFKDIMSKDSYIKKALISIGAGAVITFLVNDSGVVSAATTMLYIIIPLLILLVNKVILKEK |
| UPI00115BE473 | Paraclostridium bifermentans | SDHKTIVINMNRSNFVDFLEIPTLKKQLENRGYIALMNVRGDQGTGDNRSYASIGAGGRVNTYSTDFRGFKDVDKDNKITYESATGKKAKGINDTDIN<br>LSINDNLDKGKYGSTLGELGQTLNQNKLSVSVLGNADIGIKPEELNRNIAAMAMDNYGRIDSGNVDDINRNDESMPFGLSTDYDKLKSETKKQYKKS<br>DVLFVELGDTYRLDMYKSYLNDNTYKSMEKKIHKNIDEYLKEVFSMVNENDTVYIMSAFPKDIDYKNKRRLSPVVKFDGSGKGLLTSATTRRDGVIGN<br>VDIGAGILADYKITSDNMVGRELENVKNEDNINFMGKEYSKIVNISDVRSGIIQFFINSTSIVLGLSIVLLIFRkklenKYNKRIKKAFNLLKELSKFGLIMPV<br>AFLLAPMFNFNTKIGIYLSIFALTLAIYFISRLIFKSDNKQIGFYATLSIFVIIVDALLGSYLMKNSIMSYDAIVGARYYGIGNEYQGIIIGSSIAAMSILIINKKLP<br>KIIVPILSLLILFISASPTMGANVGSAISESVALILFTMLVFNVKIDFKKAILILFGAGLVVLGFAAIDMLSGSESHLSLFINQILVNGPTAILETFGRKISMNIQ<br>LMQNSSWVYIIYVGVIGLIVTSIFYKKEFKEMMKNESYGKKAIVSIGAGAIVTFLVNDSGVVSAATTMLYIIIPFLIILINKVIFKE |
| UPI0019824039 | Paraclostridium dentum | DSKHKTIVIDLNRSNLSDFLEIPTLKKELENRGYVALMNIRGDQGTGDNRAYASIGAGGRVNTYDTDFSGFKNVDKDNSVIYESATGKKSKGVNDVDI<br>NLSINDNLEKGKYGSTLGALGQALSDNKLSVSVLGNADIGTKPEELNRNIAAMAMDNYGRVDSGNVDNINKKDDKMPFGISTDYEKLKKETKKYYNE<br>SDTLFVELGDTYRLDLYKKYLNEKTYQDMEGQIHKNIDSYLKEVFNLIGENDKVYIISAFPKDIDYKNQRRLSPVVKFDGSDKGLLTSATTRRYGIIGNS<br>DIGASILSDYNLTSEKMVGRKLDNIKNDKNIDFITNEYSKIVNMASVRSGIIQFFVNFVSIILLLSIILLTFRKKVKNKlsmktDRVFYLVKELNKFGIIMPAAF<br>LLGPILNLNSRIGIYTSIALMGMIIYGIGKLAFKTDFKQIGFYATLTIVAILVDAVFGTYLMQNSVMSYNPIVGARYYGIGNEYEGIVIGSSIVAMTVLIMDKK<br>LKKIFVPIISVVIVFISSTPAMGANVGSAISATLALILFTMLVFDIKLDIRKIIFVFIAVGLVVFGFAAIDMLSGSESHLALFINQILVGGPAAIIETFGRKISMN<br>MQLMQTSSWVYIIYIGIAGLIISKISCKKEFKDIMSEGSYIKKALISIGAGAVITFLVNDSGVVSAATTMLYIIIPFLILLINKVVLKEK |
| UPI00202CFBEE | Paraclostridium ghonii | DDNHKTIVINMNRSNFLDFLEIPTLKKELENRGYIALMNVRGDQGTGDNRSYASIGAGGRVNTYNTDFIGFKNVDKDNSIVYQSATAKKAKGVNDTNI<br>NLSINDNLDKGKYGSTLGELGQTLNNNDLNISVLGNADIGTKPEELNRNIAAMAMDNYGRVDSGNVDDINKKDEKMPFGISTDYNKLKSETKKYYDK<br>SDVIFIELGDTYRLDLYKPYLNENTYKSMEKEIHKNIDSYLQYIFNMIDENDTVYIMSAFPKDIDYKNKRRLSPVVKFDGNGKGLLRSTTTKRDGIIGNVD<br>IGAGILADYDLQSKHMVGKELKNINRDDSREFIEKEYKKIVSISSVRSGVIQFFINSVSIILGITIVLLLFKNKidcksKLNTKKIFNILKELSKLGLIMPIAFLLA<br>PVFNVNTKIGIYLSISIFTLSIYLISKLIFKSDNKQIGFYASLSIIVIIIDALLSSYLMKNSIMSYDPIVGARYYGIGNEYQGIIIGSSISAMSILIMNKKIPKIIVPIV<br>SLIILFVSAAPTMGANVGSAISESVALILFTMLVFDVKIDLKKAILILFGAGLVVLGFAAIDMLSGSESHLSLFINQILTSGPTAIIETFGRKISMNIQLMQTS<br>SWVYIIYVGVIGLIITRIFYKKEFNEIMSDGSYVKKALISIGSGAIVTFLVNDSGVVSAATTMLYIIIPFLTIIINKVIFKEK |
| T4VML6 | Paraclostridium bifermentans ATCC 638 = DSM 14991 | HKTIVINMNRSNFVDFLEIPTLKKQLENRGYIALMNVRGDQGTGDNRSYASIGAGGRVNTYSTDFRGFKDADKDNKTTYESATGKKAKGVNDTDINL<br>SINDNLDKGKYGSTLGELGQTLNNKDLSVSVLGNADIGTKPEELNRNIAAMAMDNYGRIDSGNVDSINKIDDSMPFGLSTDYNKLKNETKNQYDKS<br>DIVFVELGDTYRLDLYKPYLNENTYKSMEKKIHKNIDEYLKEVFSLVDENDTVYVMSAFPKDIDYKNKRRLSPVVKFDGTGKGLLTSATTRRDGVIGNV<br>DIGAGILADYKLTSNNMVGRELENIKNEDNINFIEKEYSKIVNISDVRSGIIQFFINSTSIVLGLSIILLIFRKKLENKynksiKKAFNILKELSKFGLIMPVAFLL<br>APMFNFNTKIGIYLSIFALTLAIYFISRLIFKSDNKQIGFYVTLSIFVIIVDALLGSYLMKNSIMSYDAIVGARYYGIGNEYQGIIIGSSIAAMSILIINKKLPKIIV<br>PILSLIILFISASPTMGANVGSAISESVALILFTMLVFNVKIDFKKAILILFGAGLVVLGFAAIDMLSGSESHLSLFINQILVNGPTAILETFGRKISMNIQLMQ<br>NSSWVYIIYVGIIGLIATSIFYKKEFKAMMKNESYGKKAIVSIGAGAIVTFLVNDSGVVSAATTMLYIIIPFLIILINKVIFKE |
| A0A0M3DMP4 | Paraclostridium benzoelyticum | HKTIVINMNRSNFVDFLEIPTLKKQLENRGYIALMNVRGDQGTGDNRSYASIGAGGRVNTYSTDFRGFKDVDKDNKITYESATGKKAKGVNDTDINLS<br>INDNLDKGKYGSTLGELGQTLNKNNLSVSVLGNADIGTKPEELNRNIAAMAMDNYGRIDSGNVDSINKIDESMPFGLSTDYDKLKTETKKQYDKSDV<br>VFIELGDTYRLDLYKSYLNENTYKSMEKKIHKDIDEYLKEVFSIVDKNDTVYIMSAFPKDIDYKNKRRLSPVVKFDGSGKGLLTSATTRRDGVIGNVDIGA<br>GILADYKLTSDNMVGRVLESIKNEDNINFIEKEYSKIVNISDVRSGIIQFFINSISIVLGLSIVLLLFRKKLESKykksiKKVFSVLRELSKFGLIMPIAFLLAPM<br>FNFNTKLGIYLSICTLTLVIYFISKLIFKSDNKQIGFYATLSIFVIIVDALLGSYLMKNSIMSYDAIVGARYYGIGNEYQGIIIGSSIASMSILVINKKLPKLIVPIL<br>SLIILFISASPTMGANVGSAISESVALILFTMLVFNVKIDFKKAILILFGAGLVVLGFATIDMLSGSESHLSLFINQILANGPTAILETFGRKISMNIQLMQTS<br>SWVYIIYVGIIGLIVTSIFYKKDFTEMMKSESYGKKAIVSIGAGAIVTFIVNDSGVVSAATTMLYIIIPLLILLVNKVILKEK |

| | | |
|---|---|---|
| UPI0022E63257 | Paeniclostridium hominis | HKTIVINMNRSNLSDFLEIPALKNELKNRGYIALMNVRGDQGTDDKRSYATIGAGGRVNTYSDDFKGFKDVDKENRVLYESATGKKAKGVNDTDINL<br>SINDNLENGKYGSILGELGQTLSDKNKKVSVLGNADTGLEPKDLNRNIAAMAMDNYGRVDIGNVDKINKKNDEMPFGISTDYKKLENETKKYYNESD<br>VLFVELGDTYRLDLYKKYLNENTYAKMESKIHKNISSYLKELFNSISKNDKVYIISAFPKDIDYKNKRRLSPVVKFDGSGKGLLTSATTRRDGIIGNIDIAAS<br>ILADYNLSSKNIVGRPLETIKMKDNKSFIENEYKKIVSISDVRGNIIHSFINSMSIILILIIGLLVLKKPIKRKfkekyNKMFNILKELIKFGLILPLAFLLAPILNFN<br>TQLGIYISITAISIVIYVLGKLIFKEDNKQVGFYATISIIAITLDALLGSYLMKNSILSYDAIVGARYYGIGNEYQGIIIGSSIISMTILLVNNKLSKIMVPIISCIILF<br>LSASPSMGANVGAAISESVALILFTMLILDIKIDLKKSIFILVGAGAVVLGFAVLDILSGSESHLSLFVNQILTNGPSAIIETFTRKISMNIQLMQNSDWVNII<br>YIGILSIIIIKIVYSKVFKGLMDECVYAKKALISMGAGAIVTLLVNDSGVVSAATTMLYIVIPFLIIMVNKIVFKDK |
| A0A1M6PCK6 | Tepidibacter formicigenes DSM 15518 | NKGKVILININRTSINDLKEINSLSKELNKRGYIGLMNIRGSKGTSDIRSYASIGAGTRVYLNNDDINFRTINEENKDIYLRRTGINPKYINNLNINKLKSD<br>NEKGEYGAFLGALGYEIRNNNLSLAVLGNADIDEEKHREICLIGMDENGQIKKGNISDLNIKDSLMPFGVRTDYEKLLDETKKYYENSDLVIVELGDTY<br>RLDLYKENLSQNAYKNIKDKINKNISRYLNEVFKMAQEQDRIYIISPYPQIQDYKNGYRLSPIVIFEGNEKGLLSSSTTRRKGIIGNVDVSADILDYFDVK<br>SKVMVGKKIEKTVEDNNIGFLLGDYEKTVSNSKIRVPVLYTYAVLEMIAWILTVLAVFFRNKIPRNSFKILSSFLKFTMTIPFVLLIAPLFNFKSIISIVICIITTL<br>SLIYYLIHKFVKDDLSKLVILSLLVALGVLIDGATGQNMIKNSLLGYDPIIGARYYGIGNEYMGVLIGSLIFSLVGLLENKKIGKKIVSIILLISIVILGHPKMG<br>ANVGGTITAVFSFLYLILRLYDVKIDIKKIILICIAVVCVVSTMAIIDiYFIGSKSHLAGAVQKIISGGPLTIIQIIKRKLEMNLRLIGITIWSKVLILGVVIVGILFYK<br>PFGILKKVCEKYSYLTKGWTAIVLGSIVGFLVNDSGVVAAATSIAYVIIPLLVL |
| UPI0005E0CCCE | Paraclostridium sordellii | DNNHKTIVINMNRSNFVDFLEIPALKSELEKRGYIGLMNVRGDQGTDDKRSYATIGAGGRVNTYSDDFKDFKDVDDASRILYESATGKKAEGVNNTAI<br>NLSINSNVESGKYGSTLGALGQTLSTNNKKVSVLGNADTSIEEKDFNRNIGAMAMDNFGRIDIGNVDKINKTNNKMPFGLQTDYKKLKNETKKCLDE<br>SDVLFIELGDTYRLDLYKQYLNDNTYKEMEKAIHKNISSYLNDVFKTLNKDDKVYVMSAFPKDIDYKNKRRLSPVVKFDGDGKGLLTSATTRRDGIIGN<br>VDIAVDILQDYNLKSENMVGRFLEKIKKSDNIEFISKEYSKIVNISIVRSGIIQFFTNSVAILLAIMIVLLAFKKPLKNKfnvkyEKACNLVKEISKFGLILPLGF<br>LLAPILNLNSPIGIYVSISAISLFIYFIGRLIFKEDSKQIGFYALVSVVVIILDSLLGSYLMKNSIMSYDAIVGARYYGIGNEYQGITIGSSIIAMAILIKNKKLPKI<br>LVPIISCIILFVSASPAMGANVGSAISESVALILFTMLIFDIKIDLKKAILILIGAGLVVLGFAAIDIFTGSESHLSLFINQILVGGPAAIIETFARKISMNIKLMQ<br>NSAWVNVIYIGIFALIIMRVLYSKVFKKLMKESAYFKKALIAIGAGAFVTLLVNDSGVVSSATTMLYIAIPFLITIMNKILKNEE |
| A0A1M5Q0F0 | Tepidibacter thalassicus DSM 15285 | NKGKVILININRTSISDLKEIDSLNKEMDKRGYIGLMNIRGSNGTSDIKSYATIGSGTRAYLSKGDIDFRTINEENDYIYLRRTGIKGKHINNLNINKLKND<br>NLKGEYGAFLGALGYEFKNNNLSLAVLGNADTDEQNHREICLIAMDDNGQIEKGNIDDLNIKDDLMPFGIKTDYKKLLGETKKYYKNSDLLIVELGDT<br>YRLDLYKENLNKQTYNKLKQDINKNISGYLNEIFKMVNKEDRVYIISTYPKMQDYKNGYKLSYVILFEGDKKGLLSSATTRRKGIIGNVDIPVDILDYFGI<br>NSKLMVGKKIEKILENNNINLLIGDYEKTVSNSKIRVPVLYTYAALEMIVWILTVLALFFKNNIPKNSFKILSSFLKFTMTIPFALLIAPLFNFKSVLSIIACTVI<br>VLISVYYLVHKFIKDDLNKLIALGLLVSIGVLIDGVSGQNMIKNSLLGYDPIIGARYYGIGNEYMGVLIGSLIFSLVGLLEKNKIAKKTILIILLISVVILGHPK<br>MGANVGGTITAVFSFLYLILRLYDIKIDMKKIILICIAVVCVVASMAVIDIFFiGSKSHLAGAIQKIISGGPLVIIQIIKRKIEMNLRLIGVTIWSKVLISGILIVGV<br>LFYKPIGIFKNICEEYSYLTKGWSAIVVGSTVGFFVNDSGVVAAATSIAYVIIPLLVI |
| A0A1V4IBH7 | Alkalithermobacter paradoxus | ENNGKVIFISINRTSLDEMLSIDRIKEELDSRGYIGLMNIRGSKGTSDIRSYASIGWGSRAYLNYRNIQFETLNESNKDEYIRRSGKKPKYINNLNINNTK<br>SDAAIGEYGALPGALGMEIRNNNLKLAVLGNADTDSEQRREICLIAMDENGQIEKGNISSIDVEDGLMPFGIRTDYEKLINETKKYYDESDLLFIELGDT<br>YRLDLYRNKLNSNTYNSMKEKIYANINNYLNEVFKIVNPNDRIYIVSPYPRLEDFRTGYRLSPVIIFDKDEKGILTSSTTRRKGVVGNIDIAPDILNYFGQE<br>SNAMLGRNIDRISMDNNIDFLTYEYEKIVSTYNIRIPTLYTYAVFQMITWIISVIFIFLRKKISKKLFNIPINILKFTMIIPFALLVEPILNLNSEISIIAVALLMSLI<br>GYFIVHKLVKDDLNKIILLSTLTFIGILVDAATGQNMIKNSLLGYDPIIGARYYGIGNEYMGVLIGSIILALSSLFEKKKISRLLLSIVLLVCVIILGYPKMGAN<br>VGGTITAVFAFLFLIMKIYEIEVDIKKIVLIMCAVVLVVTAMALIDINFiGSKSHLAGAIEDIIIGGPIIILQIIRRKLEMNLRLIGISMWSRVLILGVIIVGVLFNK<br>PSGILKSLCEVYPSIAKGWQAIIVGSVVGFVVNDSGVVAAATSISYVIVPVLVLIMKDIVKSK |
| UPI00162A1A05 | Peptostreptococcus russellii | NGGKVIVIDINRTSLENFENIKFLRNKMEKEGYVGLMNIRGDKGYDDRRNYATFGATGRADINSDIPIDFKMANKKNIGIYEAATGQKAEAINLMNIND<br>LDLFNQTKGDYKSKLGYLGDTLVSDGKKISVLGNSDYYDnEGNYVKNRDFCLSVMDSKGRIGSGEIDKINYSNVNFPFGISSDYNKLKEKTDELYKDS<br>DLLFVDLGDTYRLDMYKTNLNEKTYKKMKFAIYNKVSDYIEHVFDIAGPNDTIYVMSSFPSKLDYSNNRRLAPVVRFdmSQKSRGLLQSSTTRRDGVI<br>ANMDIGVDILNRFGIKNKEMVGKVLTEKKMDGNLNYIFKEYKKIVAVSSIRMSIINIYVTFISVSWLVAALALWQKNRLPakYRDKILLMLKELVKFGLIM<br>PLAFLTAPILNPSSELQIALVIIAVSIIYYLIASKLFKGDDIMQIGFYSLVMILLITIDSAISTPLMQSNIMSYDPMIGARYYGVGNEYEGVTIGSAILGFAVLHE<br>KKNFPKWLTALLLAVILFISASPGMGANVGGAISECIAYLAFVLLIFGVKIDFKRMIGILVATVLLVAAFATADILLGMQSHLGNFVEQIRINGPMEVVYVF<br>ARKIAMNVQLAQTTVWVNILLVGLVILAITIFRPNRNFSLMKKKYPIIYDGFLATIVGCLVTLLVNDSGIIAAATDSIYLLIPILIILINKGVKNK |

| | | |
|---|---|---|
| UPI00202CC2CF | Paraclostridium ghonii | SKHKTVVINMNRSNFSDFLEIPILKKELKNRGYIGLMNVRGDQGTEDNRAYASIGAGGRVNTYSTDFTGFKSEDKDNSIIYQSVTGKKAKGINDTNINL<br>SINDNIEKGEYGSTLGELGQVLNNNDRKVSVLGNADIGIKPKELNRNIGAMGMDNYGRVDSGNVDNMNEKNESLPFGISTDYNKLKIETRKYYNES<br>DVLFIELGDTYRLDLYRKYLNENTYKSMEEEIHKNIDSYLKNVFDLINKNDKVYIMSVFPKDIDYKNQRRLSPVVKFDGNDKGLLTSATTRRTGIIGNSDI<br>GAGILADYNLVSDKMVGRKLDTIKNNNNLDFISKEYSNIINMASVRSGIIKLFINFISIILGLTVILLLIRKKTENKfntNTIFNILKELSKFGIIIPIAFLLGPILNL<br>NSQLGIYISIILIGICIYIIGKVMFKQDYKQIGFYATLTILVVLADATLGTYLMQNSVMSYNPIVGARYYGIGNEYEGVVIGASIVSMGVLIFEKKLNKVFVPLI<br>SAAIVFISSTPTMGANVGSAISTTLALILFTLFVFDKKVDFKKVVFIFIGIGVVVLGFAVIDMLSGSKSHLSLFINQILVSGPTAIIETFGRKISMNIQLIQTSS<br>WVYIIYVGIIGLIIARIFYEKEFCEIMRNESYVKKALISIGVGAVITFIVNDSGVVAAATTMLYIIIPFLIIIVN |
| A0A150FQR1 | Alkalithermobacter thermoalcaliphilus JW-YL-7 = DSM 73 | DTNGKVILVVINRTSLENMLSIDRIKQELNDRGYIGLMNIRGSRGTSDVRSYATIGSGTRAYINYRSLDFQTLDEQSKKIYERRTGKKAKYINNLNINTLR<br>SESLAGEYSAVLGALGTELKNNDLRVSVIGNADTDTEQRREICLIGMDENGQIENGNIDFINIDDNTMPFGIKTDYKRLIADTKKFYDKSDLLVVELGDT<br>YRLDLYRNNLNLNSYNKMKEKIYDNISGYLDELFNIASENDRIYILSPYPTLEHFRAGYRLSPVIVFEKDSKGILASSTTRRKGLIGNVDIPVDILEYFGKE<br>SPAMIGRTIQKISKDNNIDFLNYEYEKIVSTYNIRIPTLYTYAVFQMIVWIIALLAIFMKKKIPKQLFNAFVAILKFTLVIPFVLLIEPILNLNSELSIILAAIGLSF<br>VAFYIVHKFVKDDLNKITILSSITFLGILIDAATGQNMIKNSLLGYDPIIGARYYGIGNEYMGVLIGSIILSLSSLLEKNKISKLVLSLVLFMCIFILGHPKMGA<br>NVGGTITAVFAFTFLILKVYDVKVDIKKIILILTLVVLVVGTMAFIDInFSKSQSHLGGAIQDIITTGPIIIIQIIRRKIEMNLRLIGVSIWSKVLILGVIIVGILFNKP<br>SGILKKLCDIYPNIEKGWQAIIAGSIVGFLVNDSGVVAAATSILYVIVPVLLL |
| UPI0020BFAB6B | Tepidibacter aestuarii | EKGKVILVNINRTSINDLKNIEYLKNQMDKKGYVGLMNIRGSKGTSDIRSYAAIGAGTRTYINPSDIDFETSNENNKEVYLRRTGINPGYINNLDINKTNA<br>SNEKGEYGAFLGAIGYEFNNNNLDLAILGNSDTDEAKHREIALIGMDQNGQVQNGNIDNINIKDQTMPFGIRTDYDKLKKETQKYYDESDLLIVELGD<br>TYRLDLYKENLNEASYEKMKTEIYFNISDYLKEVFEMANENDRIYITSAYPEKQDYIDGYRLSPIVIFEKDEKGLLTSGTTRRKGVLSNLDIAADIFEYFGIK<br>SDVIVGKTIQKIPYDNNMEFLSKDYDKIVSNMKIRIPVLYTYAAFEMIMWMIMLAAIFFRHKTPKNTFKVLCAILKFTISIPFVLLIAPLFNYSTEASIIICIVGL<br>LIIVYSLVYKFVKDDLNKLILLSALVAVGVLIDAATGQNMIKSSLLGYDPIIGARYYGVGNEYMGVLIGSLIFFVVGLLEKNIINKKIASVILIVSIAILGHPKM<br>GANVGGTITSVFSFLYLIMRLYDIKIDLKKIILIGVAVVGVVSTMAIIDiYFIGSQSHLAGAIKKITSGGPLVILQIITRKIEMNMTLIGVSIWSKVLILGLLIVVVL<br>FQKPVGILKKVCDKYQYLTKGWTSIIVASIVGFLVNDSGVVAAATSIAYVIIPVLVL |
| A0A2P7Q1A7 | Peptostreptococcus russellii | NGGKVIIIDINRTSFENFENIKILREKLDKEGYIGLMNIRGDKGYDDRRNYATIGSTGRADIRSDISMKFNHSNKESANIYKSATGLEPGKINLENINSMD<br>LYNQTKGEYKSKVGYLGDSLISGGKKIAVLGNSDhYDEDGKALNNRDFCLSVMDSKGRIYAGNIDNINKSNYDFPYGISTDYKKLEEETNKYYKETDL<br>LFVNLGDTYRLDEYKENLNEKTYKQMKFSIYNKVSDYIEHVFNMVGPNDTVYIMSSFPSKLDYVNNRRLAPVVRFdmSDKSKGILQSSTTRRDGVLA<br>NMDIGVDVLNRFGLKNKEMVGKKLIGKKMDNNIEFLLKDYKKIVAVSSIRMDIINIYVTFISISWIVAAIALWQKNKLPakYRNNVLLVLKELVKFGLIMPL<br>AFLTAPIFNPTSGFQISILIVAVSLIYYLLARKLFKNDDIMQIGFFSLLMILIITIDSSISTPLMQSNIMSYDPMIGARYYGVGNEYEGVTIGSAILGFAVLVER<br>KKIPKWLTTILLAIILFISAYPGMGANVGGAISECIAYLAFVLLIYDVKIDFKKSMMILIITALLVVVFAGADIMLGMESHLGNFVEQIRINGPMEVVYVFGR<br>KIAMNVQLAQTTIWVNILLVGIVILAITIFRPNKNFSLIKKNHGTIYNGFLATIVGCLVTLLVNDSGIIAAATDSIYLLIPILIILINKGVKDK |
| UPI00189A156B | Peptostreptococcus anaerobius | NKGKVIVIDMNRVNMDNMSRFKSISGLIDKSAFIGQMNIRGDKGYDDRRNYASIGSTGRVNILPETRISFEKSTEDQKKIYQAATGTKALNINLDNINFI<br>DFYNQNKGEYKSSIGYLGQTLSSNGKKIGVLGNSDYYDSTgQFVKNRDFCLAVMDSRGRIHDGVLDGINKKKTNFPYGIGTDYKKLREETKRLYESS<br>DLLFVELGDTFRLDEYKENLNKNTYAKMKRMVYSEANKYIASTLKMAGDNDTIYIIGSFPSRLDYKNNRRLAPVIRIDKseKGSGLLETSTTRRKGVIANI<br>DLGVDILNRFGLKNKFMNGRVIKSVPMENSFEYLQKDYKKIIAISSIRMSIINIYVTIIVASWIICALALWKKEKIPQKHRgkVLNVLKEFVKLGLIMPLAFLT<br>APILQARTEPGVALSIIAATIVYYLAGRVLFKNDDMKQICLFASAMIAIIVVDSALYTPLMESNIMSYDPMIGARYYGIGNEYEGVTIGSAIIAFAILLEHKSIK<br>RWMVVPFLLLVLFTSAYPAMGANVGGAISECVAYIVFILLIYDIKIDFKKALMILLATGGLVVAFAIADIALGLGSHLGNFVNQILANGPMEIVNVFARKIE<br>MNVKLAQTTVWVNILLAGLAILALIIFRPNKNFAAMKAKYPLIYKGYLSIMVGCIVTLFVNDSGIISSATDSIYLLIPIIIIMINE |
| A0A929YXC9 | Peptostreptococcus sp. | NKGKVILIDLNRTNYDNFKNIKFLANKLENSGYLGMMNIRGDKGYDDRRNYASMGATGRVNILPDTPVNFSQLDNKDLYNYATGQSPKKINLLSINN<br>LDQYNQADGEFKSKMGYLGDTLTTNGKKISVLGNSDYYDmlTGQEIKNRDFALMVMDSRGRVEDGNVDFINKKDQSFPFGISTDYNKLKEETKKYY<br>AKSDLLMVNLGDTYRLDEYKVNLNSKTYTRMKYRVYNKISDYIEYVFKMAGKNDTIYILGSFPSKLDYANNRRLAPLVRFdmSDTGKGLLFSSTTRRTG<br>LFANLDMGVDILSRFGLKNSEMVGRQLTNKAMANRDDYMAKEYKKIVAISSIRMSIINIYVAVISISWILGALALWQRDKLPkkHKKNILNFLKEMVKLG<br>LIMPLAFLSAPILRPGSQVQITLAIVFMTFLLYILGNRLFKNDDLKQVGFFSILMILLIVIDSVISTPLMQSNIMSYDPMIGARYYGIGNEYEGATIGSAILGI<br>AILLEYRKIPKPWVAIFLIVVLFTSAYPSMGANVGGAISESIAYIAFFLLICGVKIDFKKSVLVVAVTGLIVVAFAAADIILGLGSHLGNFVSQVIANGPMEIV<br>TVFARKIEMNVQLAQTTVWVNILLVGLLILGFTIFRPNRHFSHMKERYKIVYKGYLAIVIGCLVTLLVNDSGIIAAATDSIYLLIPIIIILIN |

| | | |
|---|---|---|
| UPI001A49D4EB | Peptostreptococcus stomatis | NKGKVILIDLNRTNYDNFKNIKFLANRLENSGYLGMMNIRGDKGYDDRRNYASMGATGRVNILPETPVDFSQIEGKNLYKYATGHSPKKINLLSVNNL<br>DQYNQSNGEFKSKMGYLGDTLSSNGKKIGILGNSDYYDmvTGQEIRNRDFALMVMDSRGRIEEGNVDFINKKDQSFPFGISTDYDKLKDETKKYYAK<br>TDFLMVNLGDTFRLDEYKVNLNGKTYAKMKYRVYNKVSDYLEYVFKMAGKNDTIYILGSFPSKLDYANNRRLAPLVRFdmSESGKGLLLSATTRRAGV<br>FANLDLGVDILNRFGLKNKEMVGRQLTNKTMDNRDDYMTKEYKKIVAISSIRMTIINIYVAVISVSWILGALALWQRDKLPkkHKKKILNFLKEMVKLGLI<br>MPLAFLSAPILRPGSQAQVTLSIVFMTVLLYILGSRIFKNDDLKQVGFYSILMILLIVVDSVISTPLMQSNIMSYDPMIGARYYGIGNEYEGATIGSAILGV<br>AILLEYKKIPKSWVAIFLFAVLITSAYPSMGANVGGAISESIAYIAFFLLICGVKIDFKKSVLVVAIAALLVAVFAAADIMLGMGSHLGNFVGQVFRNGPME<br>IVTVFARKIDMNIQLAQTTVWVNILLVGLLILGFTIFRPNRHFSYMKERYKIVYDGYLAIIVGCIVTLLVNDSGIIAAATDSIYLLIPVIIILINKFNQKED |
| A0A095XQR5 | Peptostreptococcus sp. MV1 | NKGKVILIDLNRTNYENLKNIKFLAGRLEKSGYLGMMNIRGDKGYDDRRNYASMGATGRVNILPDTPVNFSQFEGKNLYRYATSQSPKKINMLGVNN<br>VDQYNQTKGEFKSKLAYLGDTLVSNGKKVAVLGNSDYIDMNTGqeIRNRDFALMVMDSRGRIADGNVDLINKKDQTFPFGISTDYDRLKKDTQNYY<br>KKSDFLMVNLGDTYRLDEYKVNLNKKTYTRMKYRIYKPISNYLEYVFDMAGPNDTIYIMGSFPSNLDYANNRRLAPLIRFdmSESSKGLLMSSTTRRKG<br>IFANLDLGVDILNRFGLKNPEMVGRVLTNKPMDNRDDYMAKEYKKIIAISSIRMSIINIYVAVISISWILGALALWQRDKLPkkHKKKILNFLKEMVKLGLI<br>MPLAFLSAPILRPGSQVQIALAIVFMTVLLYILGSKIFKGDDLKQVGFYSILMILLIVIDSVISTPLMQSNIMSYDPMIGARYYGVGNEYEGATIGSAILGLAI<br>LLEYKKIPKAWVAIFLFIVLITSAYPSMGANVGGAISESIAYIAFVLMIFGVKIDFKKSILVVLVSGLIVVVFAAADIMLGLGSHLGNFIRQVLANGPMEIFTV<br>FARKIDMNIQLAQTTVWVNILLIGLLILGFTIFRPNRNFSYMKESHKIVYDGYLAIVVGCLVTLLVNDSGIIAAATDSIYLLIPIIIILINKFNQKED |
| UPI000C06F93D | Tepidibacter mesophilus | EKGKVILININRTSITDFENIEYLNNKMNETGYVGLMNIKGSNGTSDIRSYASIGAGTRAYINPLDIDFRTLNENNKEIYLRRTGINPGYINNLDVNKTNVS<br>NEKGEYGAFLGAIGYEFNNKNLDLAILGNADTDEEKNREIALIGMDQNGQIENGNIEDINIKDQTMPFGIRTDYDKLKEETQKYYDENNLLIVELGDTY<br>RLDMYKENLNEASYEKIKREIYSNISDYLEKVFEISNKDDRIYIISAYPEKQDYIDGYSLSPIVLFDGEGKGLITSGTTRRKGVLSNLDISADIFDYFGIKSDV<br>IVGKTIQKISYDNNMEFLSKDYDKIVSNMKIRIPVLYTYAAFEMIMWIIMLVAIFFRHKTPKNTFRVLCAILKFTISIPFVLLIAPLLNYSTKGSIITCIIGLLIIVY<br>FIVYKFVKDDLSKLILLSALVAGGVLIDGATGQNMIKSSLLGYDPIIGARYYGIGNEYMGVLIGSLIFSVVGMLQKNIINKKVASLILLASIVILGHPKMGA<br>NVGGTITAVFSFLYLIMTLYDIKVDLKKMIFIGVAVVGVVSTMAIIDiYFIKSQSHLAGAIKKITSGGPFVILQIITRKIAMNMKLIGVSIWSKVLILGLLIVIVLF<br>QKPVGILKKVCDKYSYLAKGWTSIIVASIIGFLVNDSGVVAAATSIAYVIIPVLVL |
| W8T468 | Peptoclostridium acidaminophilum DSM 3953 | KGKVIVIVANRMGMQQLLEIQVLKSELEKRGYAGLMNIRGSKGTSDVRSYASIGWGSRAYLDAEDVDIRTKDLLEQDVFKRRTAMEPMDINNIGINTIK<br>AKALEGEYSVFPGMLGEELKNNGLGISVVGNADTDMEKVRYSALMAMDKDGNVPSGEIEDINKPDSRMPFGIATDYEKLKKYTQEYYEKSEVTIVELG<br>DTYRLDAYRPYLNDKAYGIMQKNIYKSANDYIGYVLGSLaSENDRVYVLAPHTRLADYKNYQRLAPIMIFDGEESGIAYSDTTRREGVIGNIDVAADILD<br>YFGIRSELASGKPIEAKPKAAALSYLEKEYAKIASTYQVRIPALYTYAVFEMLLWISVFICIMKKDRVKSKYFNILSKSLQFTGVVPFVMLIEPAFGIVSKAW<br>TVAFILAVSAVIYFVMRRTLKTDLDRLIFLSVLNGAGILCDAVLGQEMIKRSILGYDSIIGARYYGIGNEYMGILIGAALLGVSGLLQKGKIKKSAAAAILVV<br>TMAILGFPKIGANVGGLITAATCFLLFVLKAYDVKIDFKRIILIGCFVVGVVVAMALLDiYVLGSKSHLAGAIQSIGEGGPQVIVQIIQRKIEMNYRLIGASI<br>WSKVFIIALLIVGILFYRPFGILKTICSENPWLAKGWISIIVGSFVGFVVNDSGVVAAATSISFVIISALVLSMNKIVSEK |
| UPI001626B81D | Peptostreptococcus canis | QGGRVIVIDLSRTSLENFDNIKFLRERLDNSGYIGLMNIRGDKGYDDRRNYATIGATGRVDIYTDAVIDFSKSTKNSRALYKASTGHTPAEINLNNINNI<br>DLLNQTYGEFKSKMGYLGDTLKAHDKRIAVLGNMDHYDEfGEYKKNRDFCLSLMDSKGRIYRGNIDDINKKDLRFPYGISTDYKKLKEETNDYLKMA<br>DLLYVNLGDTYRLDMYKPNLNKRTYKKMKFAIYDKISDYIEYVFSQADSSDTIYVISSYPSALEYNNNKRLAPVVRFdmSDNTMGLLTSSTTRRKGIIAN<br>LDLGVDILSRFGLKNSEMVGRKIISVENEKRDEYLMKEYEKIVAISSIRIPIINIYVTLVSFFIILAAITLWQRNKISFKRRykTLNIMKELIKLGLIMPLAFLTAP<br>VLQAKTQPMVAISIVIMTILYYVVVSKIFKGDDIKQIGIYSLLMIFVITFDSLLATPLMESNIMSYDPMIGARYYGIGNEYEGVTIGSALLGFSILLQNKKLRK<br>PIVTTILIIILFISAYPAMGANVGGAISGCIAFFAFILMVYDIKIDLKKIILIFFVTLLLVVGFAIADIMLGIGSHLGNFVEQITINGPMEVVTVFTRKIEMNVQLA<br>QTTIWVNILLATIVVISFSVYKPNKNLAPIKKEYNIIYKGFLSMIAGCFVTLLVNDSGIISAATCSIYLIIPIMIMMIN |

| | | |
|---|---|---|
| UPI000C7B66C0 | Peptostreptococcus faecalis | NKGKVILIDLNRTSLSNFNNIKFLREKMKNDGHVALMNIRGDKGYDDRRNFASMGATGRVNIQSEIELNITESDKKLASLYRSATGKNPGKINLNNIN<br>EVDLYNKTKGEFKSNVGYLGETLNLNGKKIAVLGNSDYYnSEGKFIKNRDFSLAVMDGSGRLYNGNVDDINKENLSFPYGISTDYEKLISETNEFYKTS<br>DLIFVELGDTFRLDEYKLELNESTYKTMKYSIYNKVSNYIEKVFEMAGENDTIYVMSSFPSNLDYSNNRRLGTVVRFdmSKKGKGLLTSSTTRREGVMA<br>NIDIGVDVLNRFGLKNDEMVGRKLTEVDKASSLSFIEKEYKKIVAISSIRMTVINIYVAFVSLSVIFGALALWKREKIPakYRKKSLDVLKELMKFGLIMPLA<br>FLTAPILGASSEVQLALSIVVISIAYYLIGTRIFKGDDIKQIGFFSIAMILLIVIDSVISTPLMKANIMSYDPMMGARYYGIGNEYEGVTVGSAILGMAVLLEYK<br>KIPKWLATLFLTVILFTSAYPGMGANVGGAISECIAYFVFVLMIYNVKLDIKKIILVLFSTAVLVAGFAMADILLGIGSHLGNFVKQIQLNGPMEIIYVFARKI<br>DMNIQLAQTTVWVNILLVGILMISIVIIRPTREFLQLKKDHKIIYSGFIAAIIGCLVTLAVNDSGIIAAATDSIYILIPITIMMINKKN |
| A0A9E9LZ28 | Peptostreptococcus equinus | GKIIIIDINRVNLDNMRTINAIKKRMNKASIALMNIRGDKGYDDRRNYASMGSTGRVNIDQEKQLIISNSSQKEKNMYKHATGLRAESINLLNINDIDAY<br>NQTKGEYKSKIGILGDTLKSANKKISVLGNSDYInEEGQFVKNRDFALSLMDSKGRIENGVLDDLNIEDSRYPYGFRTDYDKLLEGTKKYYNSSDVLMV<br>NLGDTFRYDEYKVNLNEKTKLNMKRMVYSDVNKYLEQVFNMAGSNDTIYVIGSFPSKVDFKNNKRLGSVIRFdmSEKTAGLLTSSTTRRKGLITNTDI<br>GADILNRMGIKNSEMNGRVIKSVPKHNHFEFLKKDYEKIVSIATMRMDIINIYVVIIVISWVLGALALWQRHKLPikHKKTIIIILKEMIKLGLIMPLAFQTAP<br>IFNASSKLGVSMAILFVSLVYYVIATMAFKGKDMRQLGFYALIMIGIIVVDSILSTPLMSSNVMSYDPIMGARYYGIGNEYEGVTVGSAILGIAILLEYKKIP<br>KLLLIPIMIFILFTSAYPAMGANVGGAISETIAYLVFVLLVFNIKIDFKKTMMIGISTFALVAVFAVADIALGMGSHLGNFVKQIMNNGPMEIIYVFARKIDM<br>NIQLAQTTVWVNILLVGLAFLALTIFRPNKEFVAIKKEYPIIYKGFLSIMVGCIITLLVNDSGIIAAATDSIYLLIPIIII |
| A0A1I2C7H1 | Peptostreptococcus sp. D1 | NGGKVIVVDISRTNLENFNNIKFIRTKLKSSGYVALMNIRGDKGYDDKRNFATMGASTRANISSDLELNFNDTLEGAKLYKSATGWKAGKINLVNINNV<br>DFVNLTSGDYKSKIGYLGETINANGKKISVIGNNDYFENGQIVKNRDFCLTFMDSKGRLFSGNTDDINKKDTNFPFGISTDYEKLKKETQKYYDESDLL<br>YVNTGDTYRLDKYKMNLNNKTYKKMKFSIYNKISDYLDYVFSIVGKNDTVYVMSSYPSNLEYINNKKLTPIIRFdmSDGGFGLLNSATTRRAGIISNVDF<br>GAEILNRFGLKNEEMIGKKLTSLEKNGNIDYLLKDFEKIVSISSIRMSVINIYVSLISIVIIAAVISLWQKNRIPIRFRDRiiYTLKEASKLGLIMPLAFLSAPML<br>QSSSKPMISLSIVFFSIIYYLVPKIIFKKDDIKQLGLYSLIMIAVICLDAFFETPLMESNIMSYDPMVAARYYGIGNEYEGVVIGSAILGFSVILEYKKFPKWII<br>GILLMGVLFISAYPSMGANVGGAISGMIAFLVYILLINNAKVDIKKCVLILLATILLVVVFAVADIVLNIGSHLGNFVNRIMVNGPMEIANVFLRKIEMNILL<br>AQTTIWVNLLLVALLLLSIAIFRPSKFSGAIKNRYPYVYKGFLSIIVGCLVTLLVNDSGIISAATDSLYLIVPIVII |
| A0A845LJR4 | Peptoclostridium sp. | KGKVVVIVANRMGMQELLEIKALKRELENRGYAGLMNIRGSKGSSDIRSYASIGWGTRAYLNVEDVDIRTKDPLEQDVFERRTAAEAAHINNVNINAL<br>QTTNLDGEYNVFPGLLGEELKNNGLSISVVGNADTDVAKVRYAALIAMDKDGNVPSGEIEDVSKVDSRMPFGIATDYDKLKKYTQQYYENSDVTIVEL<br>GDTYRLDDYKIYLNDKAYGIMKDSIYKNANDYIEYVLgSLTSEDDRVYVVSPYTRLADYKNYQRLAPIFIFEGDKAGVAYSDTTRRDGIIGNIDVAADILD<br>YYGISSELASGKPIEAKDKSDALAYLEKEYAKIAATYQLRIPVLYTYAVFEMLLWIIVFTCIMKKDRVKNKYFNLLSRTLQFTGVVPFVMLIEPAFGIVSKLG<br>TVAFILAVSSGIYYMMHRMIKTDLDRLIFLSVLNGLGILSDAVLGQEMIKRSLLGYDSMIGARFYGIGNEYMGILIGAALLAVSGLLQKGKITRSVTAGVL<br>VTTMIILGFPKIGANVGGLITAATCFLLFVLKSYNIKIDFKRIILIGCFVVGAVVSMALLDiYVLGSKSHLAGAIQSIGEGGPQVILQIIQRKLEMNYRLIGASI<br>WSKVFIIALLIVGILFYRPFGILKTICGENPWLAKGWISIIVGSFVGFAVNDSGVVAAATSISFVIISALVLCMNKIVSEK |
| A0A6N7WZA2 | Peptostreptococcus porci | GKVIIVDIIRTNLENFNNIKFIRTKLKSSGYVALMNIRGDKGYDDKRNFATMGASTRANISSDLELDFNDTLDGAKLYKSATGWKPGAINLVNINNIDFV<br>NLTTGDYKSKIGYLGETMNANGKRISVIGNNDYYENGQLVKNRDFCLTFMDSKGRLFGGNTDDINKKDTNYPFGIATDYEKLKKETQKYYNESDLLYV<br>NTGDTYRLDKYKMNLNEKTYKKMKFSIYNNVSDYLDYIFSIAGKNDTVYVMSSYPSNLEYINNKKLTPIMRFdmSDGGSGLLNSATTRRDGIISNVDFG<br>AEILNRFGLKNEEIIGKKLTSLEKNGNIDYLLKDFEKIVSISSIRMSVINIYVSLVSIVIIAAVISLWQKNRIPRRFRGRVIntLKEASKLGLIMPLAFLSAPMLQ<br>SSSKSMISLSIVFFSIIYYLVPKIIFKKDDMKQIGMYSLIMISVICLDAFFATPLMESNIMSYDPMVAARYYGIGNEYEGVVIGSAILGFSVILEYKKFPKWIIG<br>IILMGVLFISAYPSMGANVGGAISGMIAFLVYILLINNVKIDIKKCVLILLATVLLVAGFAVADIALNIGSHLGNFVKQIMVNGPMEIANVFLRKIEMNILLA<br>QTTIWVNFLLVALLLLSIAVFKPGKFSGAIKNRYPYVYKGFLSIIVGCLVTLLVNDSGIISAATDSLYLIVPIVII |
| UPI001A9A6391 | Peptoclostridium litorale | ESGKVILIMLSRTGMEDMLEIKSIKEELESRGYVSLMNIKGSKGTSDIRSYASIGWGTRAYLKPEDIDFVTSDDENRVIFKRRTGIEPKHINNININMLKSS<br>NESGEYGAYLGILQKELSKNGKTISVIGNSDIPEKNHSYASLLAMDENGSIESGKIDGINVQDDYMPFGMRTDYEALKEYTRDYYSKSDVLAIELGDTS<br>RLDLYRENLNQKTYGEMQNKIYGNISDYISYVMsDLADAKDRIYIVSPYPTLLEYKNYKRLSPAFIFEGQGKGVLYSNTTRREGVIGNVDIAADVLDYFGI<br>KSDFMVGRSIQKIERSDNTEFLGSEFEKMSTTYQLRMPVLYSYAVFEMLLWISIFICILIKGMIPKKVFSVLSEVLQLTVVMPFVLLVEPIFGATSKSPVLAI<br>IISLTVILYIALKKLIKGNIERFMVISMMTGIGILADAATGQELIKRSIFGYDAMIGARYYGVGNEYMGILIGASLLSVAGLLQKEKIKKAAAIIMLAITIVILGFP<br>KMGANVGGTITASFCFLFFVLRMYNVKIDFKKLVAISAFVCVVVAGMAFIDiFLIGSKSHLAGAIQSIISGGPVVIIQIMSRKLAMNASLIGGSIWSKVFIM<br>ALFIVGVLFYRPFGILKRVCNEKEFLKKGWISIIVGSFVGFAVNDSGVVSAATSISFVIISVLLLIMEKIAHEN |
