## Supplementary material for "Identification of two novel lipoteichoic acid synthases essential for viability in *Clostridioides difficile*": Table S3

**Table S3. Phenotypic classification of LtaB alanine-substitution mutants**

| Color Class | Residues | Domain | Viability | LTA phenotype |
|---|---|---|---|---|
| White | D35, N37, D68, D73, Y77, E91, N171, Y233 | LtaS-like | WT | WT |
| Yellow | N64, L293, T311, T312 | LtaS-like | WT | Intermediate-length |
| Orange | R313, S490, Y491, D492, S604, R612, N632 | GT-C-like | Intermediate | Intermediate-length |
| Red | R498, Y499, Y500, N504, H605, R627, K628, M631, N691, D692, S693 | GT-C-like | Severe defect | Short LTA |
