## Supplementary material for "Identification of two novel lipoteichoic acid synthases essential for viability in *Clostridioides difficile*": Table S4

**Table S4. Sequence identity of LtaA and LtaB homologs from representative bacterial species**

| Protein | Organism | Length (aa) | LtaA identity (%) | LtaB identity (%) |
|---|---|---|---|---|
| C672_RS04615 | *Paraclostridium bifermentans* | 723 | 53.4 | 58.17 |
| C672_RS06385 | *Paraclostridium bifermentans* | 726 | 50.92 | 51.99 |
| C672_RS06390 | *Paraclostridium bifermentans* | 726 | 48.41 | 49.79 |
| HMPREF1639_RS06015 | *Peptostreptococcus sp. MV1* | 760 | 49.04 | 52.78 |
| BA6348_RS26125 | *Brevibacillus agri* | 746 | 24.83 | 26.18 |
| NRS818 protein | *Brevibacillus borstelensis* | 756 | 24.34 | 25.07 |
| NRS-661 protein | *Brevibacillus laterosporus* | 782 | 23.97 | 25.43 |
