## Supplementary material for "Identification of two novel lipoteichoic acid synthases essential for viability in *Clostridioides difficile*": Table S5

**Table S5. Plasmids used in this study**

| Plasmid | Relevant features | Parent vector | Restriction enzymes to digest parent plasmid | Primers used | PCR template | Assembly | Reference |
|---|---|---|---|---|---|---|---|
| pBZ101 | $P_{xyl}$ empty vector | | | | | | Muh 2022 |
| pCE1244 | $P_{xyl}$::*ltaA catP* | pAP114 | BamHI, SacI | CDEP7274, 7275 | R20291 | ITA | |
| pCE1243 | $P_{xyl}$::*ltaB catP* | pAP114 | BamHI, SacI | CDEP7272, 7273 | R20291 | ITA | |
| pIA131 | $P_{xyl}$::*MCS-rfp catP* | | | | | | Alberts 2025 |
| pIA134 | $P_{xyl}$::*cdr_985-rfp catP* | pIA131 | SacI, KpnI | CDEP6588 , 6614 | R20291 | ITA | |
| pAV101 | $P_{xyl}$::*ltaA-rfp catP* | pIA134 | PmeI, KpnI | CDEP7360, 7361 | R20291 | ITA | |
| pAV107 | $P_{xyl}$::*ltaB-rfp catP* | pIA134 | PmeI, KpnI | CDEP7362, 7363 | R20291 | ITA | |
| pIA123 | $P_{xyl}$::*Cas9-opt* $P_{gdh}$::*sgRNA-cdr_0985* homology to delete *cdr_0985 catP* | | | | | | Bollinger 2024 |
| pCE1232 | $P_{xyl}$::*Cas9-opt* $P_{gdh}$::*sgRNA-cdr_0985* homology to delete *ltaA catP* | pIA123 | PstI | CDEP7196-7199 | R20291 | ITA | |
| pCE1236 | $P_{xyl}$::*Cas9-opt* $P_{gdh}$::*sgRNA-ltaA* homology to delete *ltaA catP* | pCE1232 | MscI, NotI | CDEP4237, 7201 | pIA123 | ITA | |
| pRFP185 | $P_{tet}$::*gusA catP* | | | | | | Fagan Fairweather 2011 |
| pCE1230 | $P_{xyl}$::*Cas9-opt* $P_{gdh}$::*sgRNA-cdr_0985* homology to knock-in $P_{tet}$ *catP* | pIA123 | PstI | CDEP7203-7208 | R20291, pRPF185 | ITA | |
| pCE1233 | $P_{xyl}$::*Cas9-opt* $P_{gdh}$::*sgRNA-ltaB* homology to knock-in $P_{tet}$ *catP* | pCE1230 | MscI, NotI | CDEP4237, 7211 | pIA123 | ITA | |
| pEW22 | $P_{xyl}$::*Cas9-opt* $P_{gdh}$::*sgRNA-ltaB* homology to delete *ltaB catP* | pCE1233 | AgeI, KpnI | CDEP7782, 7783 | R20291 | ITA | |
| pIA33 | $P_{xyl}$::d*Cas9-opt* $P_{gdh}$::*sgRNA-rfp catP* | | | | | | Muh 2019 |
| pIA34 | $P_{xyl}$::d*Cas9-opt* $P_{gdh}$::*sgRNA-neg catP* | | | | | | Muh 2019 |
| pBZ180 | $P_{xyl}$::d*Cas9-opt* $P_{gdh}$::*sgRNA-ltaA-1 catP* | pIA33 | MscI, NotI | CDEP4237, 8453 | pIA33 | ITA | |
| pBZ181 | $P_{xyl}$::d*Cas9-opt* $P_{gdh}$::*sgRNA-ltaA-2 catP* | pIA33 | MscI, NotI | CDEP4237, 8454 | pIA33 | ITA | |
| pAP114 | $P_{xyl}$::*rfp catP* | | | | | | Muh 2019 |
| pCE1321 | $P_{xyl}$::*ltaB*$^{D35A}$*-rfp catP* | pIA134 | PmeI, KpnI | CDEP7773, 7774 | R20291 | ITA | |
| pCE1322 | $P_{xyl}$::*ltaB*$^{N37A}$*-rfp catP* | pIA134 | PmeI, KpnI | CDEP7775, 7776 | R20291 | ITA | |
| pHL33 | $P_{xyl}$::*ltaB*$^{N64A}$*-rfp catP* | pIA134 | PmeI, KpnI | CDEP7865, 7866 | R20291 | ITA | |
| pHL40 | $P_{xyl}$::*ltaB*$^{D68A}$*-rfp catP* | pIA134 | PmeI, KpnI | CDEP7934, 7935 | R20291 | ITA | |
| pCE1323 | $P_{xyl}$::*ltaB*$^{D73A}$*-rfp catP* | pIA134 | PmeI, KpnI | CDEP7777, 7778 | R20291 | ITA | |
| pHL29 | $P_{xyl}$::*ltaB*$^{Y77A}$*-rfp catP* | pIA134 | PmeI, KpnI | CDEP7829, 7830 | R20291 | ITA | |
| pHL31 | $P_{xyl}$::*ltaB*$^{E91A}$*-rfp catP* | pIA134 | PmeI, KpnI | CDEP7831, 7832 | R20291 | ITA | |
| pHL32 | $P_{xyl}$::*ltaB*$^{N171A}$*-rfp catP* | pIA134 | PmeI, KpnI | CDEP7859, 7860 | R20291 | ITA | |
| pHL34 | $P_{xyl}$::*ltaB*$^{Y233A}$*-rfp catP* | pIA134 | PmeI, KpnI | CDEP7861, 7862 | R20291 | ITA | |
| pHL36 | $P_{xyl}$::*ltaB*$^{L293A}$*-rfp catP* | pIA134 | PmeI, KpnI | CDEP7936, 7937 | R20291 | ITA | |
| pHL37 | $P_{xyl}$::*ltaB*$^{T311A}$*-rfp catP* | pIA134 | PmeI, KpnI | CDEP7938, 7939 | R20291 | ITA | |
| pHL38 | $P_{xyl}$::*ltaB*$^{T312A}$*-rfp catP* | pIA134 | PmeI, KpnI | CDEP7940, 7941 | R20291 | ITA | |
| pHL39 | $P_{xyl}$::*ltaB*$^{R313A}$*-rfp catP* | pIA134 | PmeI, KpnI | CDEP7942, 7943 | R20291 | ITA | |
| pHL30 | $P_{xyl}$::*ltaB*$^{S490A}$*-rfp catP* | pIA134 | PmeI, KpnI | CDEP7833, 7834 | R20291 | ITA | |
| pBZ169 | $P_{xyl}$::*ltaB*$^{Y491A}$*-rfp catP* | pIA134 | PmeI, KpnI | CDEP7835, 7836 | R20291 | ITA | |
| pBZ170 | $P_{xyl}$::*ltaB*$^{D492A}$*-rfp catP* | pIA134 | PmeI, KpnI | CDEP7845, 7846 | R20291 | ITA | |

| | | | | | |
|---|---|---|---|---|---|
| pCE1317 | $P_{xyl}$::*ltaB*$^{R498A}$-*rfp catP* | pIA134 | PmeI, KpnI | CDEP7765, 7766 | R20291 | ITA |
| pBZ171 | $P_{xyl}$::*ltaB*$^{Y499A}$-*rfp catP* | pIA134 | PmeI, KpnI | CDEP7847, 7848 | R20291 | ITA |
| pBZ172 | $P_{xyl}$::*ltaB*$^{Y500A}$-*rfp catP* | pIA134 | PmeI, KpnI | CDEP7849, 7850 | R20291 | ITA |
| pBZ173 | $P_{xyl}$::*ltaB*$^{N504A}$-*rfp catP* | pIA134 | PmeI, KpnI | CDEP7851, 7852 | R20291 | ITA |
| pBZ179 | $P_{xyl}$::*ltaB*$^{S604A}$-*rfp catP* | pIA134 | PmeI, KpnI | CDEP7867, 7868 | R20291 | ITA |
| pCE1318 | $P_{xyl}$::*ltaB*$^{H605A}$-*rfp catP* | pIA134 | PmeI, KpnI | CDEP7767, 7768 | R20291 | ITA |
| pCE1319 | $P_{xyl}$::*ltaB*$^{R612A}$-*rfp catP* | pIA134 | PmeI, KpnI | CDEP7869, 7870 | R20291 | ITA |
| pCE1320 | $P_{xyl}$::*ltaB*$^{R627A}$-*rfp catP* | pIA134 | PmeI, KpnI | CDEP7771, 7772 | R20291 | ITA |
| pBZ175 | $P_{xyl}$::*ltaB*$^{K628A}$-*rfp catP* | pIA134 | PmeI, KpnI | CDEP7871, 7872 | R20291 | ITA |
| pHL41 | $P_{xyl}$::*ltaB*$^{M631A}$-*rfp catP* | pIA134 | PmeI, KpnI | CDEP7956, 7957 | R20291 | ITA |
| pBZ176 | $P_{xyl}$::*ltaB*$^{N632A}$-*rfp catP* | pIA134 | PmeI, KpnI | CDEP7873. 7874 | R20291 | ITA |
| pBZ177 | $P_{xyl}$::*ltaB*$^{N691A}$-*rfp catP* | pIA134 | PmeI, KpnI | CDEP7875, 7876 | R20291 | ITA |
| pBZ178 | $P_{xyl}$::*ltaB*$^{D692A}$-*rfp catP* | pIA134 | PmeI, KpnI | CDEP7877, 7878 | R20291 | ITA |
| pHL42 | $P_{xyl}$::*ltaB*$^{S693A}$-*rfp catP* | pIA134 | PmeI, KpnI | CDEP7958, 7959 | R20291 | ITA |
| pMK03 | $P_{xyl}$::*cdr_0985 catP* | pAP114 | BamHI, SacI | CDEP6588, 6589 | R20291 | ITA |
| pCE1324 | $P_{xyl}$::*ltaB*$_{P.\ bifermentans\ RS04615}$ *catP* | pMK03 | PmeI, KpnI | CDEP7805, 7806 | *Paraclostridium birfermentans* ATCC 638 | ITA |
| pCE1325 | $P_{xyl}$::*ltaB*$_{P.\ bifermentans\ RS06385}$ *catP* | pMK03 | PmeI, KpnI | CDEP7807, 7808 | *Paraclostridium birfermentans* ATCC 638 | ITA |
| pCE1326 | $P_{xyl}$::*ltaB*$_{P.\ bifermentans\ RS06390}$ *catP* | pMK03 | PmeI, KpnI | CDEP7809, 7810 | *Paraclostridium birfermentans* ATCC 638 | ITA |
| pCE1327 | $P_{xyl}$::*ltaB*$_{B.\ agri\ RS26125}$ *catP* | pMK03 | PmeI, KpnI | CDEP7815, 7816 | *Brevibacillus agri* DSM 6348 | ITA |
| pCE1328 | $P_{xyl}$::*ltaB*$_{Peptostreptococcus\ RS06015}$ *catP* | pMK03 | PmeI, KpnI | CDEP7813, 7814 | *Peptostreptococcus* sp. Strain MV1 HM-1102 | ITA |
| pCE1350 | $P_{xyl}$::*ltaB*$_{B.\ borstelensis\ EDM54\_08085}$ *catP* | pMK03 | PmeI, KpnI | CDEP8104, 8105 | *Brevibacillus borstelensis* NRS-818 | ITA |
| pCE1351 | $P_{xyl}$::*ltaB*$_{B.\ laterosporus\ ABER04\_18820}$ *catP* | pMK03 | PmeI, KpnI | CDEP8106, 8107 | *Brevibacillus laterosporus* ATCC 6456 | ITA |
