## Supplementary material for "Identification of two novel lipoteichoic acid synthases essential for viability in *Clostridioides difficile*": Table S6

**Table S6. Oligonucleotides used in this study**

| **Oligo** | **Sequence** | **Relevant features** |
|---|---|---|
| CDEP7274 | cgatagttatgaagtgagctctaaggagggagaatttatgagaaaaaagattatatcttt | Clone *ltaA* onto pAP114 |
| CDEP7275 | tttattaaaacttataggatccatatttacatattattcatacagtacttattcttttaatgct | Clone *ltaA* onto pAP114 |
| CDEP7272 | cgatagttatgaagtgagctctaaggaggtcaaataaatgagaaaaaagattatgtcg | Clone *ltaB* onto pAP114 |
| CDEP7273 | agttttattaaaacttataggatccctatatgaacagtaaaaaatttttagttgt | Clone *ltaB* onto pAP114 |
| CDEP6588 | cgatagttatgaagtgagctaaggagtttaaacaatggaagaaaataaacaagttgatagac | Clone *cdr_0985* onto pIA131 and pAP114 |
| CDEP6614 | agataccatagatccggtacctctattaatttgtgactcgaaatttatat | Clone *cdr_0985* onto pIA131 |
| CDEP6588 | cgatagttatgaagtgagctaaggagtttaaacaatggaagaaaataaacaagttgatagac | Clone *cdr_0985* onto pAP114 |
| CDEP7360 | gaagtgagctaaggagtttaaacaatgagaaaaaagattatatctttcttagc | Clone *ltaA* onto pIA134 |
| CDEP7361 | gataccatagatccggtaccttcttttaatgctgtgagatttatac | Clone *ltaA* onto pIA134 |
| CDEP7362 | gaagtgagctaaggagtttaaacaatgagaaaaaagattatgtcgttg | Clone *ltaB* onto pIA134 |
| CDEP7363 | gataccatagatccggtaccgttgttttttcaaatattatcatattta | Clone *ltaB* onto pIA134 |
| CDEP7196 | caggaagggcgaattctgcagaattaaatatagaagttgaaataagtgaga | Clone homology for *ltaA* deletion |
| CDEP7197 | acatattattcatacagtacaaattctccctgctttctga | Clone homology for *ltaA* deletion |
| CDEP7198 | tcagaaagcagggagaatttgtactgtatgaataatatgtaaatatgaaca | Clone homology for *ltaA* deletion |
| CDEP7199 | ccggtcagatctgcactgcagcagtacttgatttgtcaaaatttga | Clone homology for *ltaA* deletion |
| CDEP4237 | cttataggatccgcggccgctag | Clone *sgRNA* from pIA134 |
| CDEP7201 | aattaaactgtaaatggccacatataaaaatataccaaaggttttagagctagaaatagc | *sgRNA-ltaA* for *ltaA* deletion |
| CDEP7203 | caggaagggcgaattctgcaggaaagctaagatgagtttacaatc | Clone homology to knock-in $P_{tet}$ |
| CDEP7204 | gcttcttatttttatgctagctactttctattcagatagtgcg | Clone homology to knock-in $P_{tet}$ |
| CDEP7205 | actatctgaatagaaagtagctagcataaaaataagaagcctgc | Clone *tetR*-$P_{tet}$ from pRPF185 |
| CDEP7206 | tcatttatttgacctccttaggagctcagatctgttaacg | Clone *tetR*-$P_{tet}$ from pRPF185 |
| CDEP7207 | gatctgagctcctaaggaggtcaaataaatgagaaaaaagattatgtcg | Clone homology to knock-in $P_{tet}$ |
| CDEP7208 | ccggtcagatctgcactgcaggattattttcaattatatcagcatttcc | Clone homology to knock-in $P_{tet}$ |
| CDEP7211 | aattaaactgtaaatggccagcactatctgaatagaaagtgttttagagctagaaatagc | *sgRNA-ltaB* |
| CDEP7782 | cgcactatctgaatagaaagtagctgttcatataggatttaaataaaaaaatgc | Clone homology for *ltaB* deletion |
| CDEP7783 | tttatattttcagagaccgggcctctcatcccaaaactct | Clone homology for *ltaB* deletion |
| CDEP4237 | cttataggatccgcggccgctag | Cloning sgRNA into pIA33 |
| CDEP8453 | aattaaactgtaaatggccaatagtatctacatttccattgttttagagctagaaatagc | *sgRNA-ltaA-1* |
| CDEP8454 | aattaaactgtaaatggccatttttataatctaaatcactgttttagagctagaaatagc | *sgRNA-ltaA-2* |
| CDEP7773 | gaaaaaggcaaggtcatatttattgcaatgaataggaccaatcttgg | Clone *ltaB*$^{D35A}$ onto pIA134 |
| CDEP7774 | ccaagattggtcctattcattgcaataaatatgaccttgcctttttc | Clone *ltaB*$^{D35A}$ onto pIA134 |
| CDEP7775 | ggcaaggtcatatttattgatatggcaaggaccaatcttggaagtatg | Clone *ltaB*$^{N37A}$ onto pIA134 |
| CDEP7776 | catacttccaagattggtccttgccatatcaataaatatgaccttgcc | Clone *ltaB*$^{N37A}$ onto pIA134 |
| CDEP7865 | gtagctctcatggcaataagaggggatc | Clone *ltaB*$^{N64A}$ onto pIA134 |

| | | |
|---|---|---|
| CDEP7866 | gatcccctcttattgccatgagagctac | Clone *ltaB*$^{N64A}$ onto pIA134 |
| CDEP7934 | gaatataagaggggcacaaggtactg | Clone *ltaB*$^{D68A}$ onto pIA134 |
| CDEP7935 | cagtaccttgtgcccctcttatattc | Clone *ltaB*$^{D68A}$ onto pIA134 |
| CDEP7777 | ggggatcaaggtactgatgcaagaaggtcttatgcaagtatg | Clone *ltaB*$^{D73A}$ onto pIA134 |
| CDEP7778 | catacttgcataagaccttcttgcatcagtaccttgatcccc | Clone *ltaB*$^{D73A}$ onto pIA134 |
| CDEP7829 | gatagaaggtctgcagcaagtatggg | Clone *ltaB*$^{Y77A}$ onto pIA134 |
| CDEP7830 | cccatacttgctgcagaccttctatc | Clone *ltaB*$^{Y77A}$ onto pIA134 |
| CDEP7831 | gtgtaactactgcagactatatc | Clone *ltaB*$^{E91A}$ onto pIA134 |
| CDEP7832 | gatatagtctgcagtagttacac | Clone *ltaB*$^{E91A}$ onto pIA134 |
| CDEP7859 | caattagttaaagcaagaaacttatg | Clone *ltaB*$^{N171A}$ onto pIA134 |
| CDEP7860 | cataagtttcttgctttaactaattg | Clone *ltaB*$^{N171A}$ onto pIA134 |
| CDEP7861 | gaacttggagatacagcaagattagatatg | Clone *ltaB*$^{Y233A}$ onto pIA134 |
| CDEP7862 | catatctaatcttgctgtatctccaagttc | Clone *ltaB*$^{Y233A}$ onto pIA134 |
| CDEP7936 | gaataagagaagagcatctccaataataaaatttag | Clone *ltaB*$^{L293A}$ onto pIA134 |
| CDEP7937 | ctaaattttattattggagatgctcttctcttattc | Clone *ltaB*$^{L293A}$ onto pIA134 |
| CDEP7938 | ctggaatctgccgcaacaagaaaggc | Clone *ltaB*$^{T311A}$ onto pIA134 |
| CDEP7939 | gcctttcttgttgcggcagattccag | Clone *ltaB*$^{T311A}$ onto pIA134 |
| CDEP7940 | ggaatctgccacagcaagaaaggctgg | Clone *ltaB*$^{T312A}$ onto pIA134 |
| CDEP7941 | ccagcctttcttgctgtggcagattcc | Clone *ltaB*$^{T312A}$ onto pIA134 |
| CDEP7942 | ctgccacaacagcaaaggctggaattg | Clone *ltaB*$^{R313A}$ onto pIA134 |
| CDEP7943 | caattccagcctttgctgttgtggcag | Clone *ltaB*$^{R313A}$ onto pIA134 |
| CDEP7833 | taacataatggcatatgatgctg | Clone *ltaB*$^{S490A}$ onto pIA134 |
| CDEP7834 | cagcatcatatgccattatgtta | Clone *ltaB*$^{S490A}$ onto pIA134 |
| CDEP7835 | taacataatgagtgcagatgctgtc | Clone *ltaB*$^{Y491A}$ onto pIA134 |
| CDEP7836 | gacagcatctgcactcattatgtta | Clone *ltaB*$^{Y491A}$ onto pIA134 |
| CDEP7845 | cataatgagttatgcagctgtcatagg | Clone *ltaB*$^{D492A}$ onto pIA134 |
| CDEP7846 | cctatgacagctgcataactcattatg | Clone *ltaB*$^{D492A}$ onto pIA134 |
| CDEP7765 | gatgctgtcataggagctgcatactatggtgtaggaaatgaatatg | Clone *ltaB*$^{R498A}$ onto pIA134 |
| CDEP7766 | catattcatttcctacaccatagtatgcagctcctatgacagcatc | Clone *ltaB*$^{R498A}$ onto pIA134 |
| CDEP7847 | cataggagctagggcatatggtgtagg | Clone *ltaB*$^{Y499A}$ onto pIA134 |
| CDEP7848 | cctacaccatatgccctagctcctatg | Clone *ltaB*$^{Y499A}$ onto pIA134 |
| CDEP7849 | ggagctaggtacgcaggtgtaggaaatg | Clone *ltaB*$^{Y500A}$ onto pIA134 |
| CDEP7850 | catttcctacacctgcgtacctagctcc | Clone *ltaB*$^{Y500A}$ onto pIA134 |
| CDEP7851 | ctatggtgtaggagcagaatatgaaggtg | Clone *ltaB*$^{N504A}$ onto pIA134 |
| CDEP7852 | caccttcatattctgctcctacaccatag | Clone *ltaB*$^{N504A}$ onto pIA134 |
| CDEP7867 | caggtagtgaagcacatttaggtatg | Clone *ltaB*$^{S604A}$ onto pIA134 |
| CDEP7868 | catacctaaatgtgcttcactacctg | Clone *ltaB*$^{S604A}$ onto pIA134 |

| | | |
|---|---|---|
| CDEP7767 | ctttcaggtagtgaatctgctttaggtatgtttgtacagag | Clone *ltaB*$^{H605A}$ onto pIA134 |
| CDEP7768 | ctctgtacaaacatacctaaagcagattcactacctgaaag | Clone *ltaB*$^{H605A}$ onto pIA134 |
| CDEP7869 | ggtatgtttgtacaggcaattttcctaaatg | Clone *ltaB*$^{R612A}$ onto pIA134 |
| CDEP7870 | catttaggaaaattgcctgtacaaacatacc | Clone *ltaB*$^{R612A}$ onto pIA134 |
| CDEP7771 | gatattacaaacatttggagcaaaaattcaaatgaatatg | Clone *ltaB*$^{R627A}$ onto pIA134 |
| CDEP7772 | catattcatttgaatttttgctccaaatgtttgtaatatc | Clone *ltaB*$^{R627A}$ onto pIA134 |
| CDEP7871 | catttggaagagcaattcaaatgaatatg | Clone *ltaB*$^{K628A}$ onto pIA134 |
| CDEP7872 | catattcatttgaattgctcttccaaatg | Clone *ltaB*$^{K628A}$ onto pIA134 |
| CDEP7956 | ggaagaaaaattcaagcaaatatgcaactagcacagac | Clone *ltaB*$^{M631A}$ onto pIA134 |
| CDEP7957 | gtctgtgctagttgcatatttgcttgaatttttcttcc | Clone *ltaB*$^{M631A}$ onto pIA134 |
| CDEP7873 | gaaaaattcaaatggcaatgcaactagc | Clone *ltaB*$^{N632A}$ onto pIA134 |
| CDEP7874 | gctagttgcattgccatttgaatttttc | Clone *ltaB*$^{N632A}$ onto pIA134 |
| CDEP7875 | ctcttttagttgcagattctggaatag | Clone *ltaB*$^{N691A}$ onto pIA134 |
| CDEP7876 | ctattccagaatctgcaactaaaagag | Clone *ltaB*$^{N691A}$ onto pIA134 |
| CDEP7877 | cttttagttaatgcatctggaatagttg | Clone *ltaB*$^{D692A}$ onto pIA134 |
| CDEP7878 | caactattccagatgcattaactaaaag | Clone *ltaB*$^{D692A}$ onto pIA134 |
| CDEP7958 | cttttagttaatgatgcaggaatagttgctgc | Clone *ltaB*$^{S693A}$ onto pIA134 |
| CDEP7959 | gcagcaactattcctgcatcattaactaaaag | Clone *ltaB*$^{S693A}$ onto pIA134 |
| CDEP7805 | tgaagtgagctaaggagtttaacatgaaaaaaaagattgtatcct | Clone *ltaB* (*RS04615*) from *P. bifermentans* ATCC 638 onto pMK03 |
| CDEP7806 | atcagccagccagccggtaccctactctttatcaaaaatcatcatatt | Clone *ltaB* (*RS04615*) from *P. bifermentans* ATCC 638 onto pMK03 |
| CDEP7807 | tgaagtgagctaaggagtttaacgtgaagaaaaaagcaac | Clone *ltaB* (*RS06385*) from *P. bifermentans* ATCC 638 onto pMK03 |
| CDEP7808 | atcagccagccagccggtacccgacatgagcgaagcgaat | Clone *ltaB* (*RS06385*) from *P. bifermentans* ATCC 638 onto pMK03 |
| CDEP7809 | tgaagtgagctaaggagtttaacgtgaattgtaaaataaagaaattatt | Clone *ltaB* (*RS06390*) from *P. bifermentans* ATCC 638 onto pMK03 |
| CDEP7810 | atcagccagccagccggtaccttatttttcttttaaaaccactttattaatt | Clone *ltaB* (*RS06390*) from *P. bifermentans* ATCC 638 onto pMK03 |
| CDEP7815 | tgaagtgagctaaggagtttaacgtgaagacaaaattcgtgt | Clone *ltaB* (*RS26125*) from *B. agri* DSM 6348 onto pMK03 |
| CDEP7816 | atcagccagccagccggtacctcacgtcgagcagcgctcc | Clone *ltaB* (*RS26125*) from *B. agri* DSM 6348 onto pMK03 |
| CDEP7813 | tgaagtgagctaaggagtttaacatgtttaagaaaagtttggc | Clone *ltaB* (*RS06015*) from *Peptostreptococcus* sp. Strain MV1 HM-1102 onto pMK03 |
| CDEP7814 | atcagccagccagccggtaccctaattaatatacttgtcttcttttga | Clone *ltaB* (*RS06015*) from *Peptostreptococcus* sp. Strain MV1 HM-1102 onto pMK03 |
| CDEP8104 | tgaagtgagctaaggagtttttgatgaaaaaaatggttgctg | Clone *ltaB* (*EDM54_08085*) from *B. borstelensis* NRS-818 onto pMK03 |
| CDEP8105 | atcagccagccagccggtactcaagagctggaatagcgtt | Clone *ltaB* (*EDM54_08085*) from *B. borstelensis* NRS-818 onto pMK03 |
| CDEP8106 | tgaagtgagctaaggagtttttgcttcagtggatacttatttt | Clone *ltaB* (*ABER04_18820*) from *B. laterosporus* ATCC 6456 onto pMK03 |
| CDEP8107 | atcagccagccagccggtacctaatgttttcgtgctacctt | Clone *ltaB* (*ABER04_18820*) from *B. laterosporus* ATCC 6456 onto pMK03 |
