## Supplementary material for "Identification of two novel lipoteichoic acid synthases essential for viability in *Clostridioides difficile*": Figure S1

Fig. S1

```
LtaB      MRKKIMSLFAIVIMIFVSTASYGFAE---EKGKVIFIDMNRTNLGSMMNIPILKNEIEKR   57
LtaA      MRKKIISFLATFIIILTSASQYSFADDMSTRGKVIFIDMNRTSMSNMLRIKSLREELDNR   60
          *****:*::* .*:*:.*::.*.**:    :***********.:..*:.*  *::*:::*

LtaB      GYVALMNIRGDQGTDDRRSYASMGAGTRASVTTEDYINFQKVDQDTAKAFEVATGQKAKE   117
LtaA      GYIGLMNIRGDKGSDDRRSYASMGAGGRANVANEEDINFESSSKDRNIVFESATGKSAKG   120
          **:.*******:*:************* **.*:.*: ***:. .:*   .** ***:.**

LtaB      INDLGINRSINENLEYGEYGANLGSLGKTLSENGLKTSVIGNADIIENNQLVKNRNLCLV   177
LtaA      INNLTINKSINENLNFGEYGSVLGSLGQSLSENGLKASVLGNSDIIENGQLIKNRNLCLT   180
          **:* **:******::****: *****::*******:**:**:*****.**:*******.

LtaB      AMDQYGRIDSGNIDNINKKDSKMPFGISTDYDKLLDETKKSYKNSDAIFVELGDTYRLDM   237
LtaA      AMDEYGRIPNGNVDTINKKDLSMPYGISTDYDKLIVETKEAYKNNDVVFVELGDTYRLDL   240
          ***:**** .**:*.***** .**:*********: ***::***.*.:***********:

LtaB      YRLNLNDNTYGTMKKNIVKNINKYLEQVFSMVGENDTVYIASAFPSDLDYKNKRRLSPII   297
LtaA      YKPNLNEKTYESVKDNIEKNIDVYLKNIFSMVEENDTVYIASAFPSDLDYKNKRRLSPII   300
          *: ***::** ::*.** ***: **:::**** ***************************

LtaB      KFSGEGKGLLESATTRKAGIVANLDVGVDILGTFGLENKEMVGRSYTLVNKEDNVGYLKS   357
LtaA      KLNGEGKGILSSSTTRREGIVTNLDVGVEILDNFNIKNQNMVGRKYELINRDDNKEFLMD   360
          *:.*****:*.*:***: ***:******:**..*.::*::****.* *:*::**  :* .

LtaB      LYERIVSISSIRSTIINGFVGVISASWVIAMFAIFFRNYMPH--KERVFKVLKEFIKLGI   415
LtaA      EYQKIVSISSIRSTILNGFVSIVFLSWIVAMIAILFRKHISKNYKETTFFILKELLKIGI   420
           *::***********:****.::  **::**:**:**::: :  ** .* :***::*:**

LtaB      IMPLSFMLAPLMNFKTPISISLGIFATTLILYLLGRKLFKDNDIKHMGFFATLTILVIVI   475
LtaA      VMPLSFMITPIMNFKTPLAISLGIIIITLTIYLISKVL-IKNDLKNMLFFTGLTIVIMVI   479
          :******::*:******::*****:  ** :**:.: *  .**:*:* **: ***:::**

LtaB      DSVFGTYLMQNNIMSYDAVIGARYYGVGNEYEGVTIASAVFGLAVLLHYKKVSKLLVVIF   535
LtaA      DAGFGSYLMKSNVMSYDCIIGARYYGVGNEYQGVAIGSAIFTFAILLTYKNIPKWSIIVF   539
          *: **:***:.*:****.:************:**:*.**:* :*:** **:: *  :::*

LtaB      SLIILITSAYPSMGANVGGAISECVAYLLFIMLIYDIKIDFKKAVLLAVSAVVVVFAFAA   595
LtaA      SLVILITSASPIMGANVGSAISECVAFLLFILLIYNVKIDFKKIVLLGIAVLFVLGVFVA   599
          **:****** * ******.*******:****:***::****** ***.::.:.*: .*.*

LtaB      LDILSGSESHLGMFVQRIFLNGPGEILQTFGRKIQMNMQLAQTSVWVNILLVGIGVIAVL   655
LtaA      IDMILGSNSHLGMFVKEIYFNGPGEIIQTFSRKIEMNLKLAQTSAWVNILLTGIGVILVL   659
          :*:: **:*******:.*::******:***.***:**::*****.******.***** **

LtaB      IFKPSRHFRKIMNNFPILFKGFIASMVGCIITLLVNDSGIVAASTASIYILIPLIIININ   715
LtaA      MINQIRYFKQLMDEYPIVFKGFIASIAGCLVTLLVNDSGIVSSATAFIYVIVPMITLSIN   719
          :::  *:*:::*:::**:*******:.**::**********:::** **:::*:* :.**

LtaB      MIIFEKNN  723
LtaA      LTALKE--  725
          :  :::
```

**Figure S1. LtaA and LtaB are closely related.** Overall, LtaA and LtaB share 63% amino acid identity and 82% similarity across their full-length sequences, consistent with partially overlapping functions in LTA synthesis. Shown is an alignment of LtaA and LtaB from *Clostridioides difficile* strain R20291 generated using Clustal Omega (74). Identical residues are indicated by asterisks (*), strongly similar residues by colons (:), and weakly similar residues by periods (.). Residues of LtaB subsequently selected for alanine-substitution analysis are in bold and highlighted based on their effects on viability and LTA synthesis: **Red** = severe viability defect and production of short LTA; **Orange** = partial viability defect and production of intermediate-length LTA; **Yellow** = no detectable viability defect and production of intermediate-length LTA; **Bolded but not highlighted** = no detectable effect on viability or LTA synthesis.
