## Supplementary material for "Identification of two novel lipoteichoic acid synthases essential for viability in *Clostridioides difficile*": Figure S2

Fig. S2

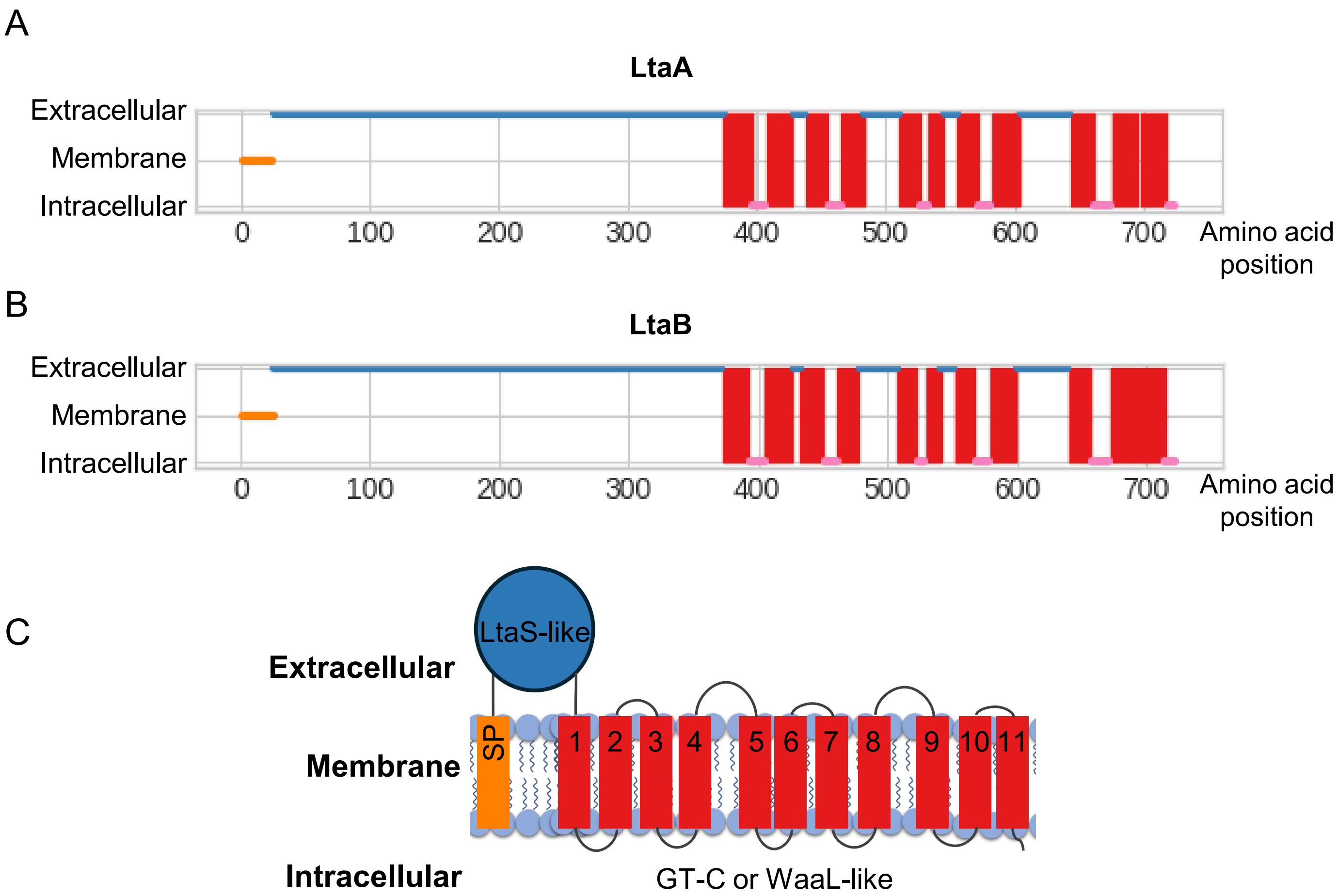


**Figure S2. Predicted topology of LtaA and LtaB.** LtaA and LtaB share a conserved membrane topology consistent with membership in the glycosyltransferase C superfamily. **(A,B)** Topology predictions for LtaA (A) and LtaB (B) generated using DeepTMHMM, showing an N-terminal cleavable signal peptide (orange), a large extracellular domain (blue), and a C-terminal region containing multiple predicted transmembrane helices (red). **(C)** Schematic representation of the predicted topology of LtaA and LtaB. Both proteins are predicted to contain an extracellular LtaS-like domain followed by a GTC-like domain with 11 transmembrane helices.
