## Supplementary material for "Identification of two novel lipoteichoic acid synthases essential for viability in *Clostridioides difficile*": Figure S3

Fig. S3

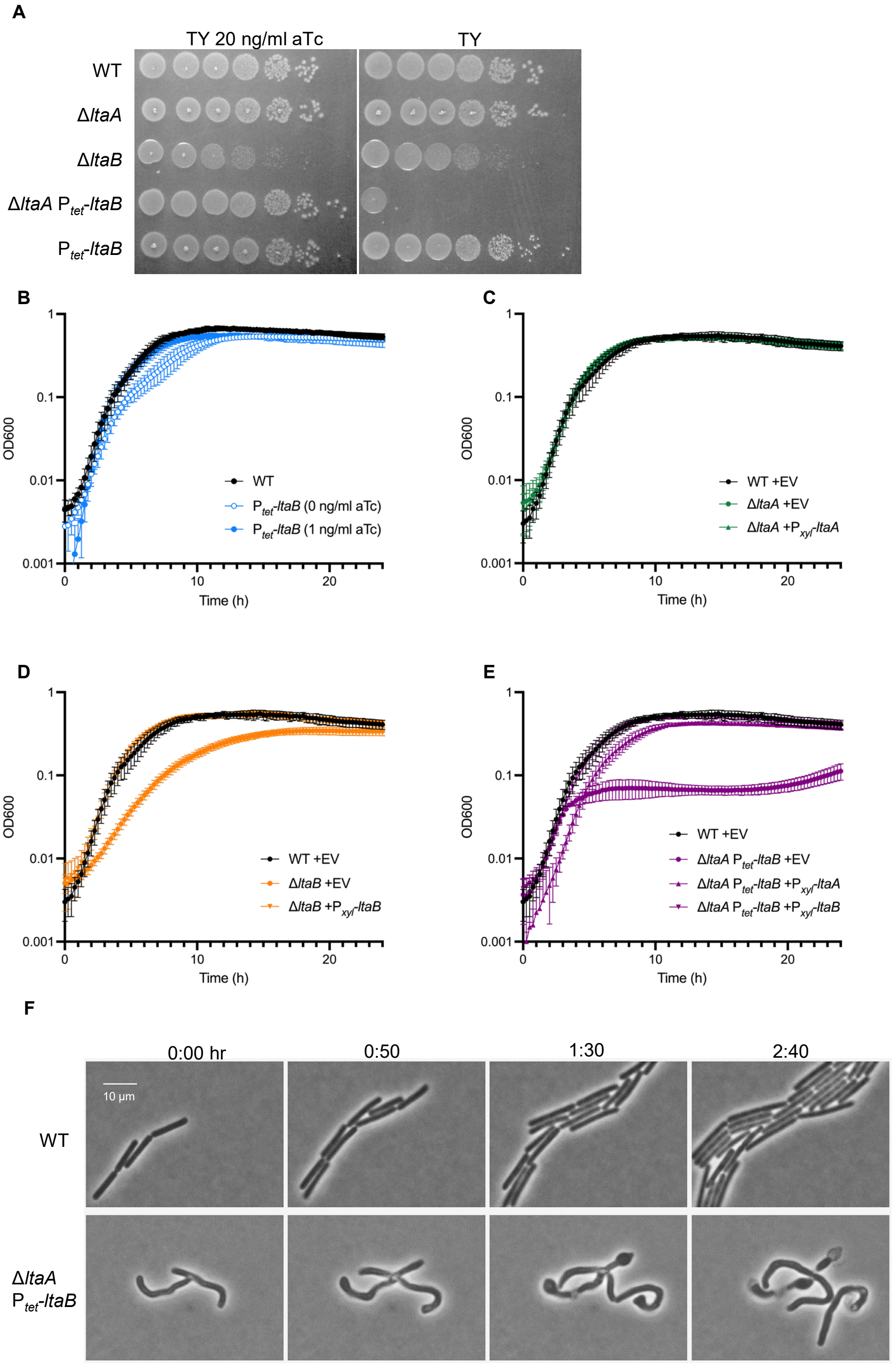


**Figure S3. Requirement for LtaA and LtaB in growth and morphology.** Loss or depletion of LtaB, but not LtaA, results in severe growth and morphological defects.
**(A)** Spot-titer viability assays of wild-type and mutant strains grown under the indicated inducing or depletion conditions. **(B–E)** Growth curves of the indicated strains monitored in liquid culture.
**(F)** Time-lapse phase-contrast microscopy of Δ*ltaA* $P_{tet}$-*ltaB* cells following depletion of LtaB, showing progressive loss of rod shape and cell lysis. Images are representative of three independent experiments. The growth curve is graphed as the mean ± sd of at least three experiments.
**Strains used: (A)** WT = R20291; Δ*ltaA* = CDE5015; Δ*ltaB* = EW169; $P_{tet}$-*ltaB* = CDE5011; Δ*ltaA* $P_{tet}$-*ltaB* = CDE5029. **(B)** WT = R20291; $P_{tet}$-*ltaB* = CDE5011. **(C)** Δ*ltaA* + EV = CDE5048; Δ*ltaA* + *ltaA* = CDE5050. **(D)** WT + EV = BZ153; Δ*ltaB* EV = EW211; Δ*ltaB* + *ltaA*= EW213. **(E)** Δ*ltaA* $P_{tet}$-*ltaB* + EV = CDE5031; Δ*ltaA* *$P_{tet}$-ltaB* + *ltaA* = CDE5033; Δ*ltaA* *$P_{tet}$-ltaB* + *ltaB* = CDE5032 (Table 3).
