## Supplementary material for "Identification of two novel lipoteichoic acid synthases essential for viability in *Clostridioides difficile*": Figure S4

### Fig. S4

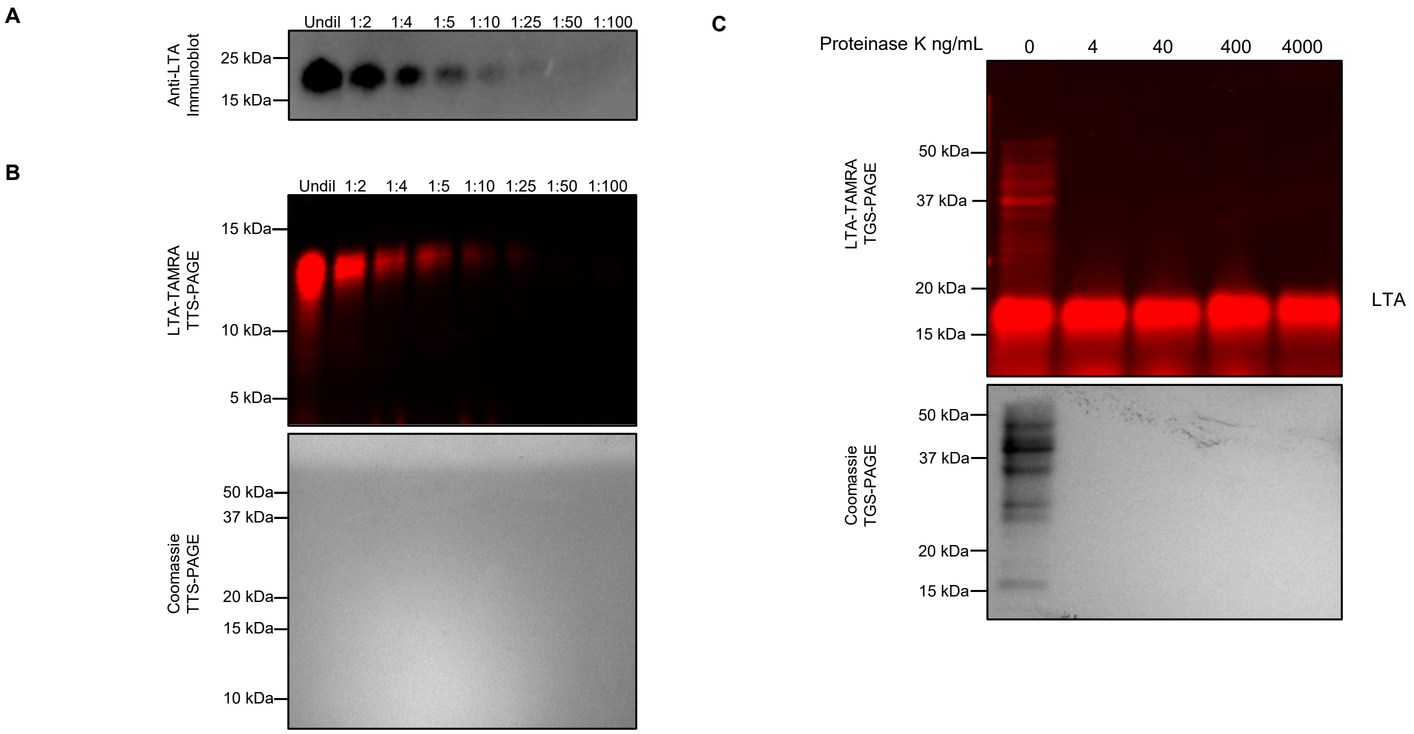


**Figure S4. LTA Detection Methods. (A)** Sensitivity of LTA detection by immuno-blotting. LTA extracts from wild-type were diluted serially as indicated above the blot and analyzed as described in Methods. **(B)** Sensitivity of LTA detection by metabolic labeling. LTA from wild-type cells grown in the presence of myristic acid alkyne was extracted with hot phenol as described in Methods. Extracts were reacted with TAMRA-azide by click chemistry and serial dilutions were analyzed by in-gel fluorescence. After imaging, the gel was stained with Coomassie Brilliant Blue, revealing the absence of proteins in the hot phenol extracts. **(C)** The fluorescent band that runs at 15-20 kDa is not a lipoprotein. Wild-type cells grown in the presence of myristic acid alkyne were lysed by sonication. Lysates were treated with increasing amounts of Proteinase K as indicated, reacted with TAMRA-azide by click chemistry and analyzed by in-gel fluorescence. After imaging, the gel was stained with Coomassie Brilliant Blue.

**Strains used:** WT = R20291
