## Supplementary material for "Identification of two novel lipoteichoic acid synthases essential for viability in *Clostridioides difficile*": Figure S5

Fig. S5

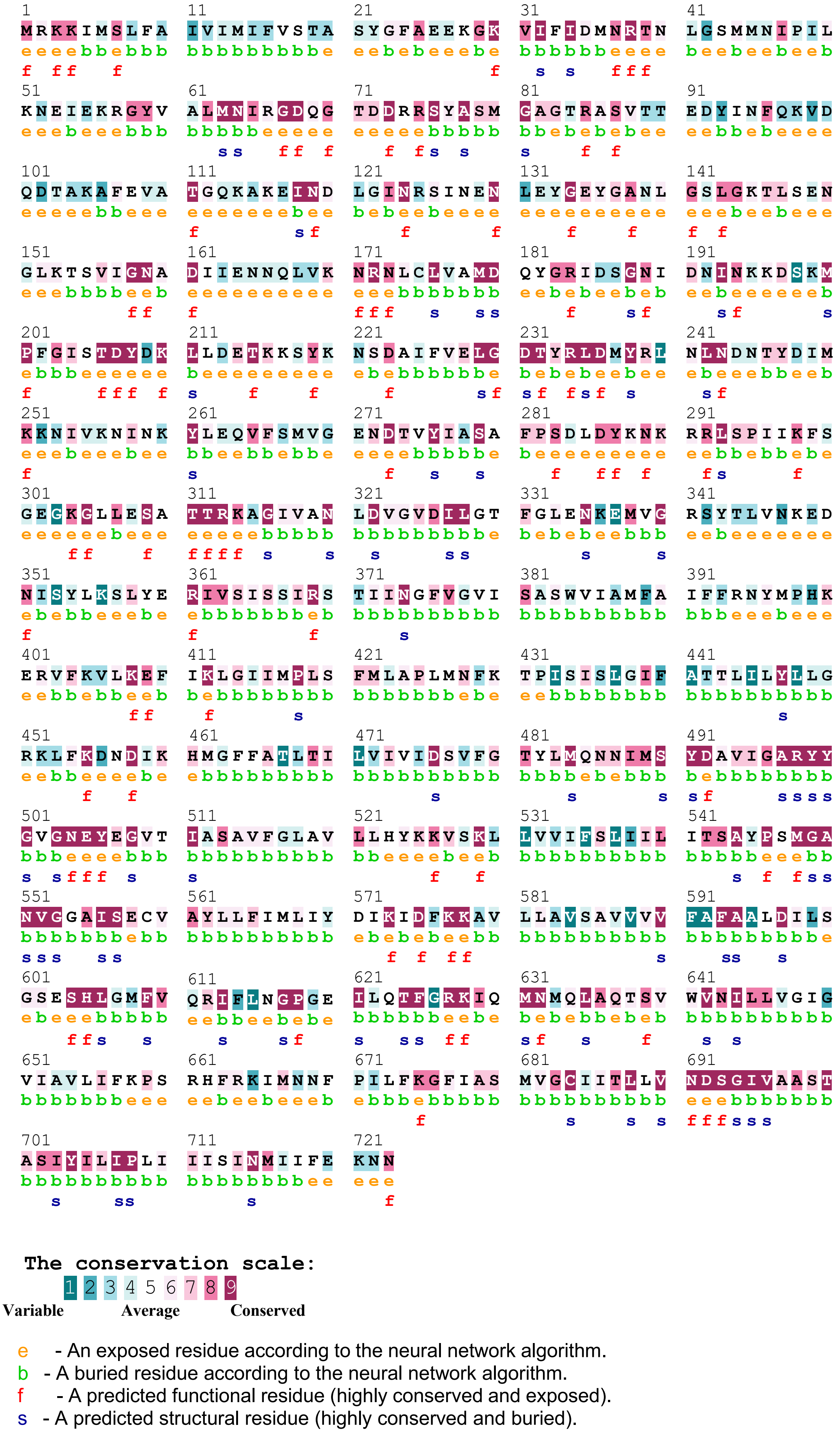


**Figure S5. Conservation of amino acids in LtaB.** ConSurf was used to map a multiple sequence alignment of 67 LtaB homologs from various *Peptostreptococcaceae* (see Table S2) onto the AlphaFold3 structural model of *C. difficile* LtaB. Highly conserved residues cluster within the predicted glycosyltransferase C–like transmembrane domain.
