## Supplementary material for "Identification of two novel lipoteichoic acid synthases essential for viability in *Clostridioides difficile*": Figure S6

### Fig. S6

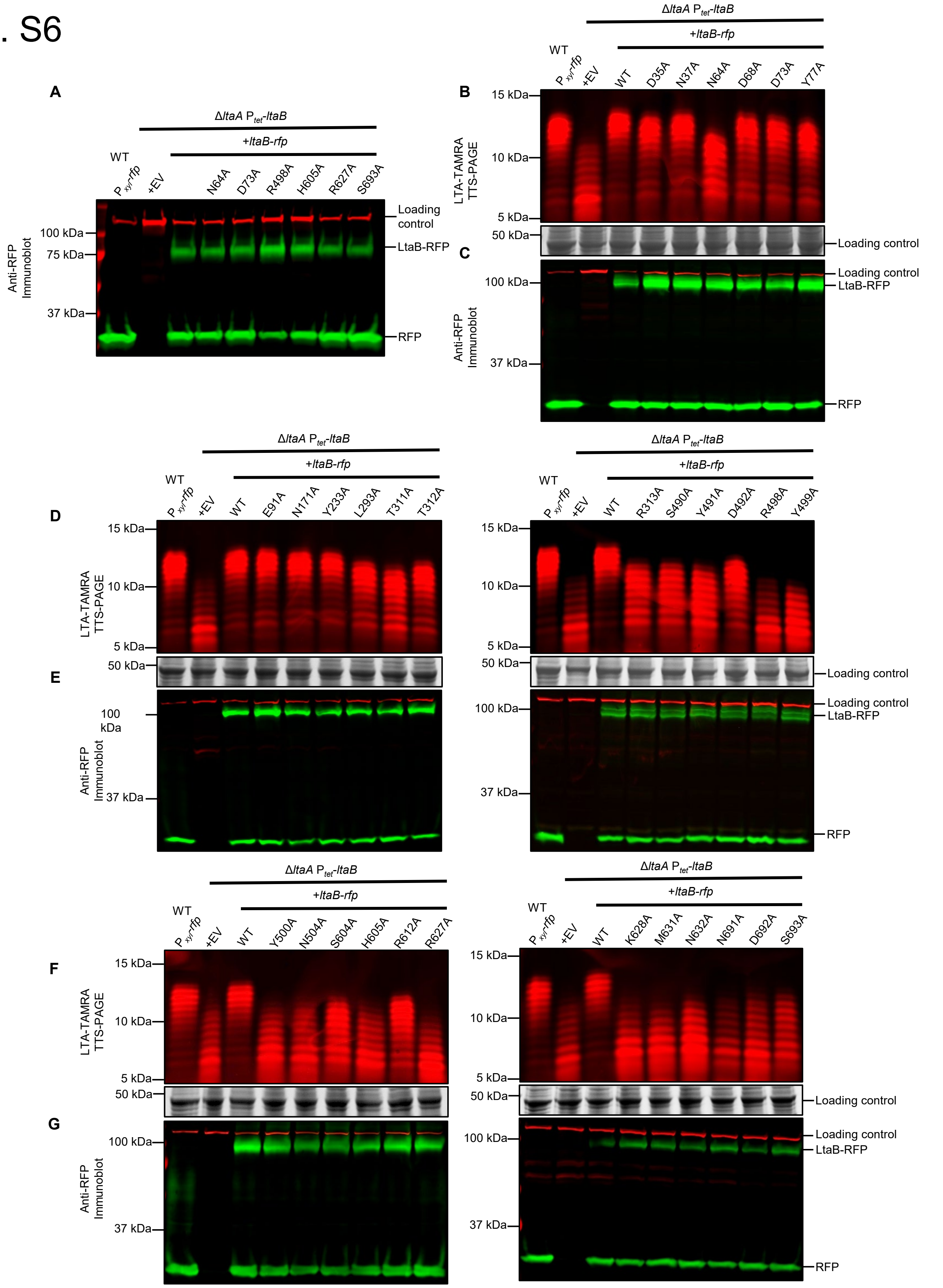


**Figure S6. Effect of alanine substitutions in the GTC-like domain of LtaB on LTA synthesis. (A,C,E,G)** Immunoblot detection of LtaB-RFP alanine-substitution mutants expressed in the Δ*ltaA P$_{tet}$-ltaB* background. **(B,D,F)** Metabolic labeling of LTA produced by the corresponding strains, resolved on Tris-Tricine gels. Mutations that impair viability are associated with the production of short LTA polymers. All images are representative of at least three independent experiments.

**Strains used:** WT + *rfp* = AP441; Δ*ltaA P$_{tet}$-ltaB* + EV = CDE5029; Δ*ltaA* P$_{tet}$*-ltaB* + *ltaB-rfp* = CDE5231; Δ*ltaA* P$_{tet}$*-ltaB* strains expressing individual LtaB-RFP alanine-substitution mutants = (see Table 3).
