## Supplementary material for "Identification of two novel lipoteichoic acid synthases essential for viability in *Clostridioides difficile*": Figure S7

Fig. S7

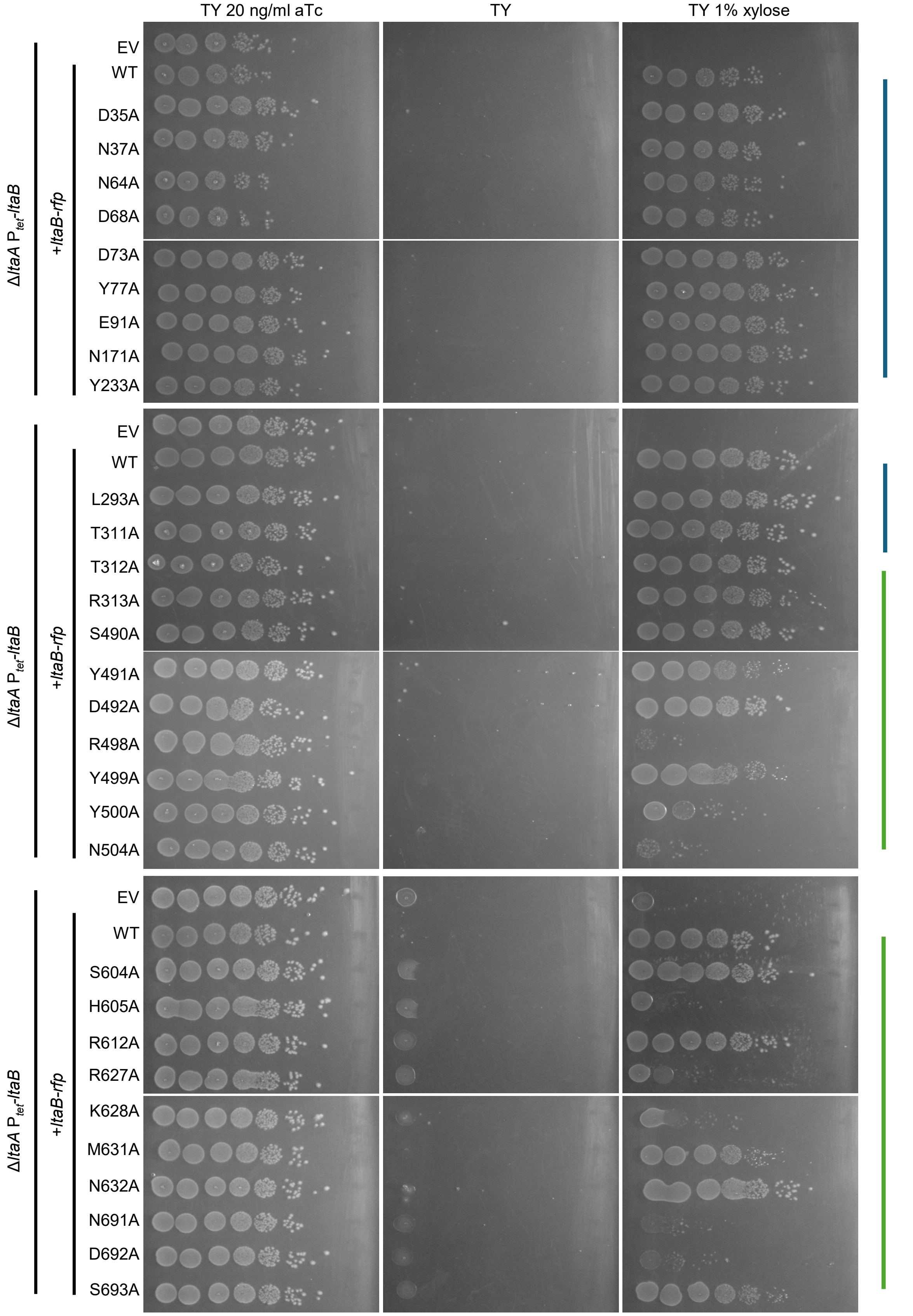


**Figure S7. Effect of LtaB alanine substitutions on viability.** Specific conserved residues in LtaB are required for cell viability. Spot-titer viability assays of Δ*ltaA* $P_{tet}$-*ltaB* strains expressing individual LtaB alanine-substitution mutants from a $P_{xyl}$ plasmid. Mutations within the conserved core of the GTC-like domain result in severe or complete loss of viability. Images are representative of at least three independent experiments.

**Strains used:** WT + *rfp* = AP441; Δ*ltaA* $P_{tet}$-*ltaB* + EV = CDE5029; Δ*ltaA* $P_{tet}$-*ltaB* + *ltaB-rfp* = CDE5231; Δ*ltaA* $P_{tet}$-*ltaB* strains expressing individual LtaB-RFP alanine-substitution mutants = (see Table 3).
