## Supplementary material for "Identification of two novel lipoteichoic acid synthases essential for viability in *Clostridioides difficile*": Figure S8

Fig. S8

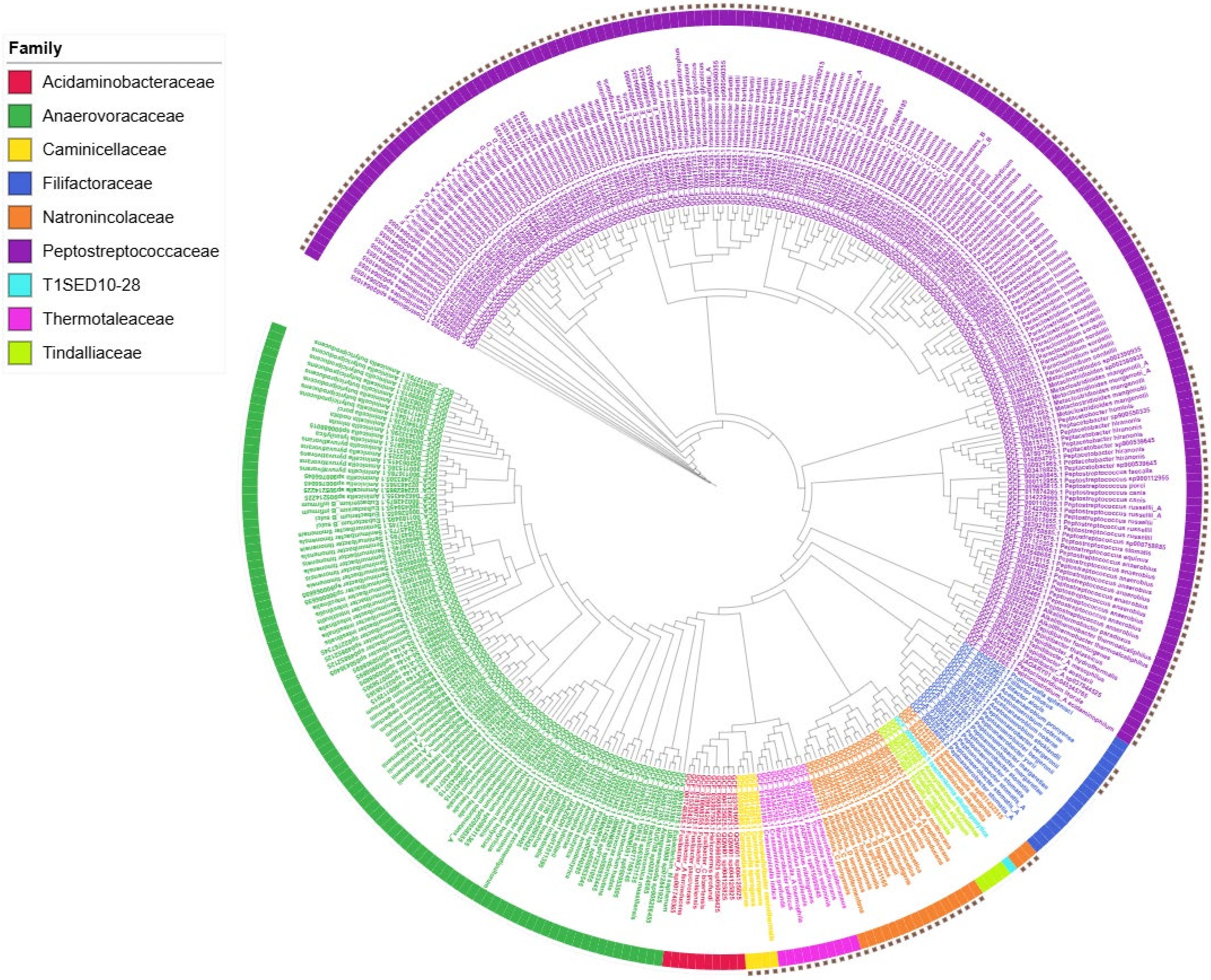


**Figure S8. Phylogenetic distribution of LtaB homologs among families within the order *Peptostreptococcales* and closely related lineages.** Tips and inner ring are colored according to family designation as indicated in the key. Filled-in squares in the outer ring denote the presence of an LtaB homolog identified using MMseqs2 searches of the Fast.genomics database.
